# Dynamic Causal Tractography Analysis of Auditory Descriptive Naming: An Intracranial Study of 106 Patients

**DOI:** 10.1101/2025.03.07.641428

**Authors:** Aya Kanno, Ryuzaburo Kochi, Kazuki Sakakura, Yu Kitazawa, Hiroshi Uda, Riyo Ueda, Masaki Sonoda, Min-Hee Lee, Jeong-Won Jeong, Robert Rothermel, Aimee F. Luat, Eishi Asano

## Abstract

Humans understand and respond to spoken questions through coordinated activity across distributed cortical networks. However, the causal roles of connectivity engagements alternating across multiple white matter bundles remain understudied at the whole-brain scale. Using intracranial high-gamma activity recorded from 7,792 non-epileptic electrode sites in 106 epilepsy patients who underwent direct cortical stimulation mapping, we constructed an atlas visualizing the millisecond-scale dynamics of functional connectivity during a naming task in response to auditory questions. This atlas, the Dynamic Causal Tractography Atlas, identified functional connectivity patterns at specific time windows most strongly associated with stimulation-induced language- and speech-related manifestations (p-value range: 2.5 × 10^-5^ to 6.6 × 10^-14^; rho range: +0.54 to +0.82). The atlas revealed that no single intra-hemispheric fasciculus was consistently engaged in all naming stages; instead, each fasciculus supported specific stages, with multiple distinct major fasciculi simultaneously contributing to each stage. Additionally, this atlas identified the specific linguistic stages and fasciculi where handedness effects became evident. Our findings clarify the dynamics and causal roles of alternating, coordinated neural activity through specific fasciculi during auditory descriptive naming, advancing current neurobiological models of speech network organization. Additionally, we have made our white matter streamline template and intracranial EEG data available as open-source material, enabling investigators to construct personalized dynamic tractography atlases.

## Introduction

To understand and answer spoken questions, the human brain transforms perceived sounds into meaning, mentally retrieves words relevant to the questions, and converts these words into overt articulation. More specifically, auditory naming-related sensorimotor and cognitive functions include the perception and short-term storage of spoken sounds, semantic-syntactic analyses, lexical retrieval, and response initiation and overt articulation (Gathercole et al., 1994; Wise et al., 1999; Hickok and Poeppel, 2000; Liberman and Whalen, 2000; Baddeley, 2003; Liebenthal et al., 2005; Skeide and Friederici, 2016). Current neurobiological models indicate that humans perform this series of functions through coordinated activity via white matter bundles, referred to as fasciculi, predominantly across the left-hemispheric cortical regions (Catani et al., 2005; Dick and Tremblay, 2012; Chang et al., 2015; Gajardo-Vidal et al., 2021). Among these, the left arcuate fasciculus, which dorsally connects the temporal and frontal lobes, is a critical structure for auditory descriptive naming as lesions lead to impairments in semantic-syntactic processing and lexical retrieval (Bernal and Ardila, 2009; Marchina et al., 2011; Wilson et al., 2011; Fridriksson et al., 2013; Griffiths et al., 2013; Gajardo-Vidal et al., 2021; Giampiccolo and Duffau, 2022). Major white matter bundles suggested to support speech processes also include the left superior longitudinal fasciculus (SLF), inferior longitudinal fasciculus (ILF), inferior fronto-occipital fasciculus (IFOF), frontal aslant fasciculus, and uncinate fasciculus (Catani et al., 2005; Mandonnet et al., 2007; Dick and Tremblay, 2012; Friederici and Gierhan, 2013; Chang et al., 2015; Lambon Ralph et al., 2017).

Although lesion, electrical stimulation, and fMRI studies have provided insights into the involvement of specific fasciculi in speech, the dynamics of neural coordination through these fasciculi have been understudied. Investigators suggest that intracranial EEG recording in patients with focal epilepsy can provide a unique opportunity to assess the dynamics of cortical modulations during overt speech, as well as the neural coordination across cortical regions (Flinker et al., 2015; Anumanchipalli et al., 2019; Woolnough et al., 2021). The advantage of intracranial EEG includes its signal fidelity, which is more than 100 times better than that of scalp EEG and magnetoencephalography (Ball et al., 2009), both of which are inevitably affected by electromyographic artifacts from ocular and temporal muscles during saccadic eye movements and overt articulation (Yuval-Greenberg et al., 2008; Carl et al., 2012). Our previous intracranial EEG study of 13 English-speaking patients with focal epilepsy developed a methodology to assess functional connectivity between cortical regions during overt naming (Kitazawa et al., 2023). In that study, we defined the enhancement of functional connectivity as significant, simultaneous, and sustained augmentation of high-gamma amplitude (70-110 Hz) directly connected by a white matter streamline on MRI tractography. However, due to the small sample size, our previous iEEG study could only assess and visualize functional connectivity across limited cortical regions and failed to detect task-related diminutions in functional connectivity. Furthermore, the study was not designed to assess the causal significance of task-related connectivity enhancement, as it did not correlate the observed connectivity modulations with symptoms elicited by direct electrical stimulation (Kitazawa et al., 2023).

The current study aimed to generate a dynamic causal tractography atlas that visualizes auditory naming-related modulations of functional connectivity through specific white matter on a millisecond scale, while assessing the causal role of the observed enhancement in functional connectivity by correlating it with stimulation-induced manifestations, including auditory hallucinations, receptive and expressive aphasia, speech arrest, and facial sensorimotor symptoms (Hamberger, 2007; Tate et al., 2014; Nakai et al., 2017; Lu et al., 2021; Woolnough et al., 2021). We predicted that cortical sites exhibiting enhanced functional connectivity during hearing (but not after auditory stimulus offset) would correlate most strongly with stimulation-induced auditory hallucinations, as acoustic processing is expected to begin early in the listening phase and precede semantic analysis (Hickok and Poeppel, 2000; Skeide and Friederici, 2016). Additionally, we predicted that functional connectivity enhancement around or after the offset of an auditory question would correlate most strongly with stimulation-induced receptive aphasia. Moreover, we expected this correlation to emerge earlier for receptive aphasia than for expressive aphasia, because listeners must first process the phonemes and comprehend the question before retrieving or generating an appropriate answer (Levelt, 1999; Hickok and Poeppel, 2000; Skeide and Friederici, 2016). Finally, we predicted that the tightest correlation of connectivity enhancement with stimulation-induced speech arrest and sensorimotor symptoms would occur around overt response (Chang et al., 2013; Anumanchipalli et al., 2019; Lu et al., 2021). We expected that this analysis would reveal the network organization supporting each stage of language- and speech-related processes necessary for auditory descriptive naming.

The current study aimed to determine the dynamics and roles of 12 major fasciculi in auditory descriptive naming while also assessing local u-fibers not classified as major fasciculi. We predicted that functional connectivity enhancement in the left arcuate fasciculus would be the earliest, most intense, and most sustained among all the fasciculi mentioned, due to its involvement in the short-term storage of spoken words, semantic-syntactic analysis, and lexical retrieval (Bernal and Ardila, 2009; Marchina et al., 2011; Griffiths et al., 2013; Gajardo-Vidal et al., 2021; Giampiccolo and Duffau, 2022; Janssen et al., 2023; Zhao et al., 2023). Because of its suggested role in semantic analysis or lexical retrieval (Mandonnet et al., 2007; Almairac et al., 2015; Harvey and Schnur, 2015), functional connectivity enhancement in the left SLF, ILF, and IFOF would be most prominent between question offset and response onset. Given its involvement in speech initiation (Chernoff et al., 2018; Zhong et al., 2022), functional connectivity enhancement in the left frontal aslant fasciculus would be most prominent immediately prior to overt responses. Given the lack of significant symptoms elicited by stimulation or resection (Duffau et al., 2009), the functional connectivity enhancement in the left uncinate fasciculus would be modest.

We aimed to enhance the generalizability of our atlas by assessing the impacts of patient age and handedness—independent of epilepsy-related profiles—on task-related neural dynamics. We hypothesized that older age and right-handedness would be associated with greater left-hemispheric dominant functional connectivity enhancement during semantic analysis and lexical retrieval periods. Conversely, we hypothesized that younger and left-handed individuals would exhibit more symmetric functional connectivity patterns during these periods. This hypothesis is based on the observation that younger individuals, compared to older ones, display more symmetric language task-related hemodynamic activation on fMRI (Szaflarski et al., 2006). and show better language function recovery following extensive left-hemispheric damage (Boatman et al., 1999). Additionally, fMRI studies have reported that left-handed healthy individuals, compared to right-handed ones, less frequently exhibit left-hemispheric dominant hemodynamic activation during language tasks (Szaflarski et al., 2002; Mazoyer et al., 2014).

## Materials and methods

### Participants

The inclusion criteria of this observational study were native English speakers with focal seizures who performed an auditory descriptive naming task during extraoperative iEEG recording at Children’s Hospital of Michigan or Harper University Hospital in Detroit between January 2007 and December 2023. The exclusion criteria were: (i) inability to complete the task, (ii) significant brain malformations affecting the central or lateral sulcus (for example, megalencephaly and perisylvian polymicrogyria), (iii) history of prior resective epilepsy surgery, and (iv) evidence of right-hemispheric language dominance, as indicated by the Wada test results or left-handedness in conjunction with left-hemispheric congenital neocortical lesions. Our eligibility criteria ensured that all study participants had essential language functions in the left hemisphere. We have previously discussed the validity of inferring right-hemispheric language dominance based on left-handedness and left-hemispheric congenital neocortical lesions (Möddel et al., 2009; Sonoda et al., 2022). The Institutional Review Board of Wayne State University approved the current study, and we obtained written informed consent from the patients’ legal guardians and written assent from pediatric patients aged 13 years or older.

### Intracranial electrode placement and localization within standard brain template

Patients underwent the placement of platinum grid and strip disk electrodes and depth electrodes (10 mm center-to-center) in the intracranial space to localize the boundaries between the epileptogenic zone and eloquent areas. We created a three-dimensional cortical surface image and determined each electrode’s location using preoperative T1 spoiled gradient-recalled MRI, post-implant CT, and the FreeSurfer software package (http://surfer.nmr.mgh.harvard.edu) (Stolk et al., 2018; Sakakura et al., 2023). For group-level analysis, we spatially normalized all electrode locations to the FreeSurfer standard brain space (Desikan et al., 2006; Ghosh et al., 2010; Sakakura et al., 2023). The FreeSurfer script automatically parcellated the cortical gyri and assigned each electrode site a corresponding region of interest (ROI) as defined in the Desikan-Killiany atlas (**Figure 1**) (Desikan et al., 2006).

**Figure 1.**
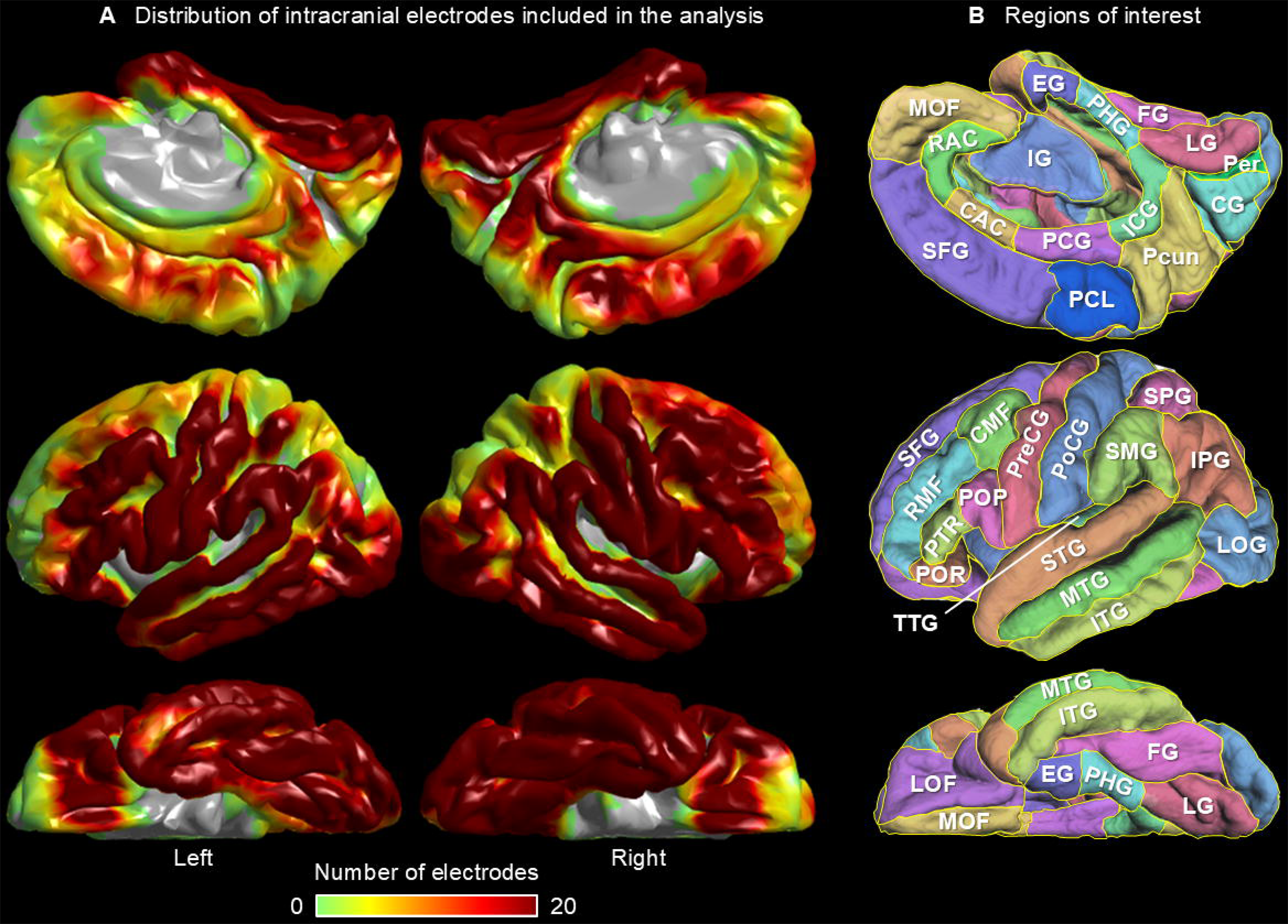
Spatial distribution of intracranial electrode sampling. **(A)** The figure shows the number of electrodes whose artifact-free, nonepileptic intracranial EEG data were available for measurement of task-related cortical high-gamma dynamics. **(B)** Regions of interest (ROIs), as defined in the Desikan-Killiany atlas (Desikan et al., 2006), are presented. In **Supplementary Table 2**, the full names corresponding to the given abbreviations of the ROIs are provided. To generate these images, we used FreeSurfer software (https://surfer.nmr.mgh.harvard.edu/fswiki/CorticalParcellation).

### Intracranial EEG (iEEG) data acquisition

Following intracranial electrode placement, patients were transferred to the epilepsy monitoring unit, where iEEG was continuously monitored with simultaneous video recording for 3-7 days using the Nihon Kohden Neurofax Digital System (Nihon Kohden America Inc, Foothill Ranch, CA, USA). We acquired iEEG signals at a sampling rate of 1,000 Hz and an amplifier bandpass filter of 0.016–300 Hz. The seizure onset zone (SOZ), responsible for generating habitual seizures (Asano et al., 2009), and regions generating interictal spike discharges (Kural et al., 2020) were clinically identified in a prospective manner by a clinical team that included board-certified clinical neurophysiologists and epileptologists. We evaluated the integrity of iEEG signals during the auditory naming task and identified signals affected by electromyographic artifacts (Nagasawa et al., 2011; Uematsu et al., 2013). For further iEEG analysis, we included only artifact-free, nonepileptic electrode sites. Nonepileptic electrode sites were defined as those unaffected by the SOZ, interictal spike discharges, or MRI-visible structural lesions (Frauscher et al., 2018; Taylor et al., 2022; Sakakura et al., 2023). Ictal propagation was not used to define nonepileptic electrode sites, as no universally accepted criterion exists for identifying electrodes involved in early propagation. Similarly, the presence or absence of afterdischarges induced by electrical stimulation was not considered when determining whether an electrode site was nonepileptic. Our analytic approach aimed to minimize the effect of epileptiform discharges on task-related high-gamma amplitude measurements (Mercier et al., 2022). Out of 13,648 implanted electrodes, 7,792 were artifact-free, nonepileptic sites. All these 7,792 sites were included for the measurement and visualization of task-related cortical high-gamma dynamics (**Figure 1A**). **Supplementary Table 2** lists the exact number of artifact-free, nonepileptic electrode sites and contributing patients within each ROI. As detailed below, we performed ROI-based iEEG analysis in 54 regions (i.e., 27 in each hemisphere; **Figure 1B**), each of which contained at least five artifact-free, nonepileptic electrode sites derived from at least three patients.

### Auditory descriptive naming task

We assigned an auditory naming task to patients to localize regions involved in language and speech based on the time-frequency analysis described below. Patients were awake and comfortably seated on a bed in a room with minimized unwanted noises, and none had a seizure within two hours prior to the task. A series of questions were delivered via a speaker, and we instructed patients to overtly provide an answer (e.g., ‘Bird’ or ‘Plane’) for each question (e.g., ‘What flies in the sky?’). Each question began with either ‘what’, ‘where’, ‘when’, or ‘who’ and was designed to elicit one- or two-word answers with nouns (Kambara et al., 2018). This naming task is patient-friendly because no complex task instructions are necessary for children to complete (Nakai et al., 2017). Up to 100 trials were given, with the duration of each question ranging from 1.2 to 2.4 seconds (median: 1.8 seconds). Once a patient was confirmed to have verbalized an answer, the examiner delivered the next question 2 or 2.5 seconds after the button press. We marked stimulus onset, stimulus offset, and response onset on iEEG signals using sound waves recorded with hands-free microphones. We defined the response time as the duration between stimulus offset and response onset. Trials were excluded from further iEEG analysis if a patient failed to provide a relevant answer.

### Time-frequency analysis

We quantified task-related modulations of high-gamma amplitude, an outstanding summary measure of neural activation due to its strong correlation with increased neural firing rates (Ray et al., 2008; Leszczyński et al., 2020), hemodynamic responses (Scheeringa et al., 2011), and glucose metabolism (Nishida et al., 2008). A meta-analysis of 15 iEEG studies reported that cortical sites showing language task-related high-gamma augmentation were more likely to be classified as part of the language area defined by electrical stimulation mapping (Arya et al., 2018). Our prior iEEG study of 65 patients found that naming task-related augmentation of high-gamma activity at 70-110 Hz, among many spectral frequency bands including slow oscillations, was the best predictor of postoperative language outcomes (Sonoda et al., 2022).

We transformed iEEG signals into 10-ms/5-Hz time–frequency bins using the complex demodulation method incorporated in the BESA EEG Software (BESA GmbH, Gräfelfing, Germany) (Papp and Ktonas, 1977; Hoechstetter et al., 2004). This process involved multiplying the time-domain iEEG signal with a complex exponential, followed by a band-pass filter. Because a Gaussian-shaped low-pass finite impulse response filter was employed, this complex demodulation method is equivalent to a Gabor transformation. The time–frequency resolution for high-gamma measurement was ±15.8 ms and ±7.1 Hz (defined as the 50% power drop of the finite impulse response filter). We aligned the iEEG traces to stimulus onset, stimulus offset, and response onset to determine the percent change in high-gamma amplitude at 70-110 Hz compared to the average value during the baseline period, defined as the 400-ms interval between 200 and 600 ms prior to stimulus onset. At all artifact-free, nonepileptic electrode sites, we quantified the percent change in high-gamma amplitude during the following periods: between -200 and +500 ms relative to stimulus onset; between -500 and +500 ms relative to stimulus offset; between -500 and +500 ms relative to response onset. Here, ‘amplitude’ is a measure proportional to the square root of ‘power’. We then visualized the percentage change in high-gamma amplitude on a FreeSurfer standard brain template, using interpolation within a 10 mm radius from the electrode center (Sakakura et al., 2023). We used MATLAB R2023 software for visualization purposes (MathWorks, Natick, MA, USA).

In each ROI, we declared a significant augmentation of high-gamma amplitude when the lower limit of the 99.99% confidence interval (CI) of the mean across electrode sites exceeded zero for a duration of 50 ms or more. Similarly, we declared a significant attenuation when the upper limit of the 99.99% CI consistently fell below zero for at least 50 ms. Using a 99.99% CI corresponds to a Bonferroni correction for 500 repeated comparisons, and a 50-ms interval would include a minimum of three high-gamma oscillatory cycles. Although this stringent significance criterion increases the likelihood of a Type II error, it reduces the risk of a Type I error.

### Visualization of white matter streamlines on tractography

Our aims included generating new white matter streamline templates and providing these templates as open-source material. iEEG investigators commonly use standard brain templates, such as the Montreal Neurological Institute (MNI) standard space image (Jenkinson et al., 2012) and the Desikan-Killiany atlas (Desikan et al., 2006), to visualize neural dynamics at a group level. The iEEG research guidelines do not mandate investigators to create an anatomical brain template derived from study participants (Mercier et al., 2022). Creating an average brain template involves labor-intensive and time-consuming processes, such as manually inspecting and correcting artifactual signals and topological defects on anatomical MRI. Additionally, standard anatomical brain templates are generally assumed to represent the average brain anatomy of patients with reasonable spatial accuracy. Our previous iEEG-tractography study of 37 children with focal epilepsy, aged 5 to 20 years, suggests that the anatomical courses of white matter pathways are quite similar between patients and healthy individuals (Sonoda et al., 2021). We found that the lengths of white matter streamlines in individual patients’ tractography were highly correlated (Pearson correlation coefficient of 0.8) with those measured in 1,065 healthy individuals participating in the Human Connectome Project (HCP). Thus, in the present study, we used this HCP tractography dataset to generate white matter streamline templates (http://brain.labsolver.org/diffusion-mri-templates/hcp-842-hcp-1021) (Yeh et al., 2018). Using the DSI Studio script (http://dsi-studio.labsolver.org/) within the MNI standard space, we constructed, validated, and visualized the streamline connecting each pair of cortical ROIs with the shortest length among those detected using the specified parameters. We have provided the details of the fiber tracking and visual validation procedures in **Supplementary Figures 1 and 2.** Board-certified neurosurgeons (A.K. and R.K.) classified each of the validated white matter streamlines into one of the following 13 categories: (i) arcuate fasciculus, (ii) SLF, (iii) middle longitudinal fasciculus (MLF), (iv) ILF, (v) IFOF, (vi) frontal aslant fasciculus, (vii) parietal aslant fasciculus, (viii) uncinate fasciculus, (ix) cingulum fasciculus, (x) extreme capsule, (xi) intra-hemispheric u-fibers (i.e., other than those listed above), (xii) callosal fibers, or (xiii) anterior commissural fibers. **Supplementary Figures 2 and 3** provide the criteria used to define these fasciculi and visualizes the spatial characteristics of white matter streamlines classified into specific categories.

### Definition and visualization of task-related functional connectivity modulations

We determined the time bins and white matter pathways showing significant task-related modulations of functional connectivity (Kitazawa et al., 2023; Ono et al., 2023; Ueda et al., 2024). We defined functional connectivity between a given pair of ROIs as either significantly enhanced or diminished based on the following two criteria: (i) both ROIs must exhibit significant, simultaneous, and sustained modulations in high-gamma amplitude for five or more consecutive 10-ms time bins, and (ii) these ROIs must be directly connected by a streamline on diffusion-weighted imaging (DWI) tractography. We then visualized the white matter streamlines showing significant functional connectivity enhancement and diminution, updating every 10 ms.

To ensure the statistical validity of significant functional connectivity enhancements, we assessed the likelihood of random co-occurrence of high-gamma augmentation (i.e., Type I error) by calculating the chance probability of high-gamma co-augmentation lasting 50 ms or longer, given that significant high-gamma augmentation occurred in χ% of the 2,500-ms analysis period on average across 54 ROIs. Given the 1431 distinct pairs of ROIs, the chance probability of simultaneous occurrence of significant high-gamma augmentation at least at two ROIs for 50 ms or longer was estimated as follows: 1 – [1 – ((χ/100)^2^)^5^]^1431^ (Ueda et al., 2024).

We visualized the temporal profiles and intensity of functional connectivity enhancement and diminution across 10 intra-hemispheric fasciculi, local u-fibers, and two inter-hemispheric white matter pathways mentioned above. We calculated the onset (ms), duration (ms) and maximum intensity (%) of functional connectivity enhancement for each white matter pathway. The intensity of functional connectivity enhancement between ROI_1_ and ROI_2_ at a given time bin was defined as the square root of [high-gamma amplitude % change at ROI_1_ × high-gamma amplitude % change at ROI_2_]. If no streamline directly connected ROI_1_ and ROI_2_, the intensity of functional connectivity enhancement between these regions was considered zero.

### Electrical stimulation mapping

As part of the routine presurgical evaluation, we performed electrical stimulation mapping (Nakai et al., 2017). Before initiating electrical stimulation mapping, the neuropsychologist conducted a practice session to confirm that the patient could readily answer a given auditory question during the baseline period. Questions that could not be answered during this period were excluded from electrical stimulation mapping. We delivered a pulse train of repetitive electrical stimuli to adjacent electrode pairs, with a stimulus frequency of 50 Hz, a pulse duration of 0.3 ms, and a train duration of up to 5 seconds. The stimulus intensity was initially set at 3 mA and increased stepwise up to 9 mA until investigators observed a clinical response or after-discharge. During each trial, patients were asked to answer auditory questions beginning with ‘what’, ‘when’, ‘where’, or ‘who’. Electrode sites where stimulation reproducibly elicited clinical symptoms without inducing afterdischarge were prospectively identified and assessed at bedside by at least two investigators, including a neuropsychologist (R.R.), both blinded to task-related high-gamma dynamics. If a patient failed to verbalize an answer during stimulation, we inquired about the reason. Additional tasks, such as humming, syllable repetition, counting, and reciting the alphabet, were performed to determine the specific causal role of each cortical site where stimulation elicited a clinical symptom. We calculated the probability of each manifestation elicited by stimulation at each electrode site within each patient. By averaging across all available patients, we presented the mean probability of each stimulation-induced symptom on the FreeSurfer standard brain template. We included the following as language- or speech-related symptoms: (i) Auditory hallucination, defined as the perception of various pitches of sounds or alterations in the neuropsychologist’s voice pitch; (ii) Receptive aphasia, defined as the inability to understand a question despite being aware that the neuropsychologist had asked one; (iii) Expressive aphasia, defined as dysnomia or the inability to provide a relevant answer during a 5-second stimulation, even though the neuropsychologist confirmed that the patient understood the question and could vocalize; (iv) Speech arrest, defined as the inability to initiate or continue vocalization, not attributable to forced jerking of the tongue or lips; and (v) Face sensorimotor symptoms, defined as involuntary movement or sensation in the facial area, with or without overt dysarthria. Stimulation-induced paraphasic and nonsensical speech responses were classified as expressive aphasia if the patient could repeat the question immediately after the stimulation trial and as receptive aphasia if the patient failed to do so. Hesitation was disregarded if the patient answered before the offset of electrical stimulation but classified as expressive aphasia if the patient answered only after the offset and the symptom could not be attributed to speech arrest.

### Assessment of patient demographics and epilepsy characteristics

We employed a mixed model analysis to determine the impact of patient demographics and epilepsy-related profiles on response time. We included the following variables in the mixed model analysis: (i) square root of patient age (√years), (ii) handedness (1 if left-handed; 0 otherwise), (iii) left-hemispheric epileptogenicity (1 if surgical intervention or SOZ involved the left hemisphere; 0 otherwise), (iv) an MRI-visible lesion (1 if present; 0 if absent), and (v) the number of antiseizure medications taken immediately prior to the iEEG recording. We incorporated the square root of age in the mixed model, assuming that the developmental changes in naming behaviors and neural systems would be greater in younger individuals. We interpreted the need for multiple antiseizure medications as indicative of greater seizure-related cognitive burdens (Kwan and Brodie, 2001; Kuroda et al., 2021). The median response time was treated as the dependent variable, while patient was included as a random factor, allowing the intercept to vary across patients. We used SPSS Statistics 28 (IBM Corp., Chicago, IL, USA) for statistical analyses. All reported p-values are two-sided, and we considered a p-value of <0.05 significant.

Likewise, we employed a mixed model analysis to determine the impact of aforementioned patient demographics, epilepsy-related profiles, and response time on high-gamma amplitude (% change compared to baseline) at each ROI. The dependent variable was the median at each electrode site during each 100-ms epoch within the 2,500-ms period between stimulus onset and 500-ms after the response onset. Given the repeated analyses across 54 ROIs and 25 100-ms epochs, we considered a Bonferroni-corrected p-value of <0.05 as significant.

### Statistical assessment of the relationship between functional connectivity and stimulation-induced manifestations

We assessed the strength of the association between the intensity of functional connectivity enhancement during specific time windows and language- or speech-related functions identified through electrical stimulation mapping. To do this, for each ROI in 10-ms time bins, we calculated the intensity of functional connectivity enhancement, averaged across all pairs. We used Spearman’s rank test to assess the relationship between the mean functional connectivity enhancement (%) at each ROI and the probability (%) of each of the five stimulation-induced manifestations (i.e., auditory hallucinations, receptive and expressive aphasia, speech arrest, and facial sensorimotor symptoms). The time bin during which the mean functional connectivity enhancement was significantly correlated with the probability of each stimulation-defined language-related function was identified. A Spearman’s rho greater than zero indicates a link between task-related functional connectivity enhancement in a specific 10-ms time bin and a specific language-related function indicated by electrical stimulation mapping. Given that we performed Spearman’s rank test for 250 time bins (i.e., 2,500-ms time periods), a Bonferroni-corrected p-value of <0.05 was considered significant.

## Data availability

The full dataset is publicly available at https://openneuro.org/datasets/ds005545/versions/1.0.2 (doi:10.18112/openneuro.ds005545.v1.0.2).

## Code availability

The analysis codes are available at https://github.com/a8k8nn0/TractographyAtlas (doi: 10.5281/zenodo.13957725).

## Results

### Patient profiles

A total of 106 patients with drug-resistant focal epilepsy, who met the inclusion and exclusion criteria, were included in the study (age range: 4-41 years; 49 females; 8 left-handed; 52 MRI-nonlesional cases). Detailed patient demographic information is provided in **Supplementary Table 1**.

### Impact of patient demographics and epilepsy-related profiles on response time

Univariate regression analysis confirmed that there was no excessive multicollinearity across √age, handedness, left-hemispheric epileptogenicity, MRI-visible lesion, and the number of antiseizure medications (range of Coefficient of Determination R^2^: 0.0001 to 0.0406; range of uncorrected p-values: 0.039 to 0.914). The mixed model analysis, incorporating all five variables as fixed effect factors, revealed that increased √age was associated with a reduction in the median response time (mixed model estimate: -0.246 [95% CI: -0.403 to -0.088]; t-value: -3.097; p-value: 0.003), independent of the other fixed effect variables. This finding indicates that each additional √year (e.g., from 9 to 16 years) was associated with a 246-ms reduction in response time. None of the other variables showed a significant association with median response time (range of p-values: 0.148 to 0.783).

### Impact of patient demographics, epilepsy-related profiles, and response time on neural dynamics

The mixed model analysis indicated that longer response time was associated with smaller high-gamma amplitude in the left precentral gyrus after stimulus offset. In other words, individuals with shorter response times showed earlier engagement of the left-hemispheric motor cortex, occurring 200-500 ms after stimulus offset. Specifically, each 1-second prolongation of response time was associated with reduced high-gamma amplitude at 400-500 ms after stimulus offset by -9.702% (95%CI: -13.122% to -6.281%; uncorrected p-value: 0.0000014; t-value: -5.746; DF: 37.095), at 300-400 ms after stimulus offset by -8.953% (95%CI: -12.195% to -5.710%; uncorrected p-value: 0.0000023; t-value: -5.596; DF: 36.790), and at 200-300 ms after stimulus offset by -8.092% (95%CI: -11.384% to -4.799%; uncorrected p-value: 0.0000154; t-value: -4.983; DF: 36.441).

The mixed model analysis also indicated that left handedness was associated with greater high-gamma amplitude in some right-hemispheric ROIs 200-500 ms before response onset. In other words, compared to right-handed individuals, left-handed individuals showed milder high-gamma attenuation in these right-hemispheric ROIs several hundred ms prior to overt responses. Specifically, left handedness increased right entorhinal high-gamma amplitude at 400-500 ms before response by 9.320% (95%CI: 5.390% to 13.249%; uncorrected p-value: 0.0000129; t-value: 4.741; DF: 62.000), at 300-400 ms before response by 11.677% (95%CI: 7.795% to 15.595%; uncorrected p-value: 0.0000030; t-value: 6.175; DF: 22.363), and at 200-300 ms before response by 12.598% (95%CI: 7.501% to 17.696%; uncorrected p-value: 0.0000361; t-value: 5.116; DF: 22.758). Left handedness likewise increased right supramarginal high-gamma amplitude at 400-500 ms before response by 14.391% (95%CI: 8.565% to 20.218%; uncorrected p-value: 0.0000121; t-value: 4.992; DF: 40.046). These observations indicate that functional connectivity diminution in the right parietal aslant fasciculus between the entorhinal and supramarginal gyri, occurring 400-500 ms before response, was 11.5% milder in left-handed individuals compared to right-handed ones.

### Statistical validity of the reported functional connectivity modulations

Significant high-gamma augmentation and attenuation were noted in 24.69% and 5.14% of the time bins, respectively, on average across ROIs, between stimulus onset and 500 ms after response onset. The chance probability of simultaneous significant high-gamma augmentation and attenuation at two or more ROIs lasting for ≥50 ms was 0.12% and 0.00000002%, respectively. In other words, the functional connectivity enhancement and diminution, reported below, had very low Type I error rates.

### Summarized view of functional connectivity modulations during auditory descriptive naming

**Supplementary Video 1** provides a comprehensive overview of task-related modulations in functional connectivity (**Figure 2**), along with cortical high-gamma dynamics (**Supplementary Figure 4**). **Figure 3** presents a probability map of stimulation-induced language-related manifestations, while **Figure 4** shows the specific time bins at which the intensity of functional connectivity enhancement was significantly correlated with the probability of a given stimulation-induced manifestation. **Figure 5** summarizes the duration (ms), extent (%: proportion of ROI pairs showing significant enhancement), and intensity (%) of functional connectivity enhancements within each fasciculus.

**Figure 2.**
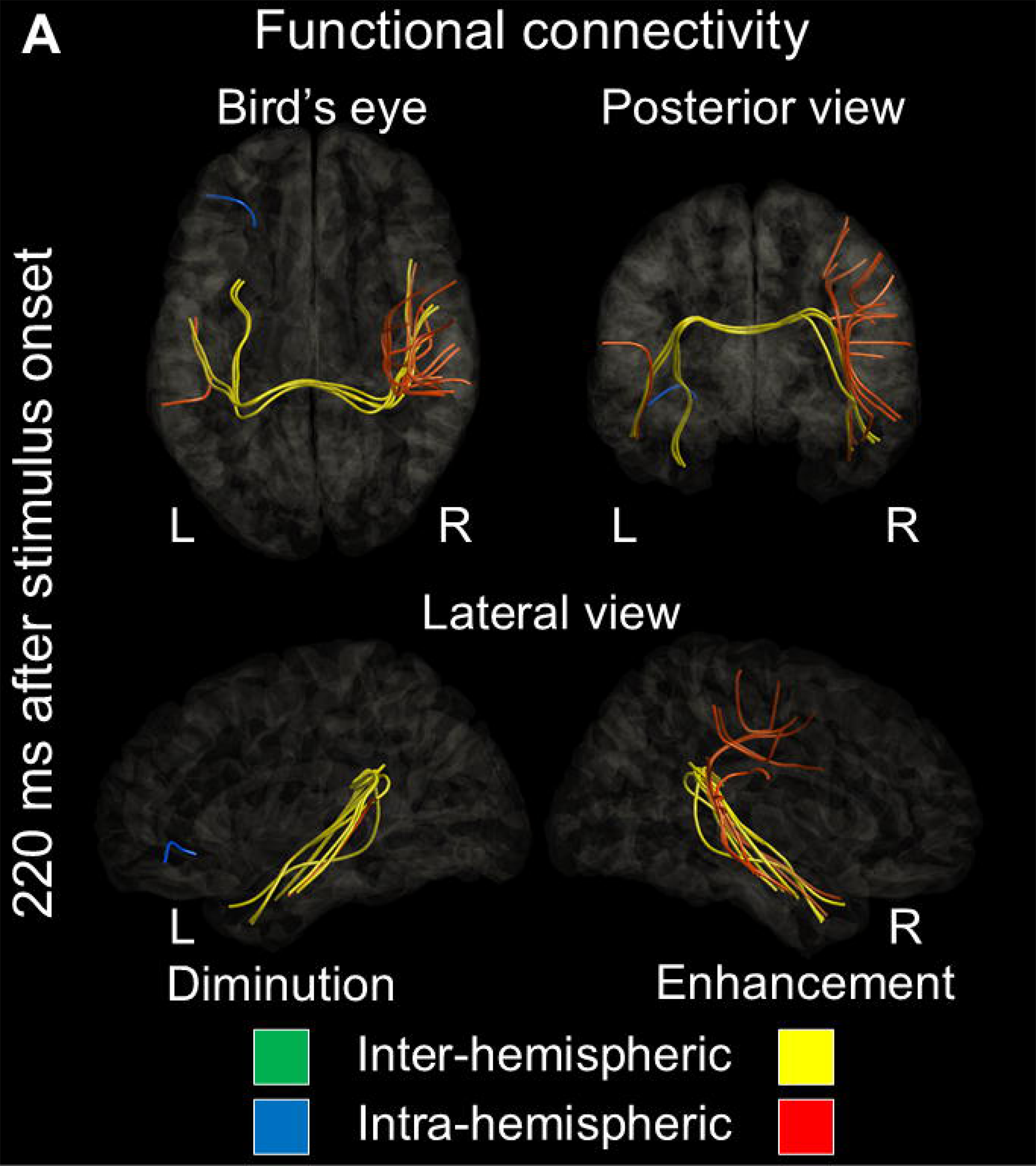
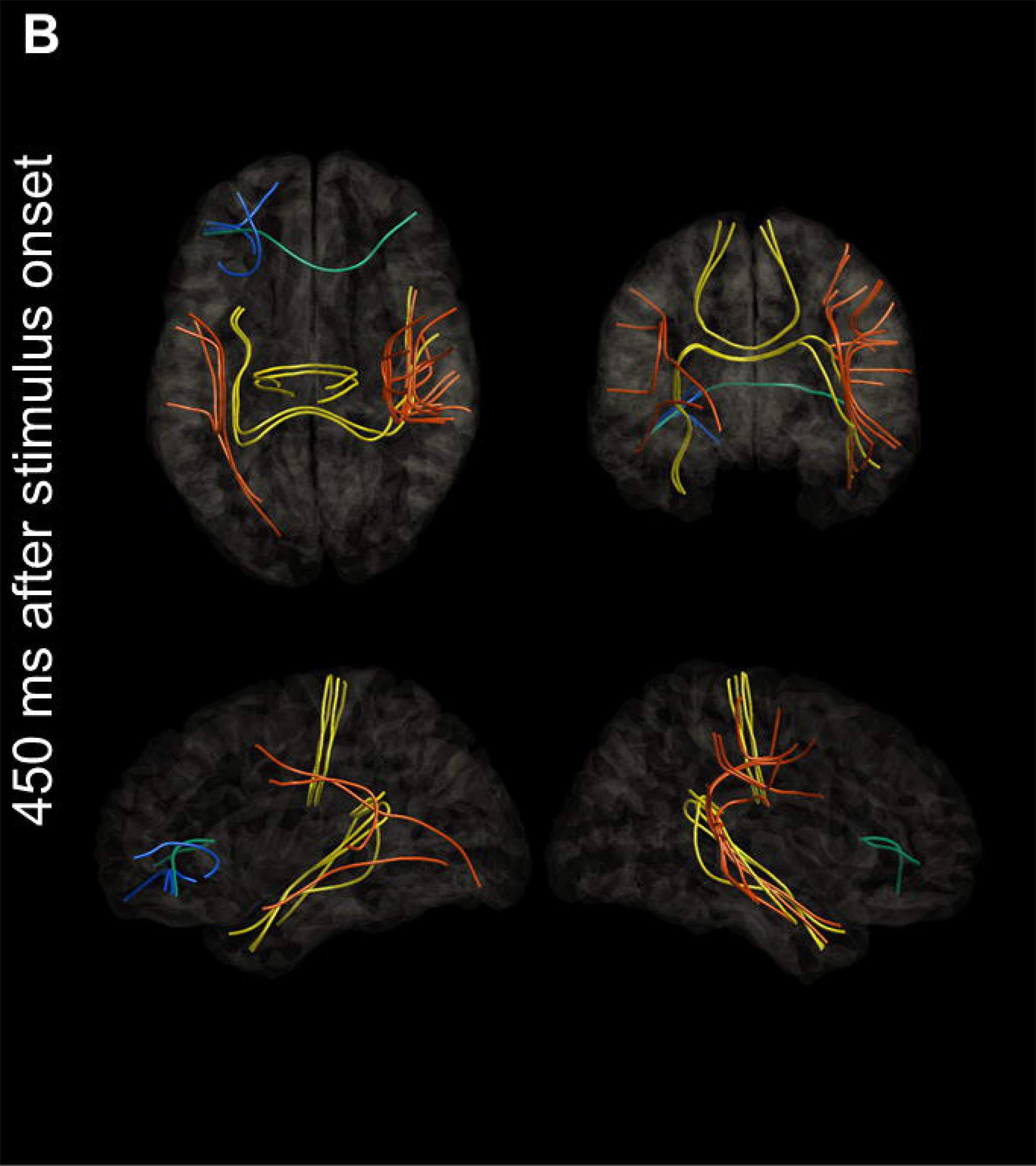
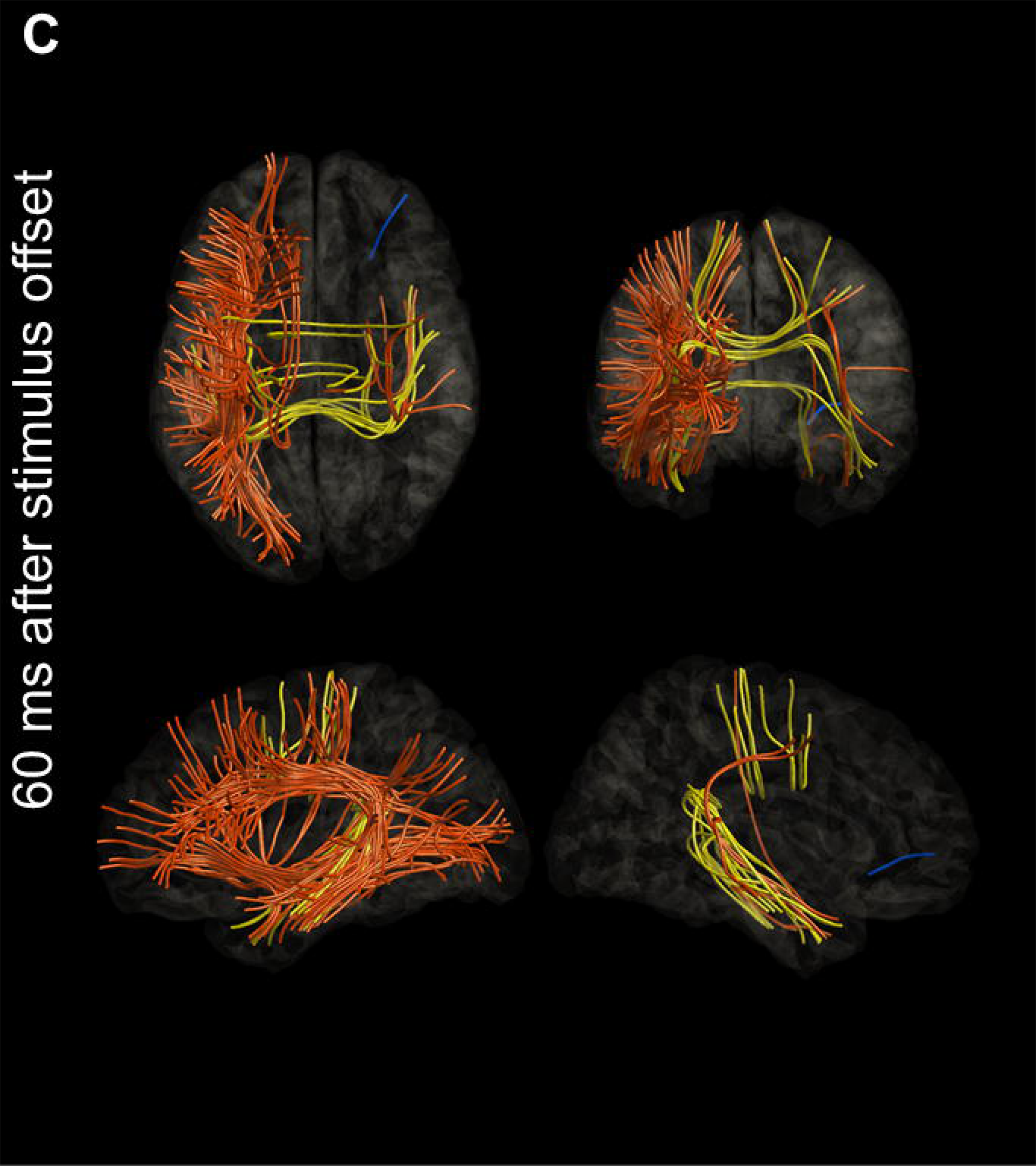
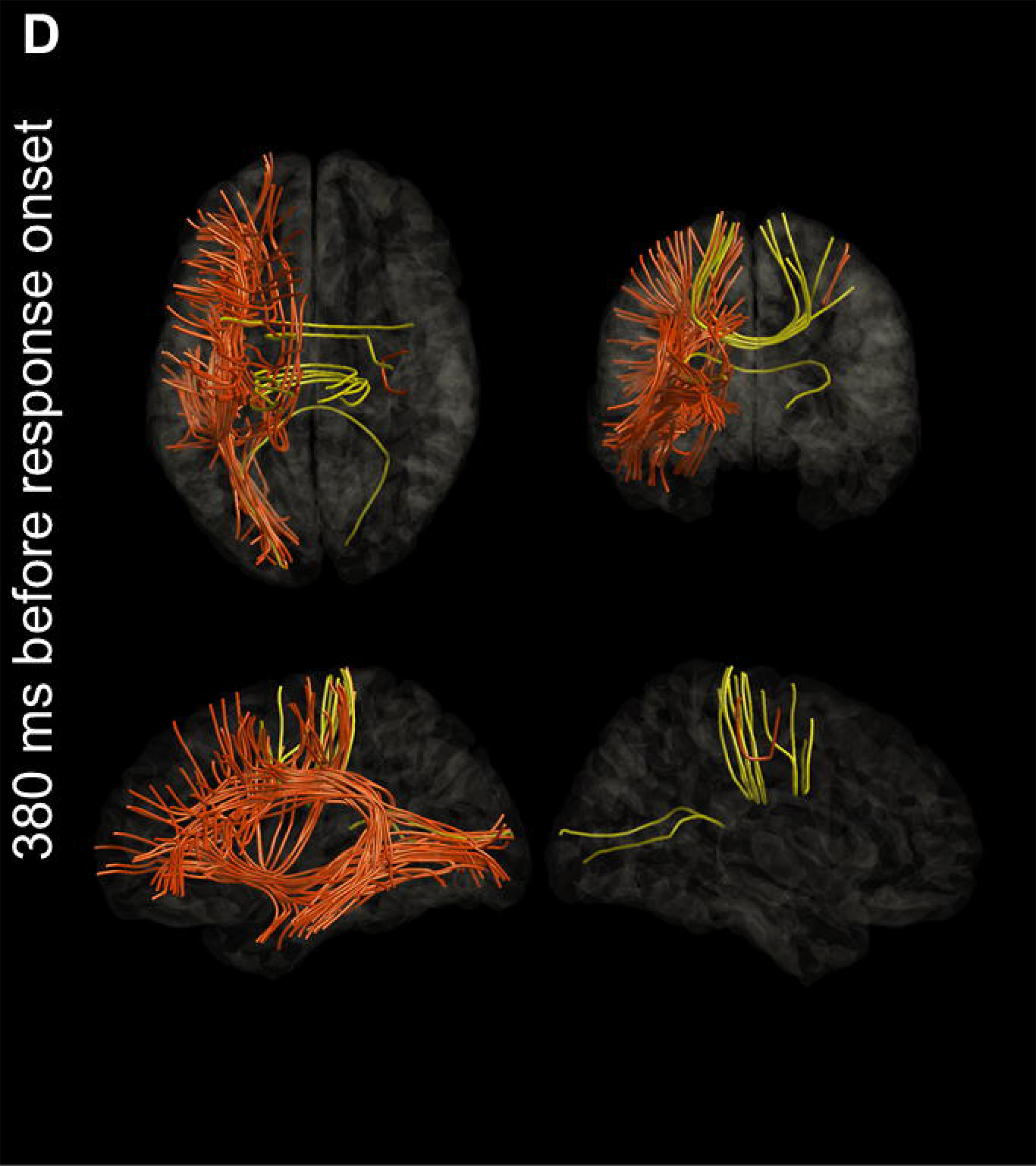
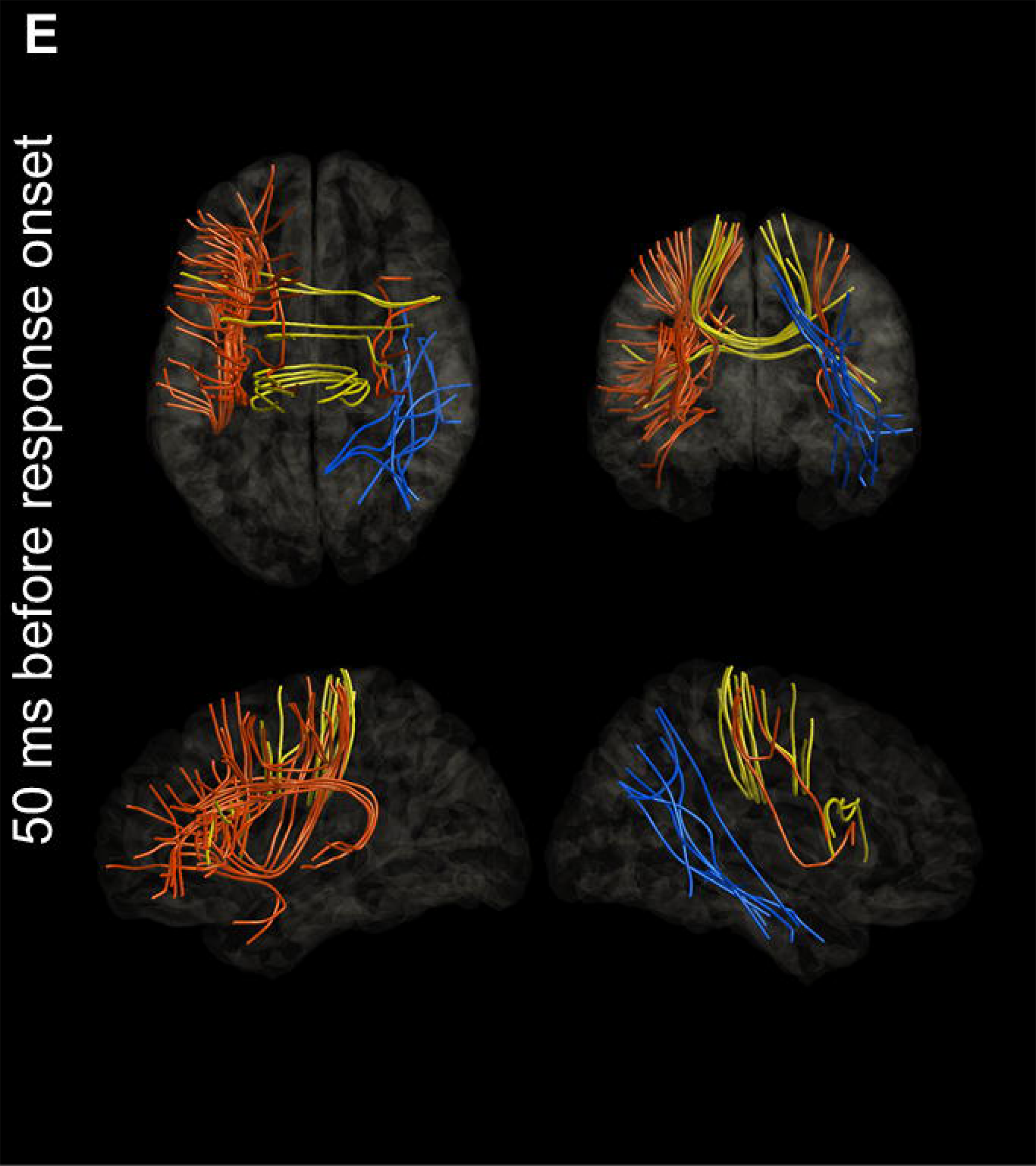
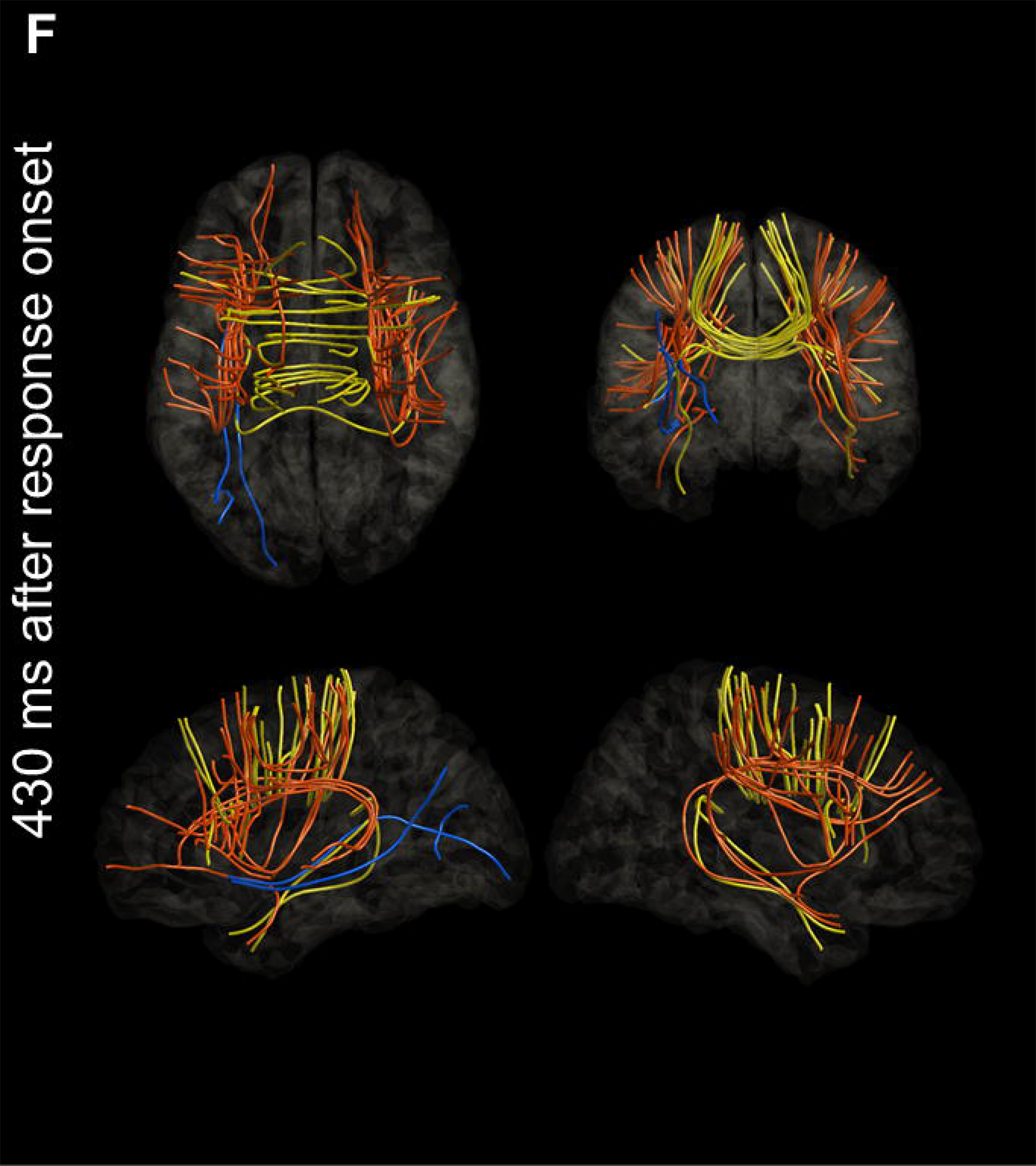
Naming-related modulations of functional connectivity. The snapshots present task-related modulations of functional connectivity. Orange and yellow streamlines: intra-hemispheric and inter-hemispheric functional connectivity enhancement. Blue and green streamlines: intra-hemispheric and inter-hemispheric functional connectivity diminution. **(A)** 220 ms after stimulus onset (association with auditory hallucination). **(B)** 450 ms after stimulus onset. **(C)** 60 ms after stimulus offset (association with receptive aphasia). **(D)** 380 ms before response onset (association with expressive aphasia). **(E)** 50 ms before response onset (association with speech arrest). **(F)** 430 ms after response onset (association with face sensorimotor symptoms). For a comprehensive overview of the network dynamics, please refer to **Supplementary Video 1**. L: left. R: right.

**Figure 3.**
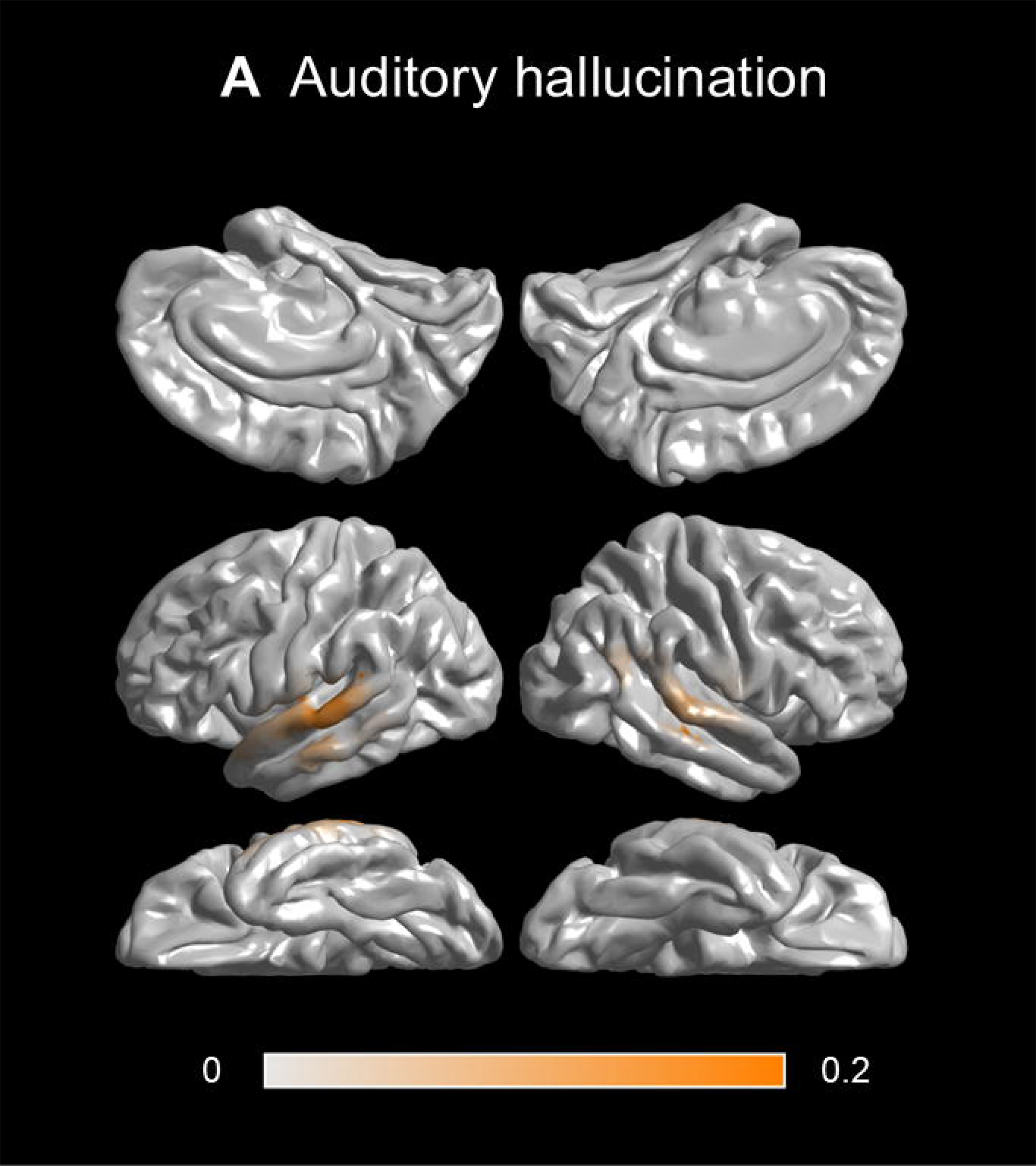
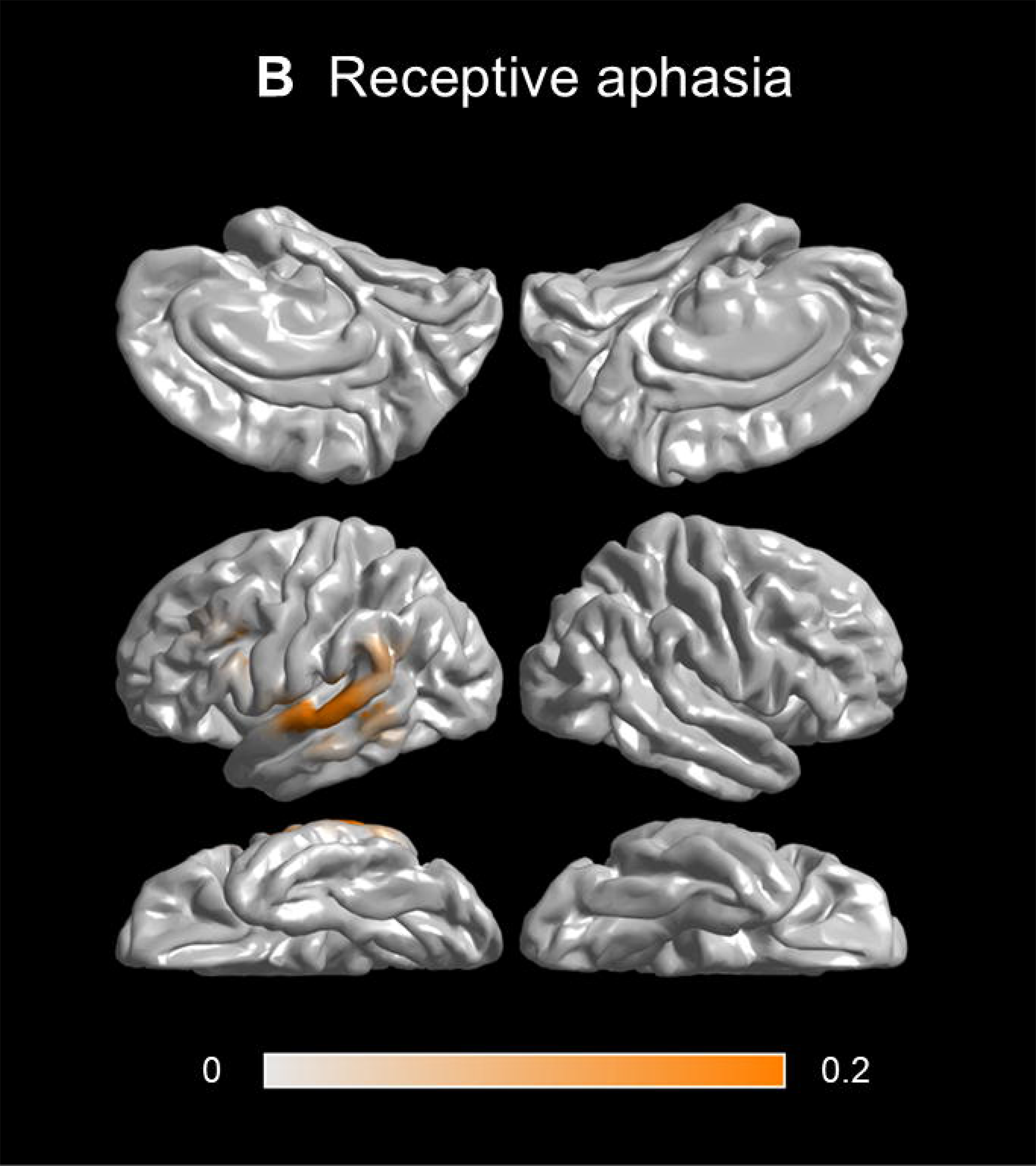
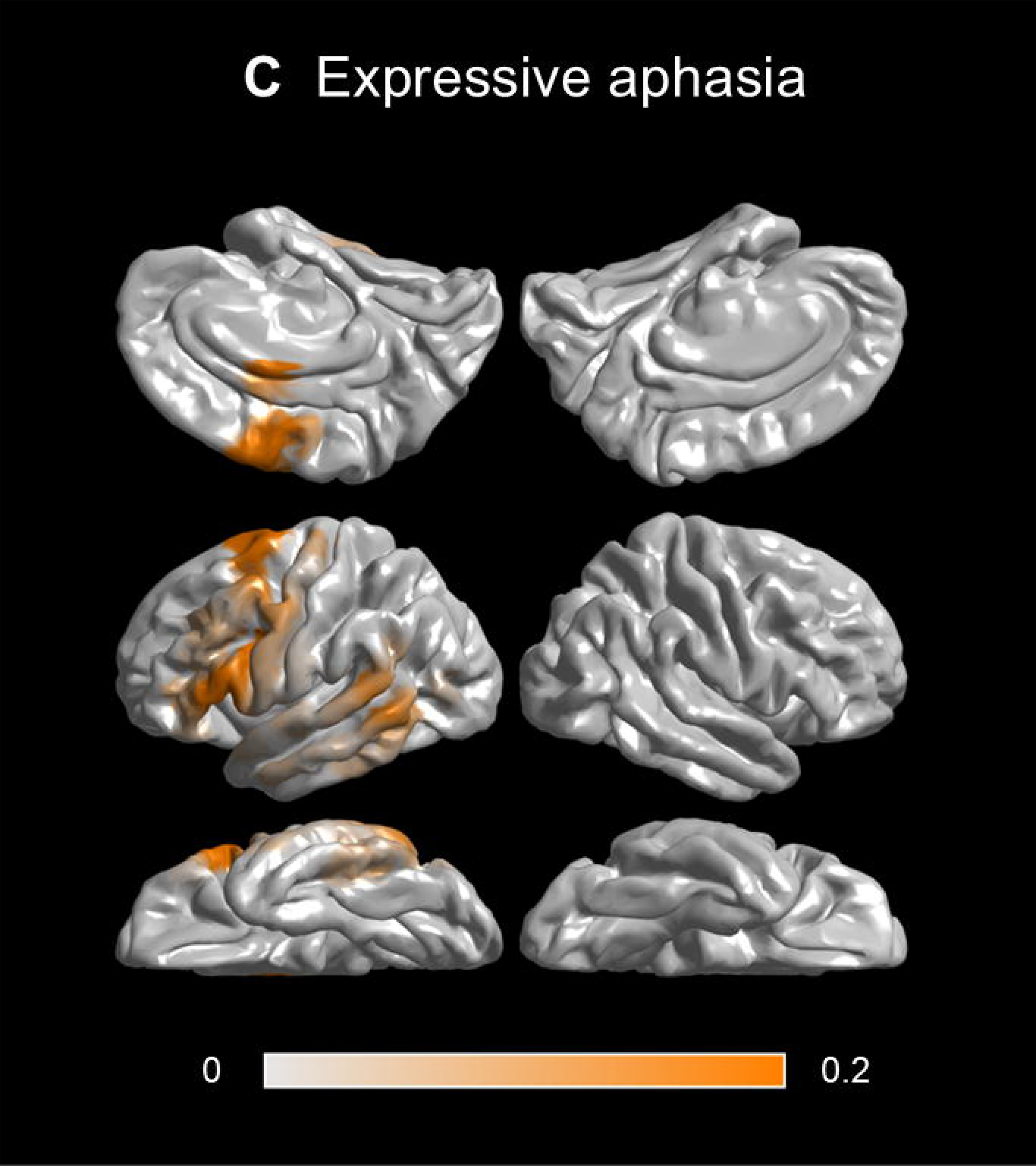
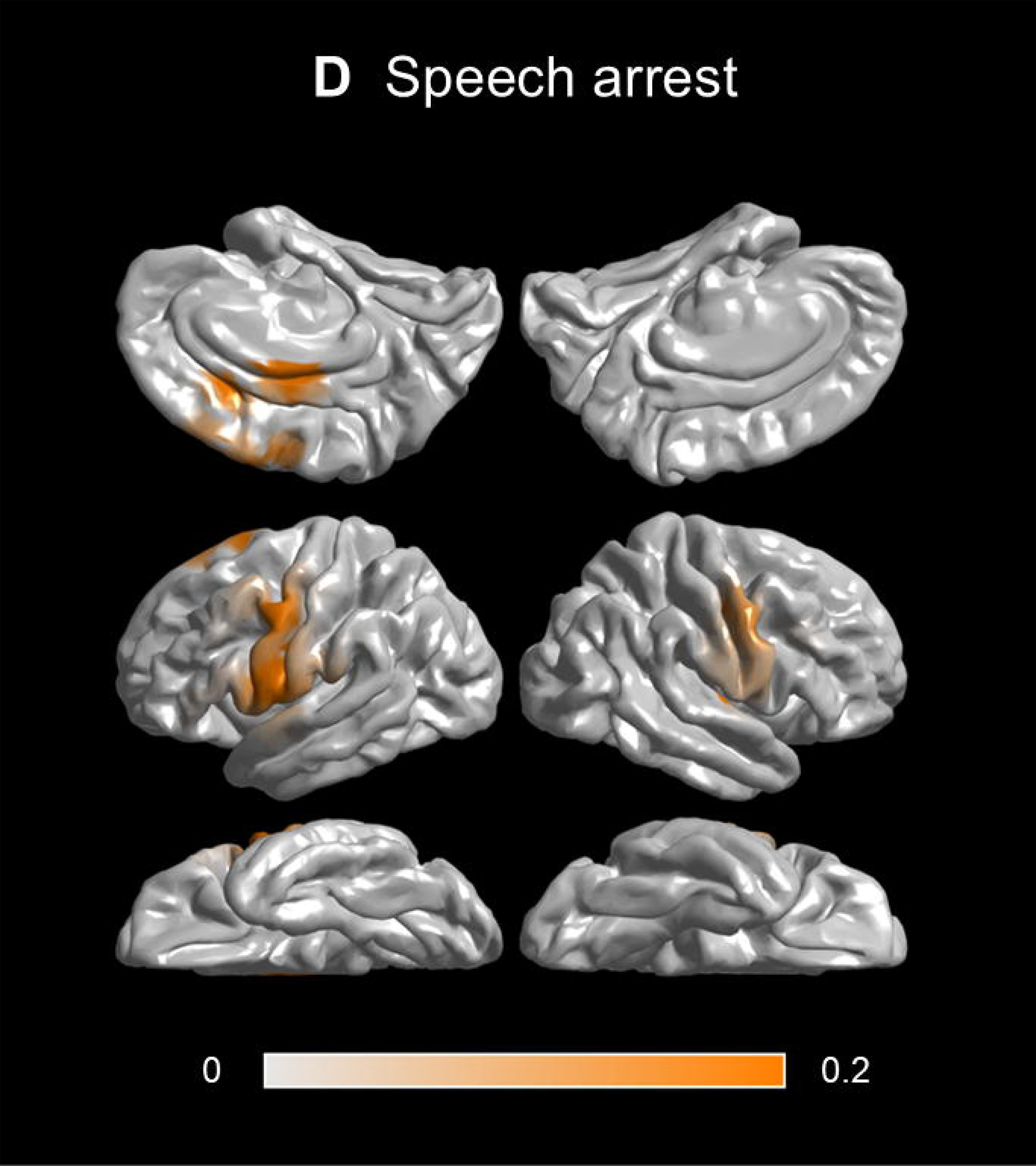
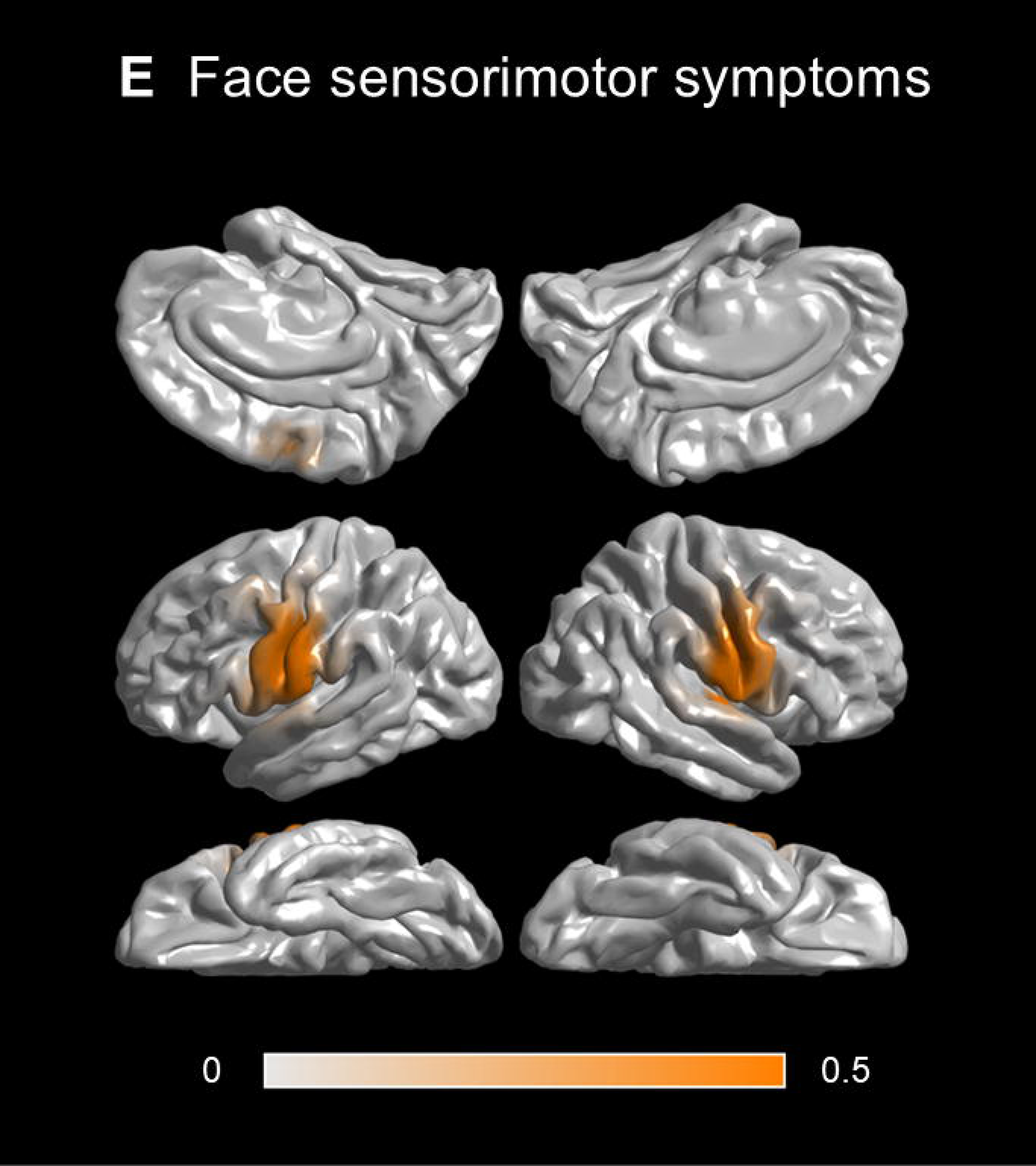
Probability map of stimulation-induced language-related symptoms. **(A)** Auditory hallucinations. **(B)** Receptive aphasia. **(C)** Expressive aphasia. **(D)** Speech arrest. **(E)** Face sensorimotor symptoms.

**Figure 4.**
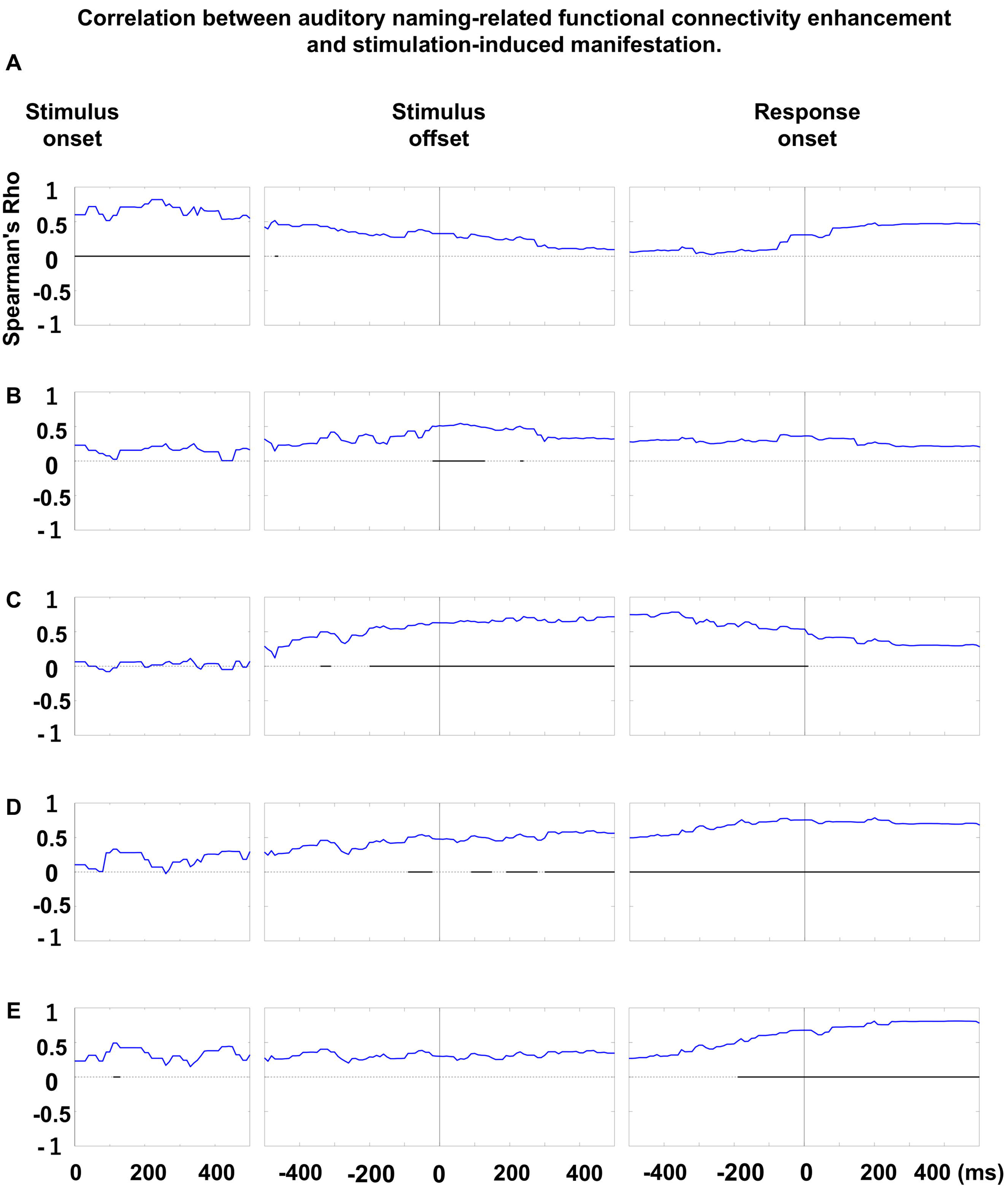
Association between functional connectivity and stimulation-defined language or speech-related function. Each plot displays Spearman’s rho, indicating the strength of the correlation between the probability of a given stimulation-induced manifestation and the mean functional connectivity enhancement at a given region of interest over time bins. **(A)** Auditory hallucination. **(B)** Receptive aphasia. **(C)** Expressive aphasia. **(D)** Speech arrest. **(E)** Facial sensorimotor symptoms. Horizontal bars indicate significant correlations based on a Bonferroni-corrected p-value of less than 0.05.

**Figure 5.**
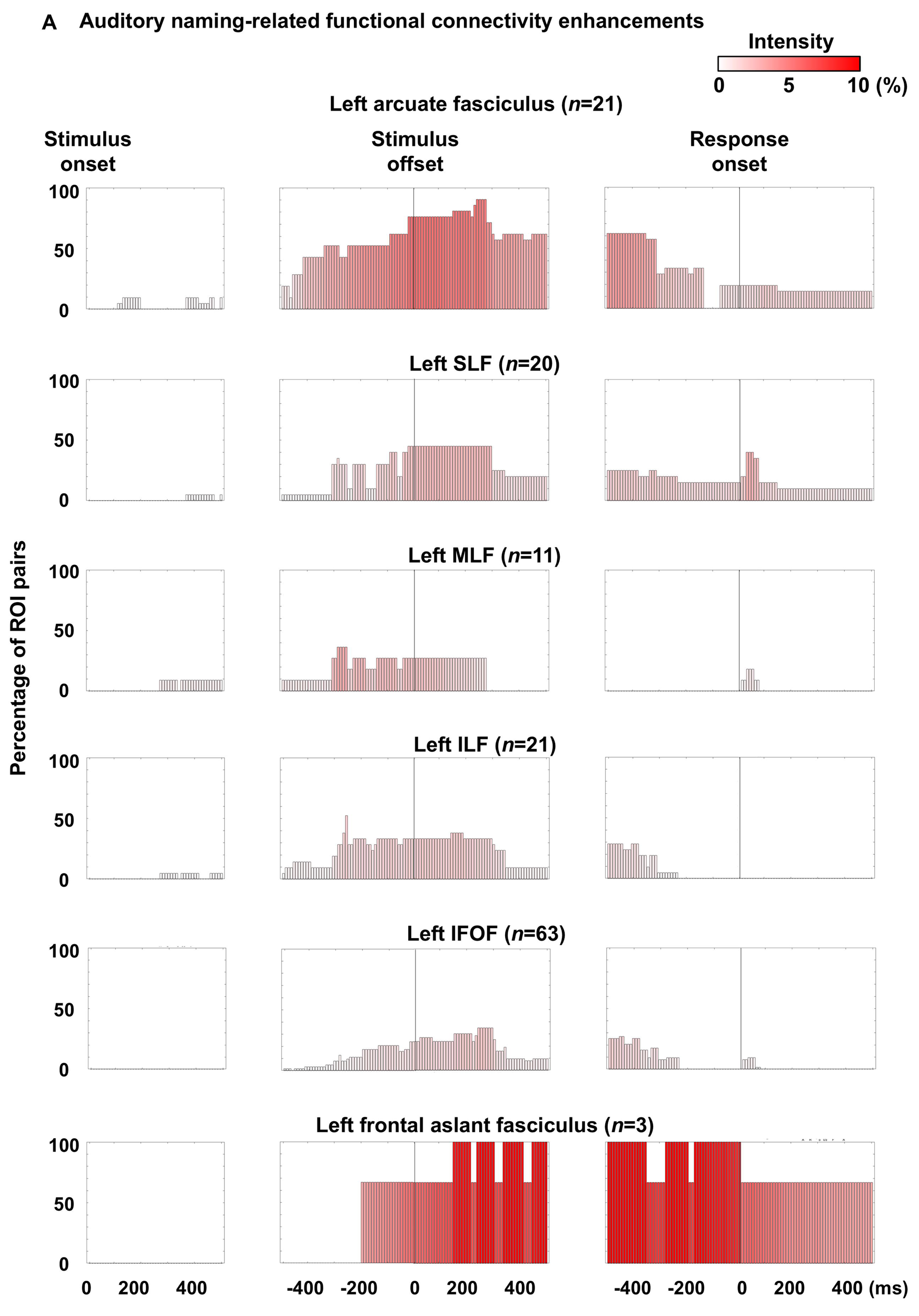
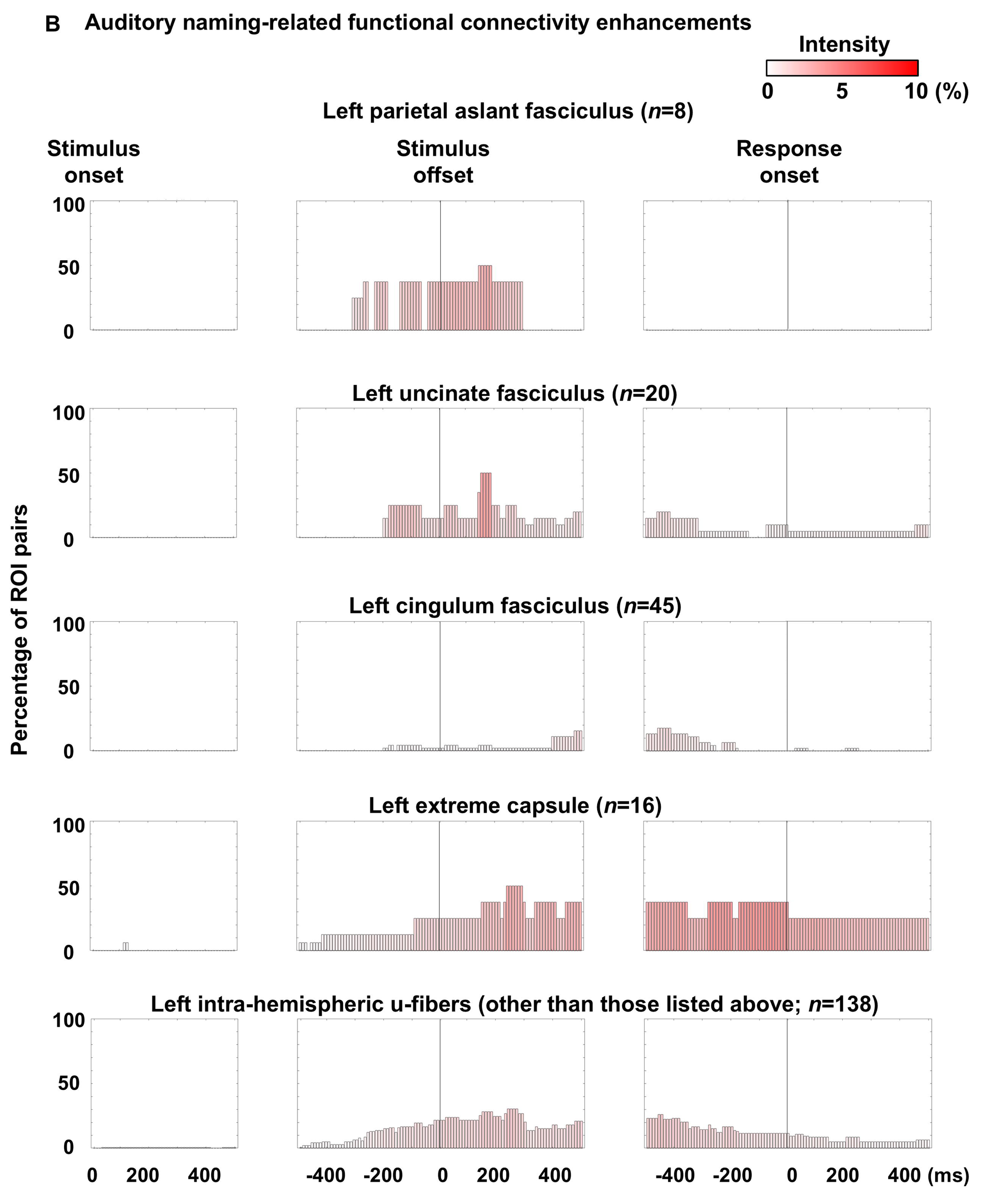
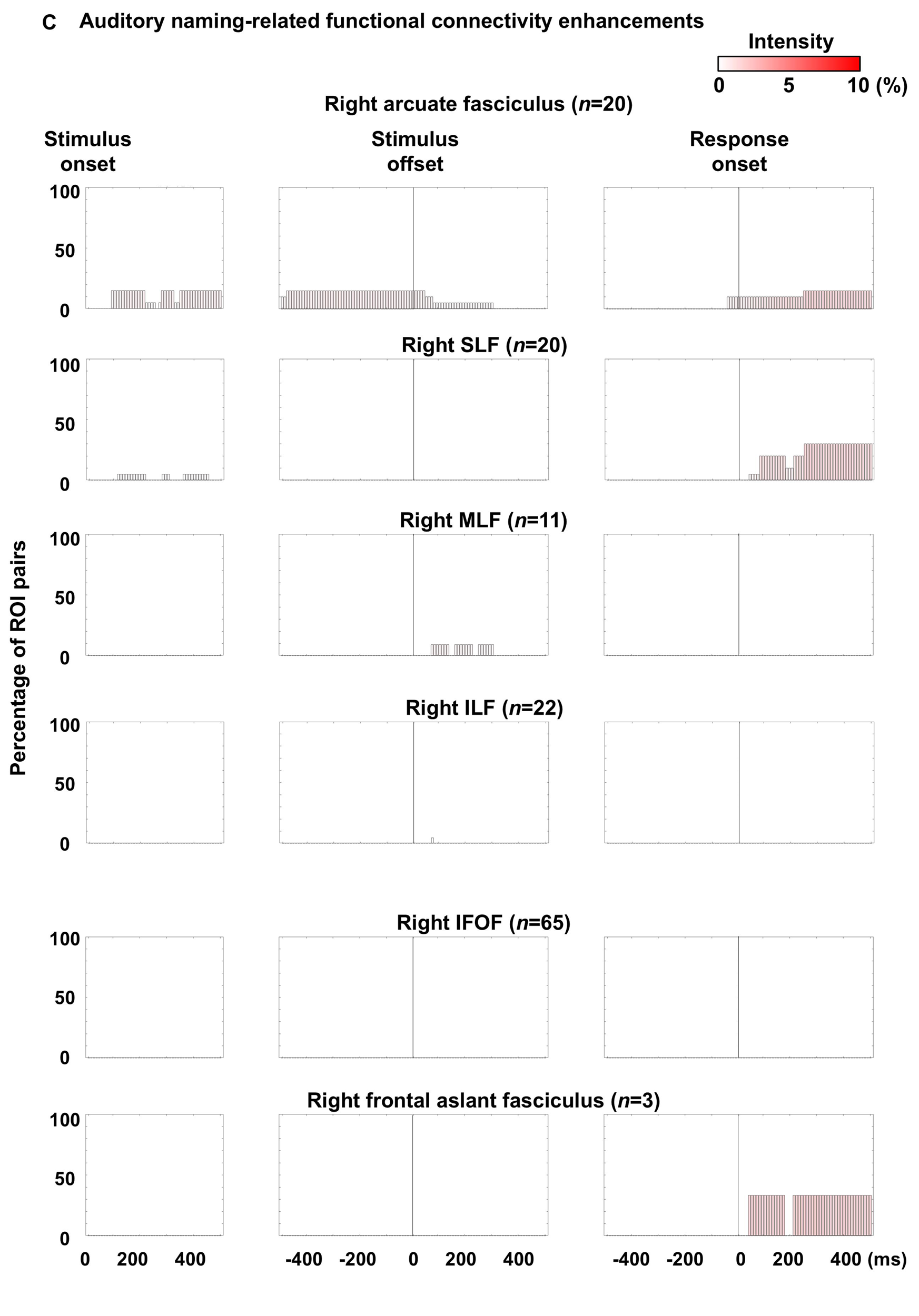
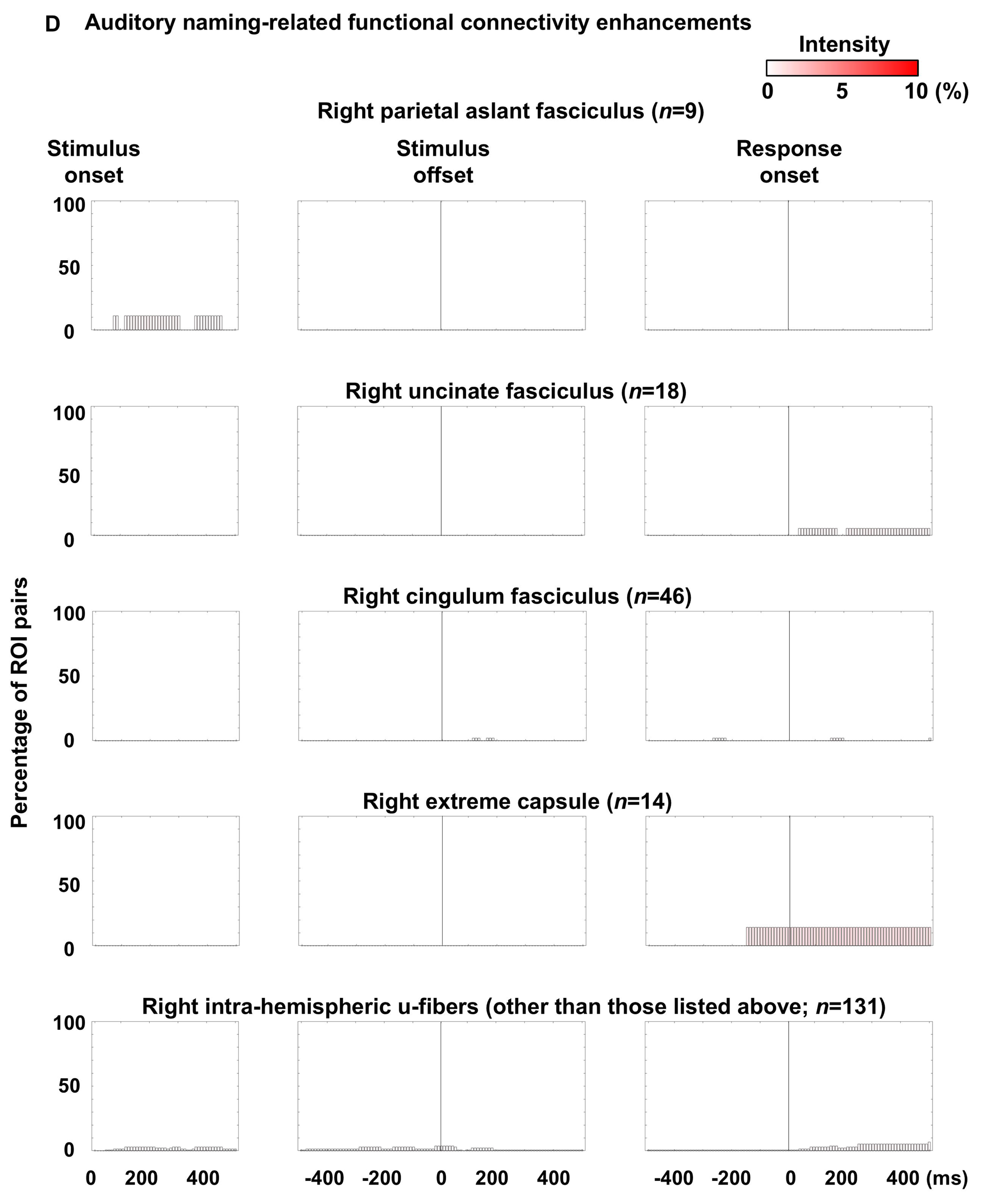
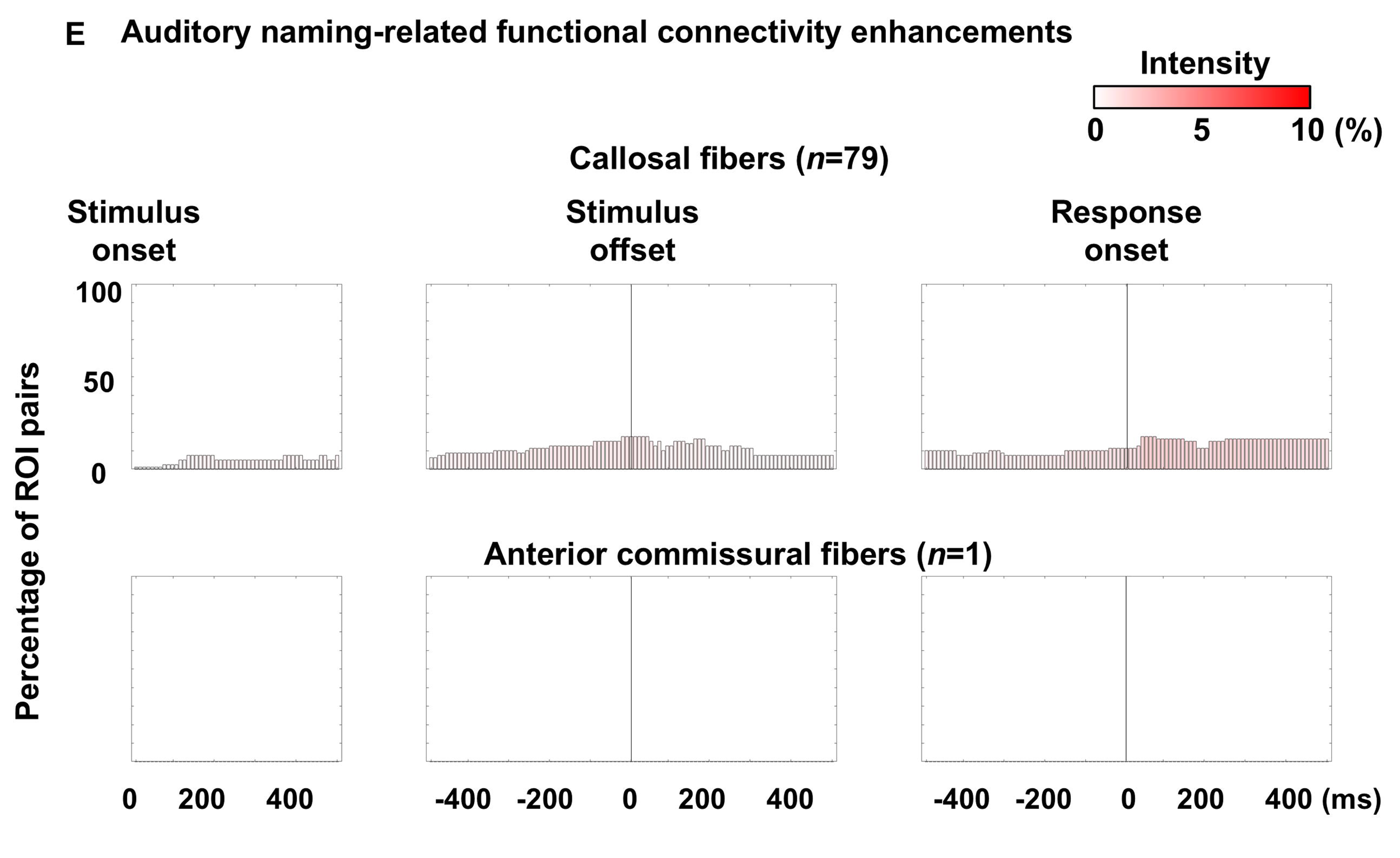
Dynamics of functional connectivity enhancement through each fasciculus. The bar height represents the proportion of functional connectivity enhancements within each fasciculus for a given time bin, while the bar color reflects the average intensity of enhancements within each fasciculus. **(A, B)** Left intra-hemispheric pathways. **(C, D)** Right intra-hemispheric pathways. **(E)** Inter-hemispheric pathways. **Supplementary Figure 5** presents the dynamics of functional connectivity diminution.

Upon hearing auditory stimuli, inter-hemispheric functional connectivity was enhanced between the superior temporal gyrus (STG) via the corpus callosum, while intra-hemispheric functional connectivity was enhanced between the STG and precentral gyrus through the arcuate fasciculus in each hemisphere (**Figure 2A**). The STG was also involved in functional connectivity enhancement through local u-fibers in each hemisphere. Conversely, the left IFG was involved in intra- and inter-hemispheric functional connectivity diminution through local frontal u-fibers and callosal fibers, while patients listened to auditory questions (**Figure 2B**). Subsequently, as stimulus offset approached, left intra-hemispheric functional connectivity enhancements became more evident through multiple fasciculi. After stimulus offset, left-hemispheric functional connectivity enhancement became extensive (**Figures 2C and 2D**), while right-hemispheric functional connectivity diminished (**Supplementary Figures 5C and 5D**). Around 100 ms prior to response onset, functional connectivity enhancement within the left frontal lobe regions persisted, while enhancement between the left frontal and temporal or occipital regions, including the arcuate and IFOF, diminished. Additionally, inter-hemispheric functional connectivity between the precentral gyri was observed through the corpus callosum immediately before and during responses (**Figure 2E**). During overt responses, intra-hemispheric functional connectivity between the STG and precentral gyrus continued to be enhanced through the arcuate fasciculus in each hemisphere. Inter-hemispheric functional connectivity enhancement between the Rolandic regions and between the STG persisted through the corpus callosum (**Figure 2F**).

### Electrical stimulation mapping

Stimulation-induced auditory hallucination, receptive aphasia, expressive aphasia, speech arrest, and face sensorimotor symptoms were observed at 50, 58, 146, 143, 518 sites, respectively. Auditory hallucination sites were primarily localized in the bilateral STG, whereas face sensorimotor sites were symmetrically distributed in the bilateral Rolandic cortices (**Figures 3A and 3E**). In contrast, receptive aphasia, expressive aphasia, and speech arrest sites were predominantly localized in the left hemisphere (**Figures 3B–3D**). Spearman’s rank correlation test did not reveal a correlation between patient age and the minimum stimulus intensity required to elicit a given clinical symptom (Bonferroni-corrected p-values: >0.05; detailed results are provided in the **Supplementary Document**).

### Functional connectivity associated with stimulation-induced auditory hallucinations

A significant association was noted between stimulation-induced auditory hallucination and functional connectivity enhancement upon hearing stimulus, and the strength of association was maximized at 220-250 ms after stimulus onset (rho: +0.82; uncorrected p-value: 6.6 × 10^-14^; t-value: +10.2; **Figure 4A**). **Figure 2A** presents the spatial characteristics of enhanced intra- and inter-hemispheric functional connectivity 220 ms after stimulus onset. Around this time, the callosal fiber connecting the left and right STG, as well as the arcuate fasciculi linking the STG and precentral gyrus, were among the major fasciculi exhibiting significant functional connectivity enhancement.

### Functional connectivity associated with stimulation-induced receptive aphasia

Similarly, a significant association was noted between stimulation-induced receptive aphasia and functional connectivity enhancement during a brief period around stimulus offset (**Figure 4B**), and the strength of association was maximized at 60 ms after stimulus offset (rho: +0.54; uncorrected p-value: 2.5 × 10^-5^; t-value: +4.6). **Figure 2C** presents the spatial characteristics of enhanced functional connectivity 60 ms after stimulus offset. At this time, significant connectivity enhancements were observed through multiple intra-hemispheric fasciculi in the left hemisphere, including local u-fibers (33/138 ROI pairs: 23.9%), arcuate (16/21: 76.2%), IFOF (17/63: 27.0%), SLF (9/20: 45.0%), ILF (7/21: 33.3%), extreme capsule (4/16: 25.0%), MLF (3/11: 27.2%), parietal aslant (3/8: 37.5%), uncinate (5/20: 25.0%), frontal aslant (2/3: 66.7%) and cingulum fasciculus (2/45: 4.4%). The callosal fibers (14/79 ROI pairs: 17.7%) also showed significant connectivity enhancement.

### Functional connectivity associated with stimulation-induced expressive aphasia

A significant association was likewise noted between stimulation-induced expressive aphasia and functional connectivity enhancement during a period between stimulus offset and response onset (**Figure 4C**), and the strength of association was maximized at 380 ms before response onset (rho: +0.78; uncorrected p-value: 3.9 × 10^-12^; t-value: +9.0). **Figure 2D** presents the spatial characteristics of enhanced functional connectivity 380 ms before response onset. At this time, significant connectivity enhancements were observed through multiple intra-hemispheric fasciculi in the left hemisphere, including local u-fibers (31/138 ROI pairs: 22.5%), arcuate (13/21: 61.9%), IFOF (16/63: 25.4%), extreme capsule (6/16: 37.5%), cingulum (6/45:13.3%), SLF (5/20: 25.0%), ILF (6/21: 28.6%), frontal aslant (3/3: 100%), and uncinate fasciculus (3/20: 15%). The callosal fibers (7/79 ROI pairs: 8.9%) also showed significant connectivity enhancement.

### Functional connectivity associated with stimulation-induced speech arrest

A significant association was noted between stimulation-induced speech arrest and functional connectivity enhancement around response onset (**Figure 4D**). Before response, the strength of association was maximized at 50 ms pre-response onset (rho: +0.78; uncorrected p-value: 7.8 × 10^-12^; t-value: +8.8). **Figure 2E** presents the spatial characteristics of enhanced functional connectivity 50 ms before response onset. At this time, significant connectivity enhancements were observed through multiple intra-hemispheric fasciculi in the left hemisphere, including local u-fibers (16/138 ROI pairs: 11.6%), extreme capsule (6/16: 37.5%), arcuate (4/21: 19.0%), frontal aslant (3/3: 100%), SLF (3/20: 15.0%), and uncinate fasciculus (2/20: 10.0%). The callosal fibers (8/79 ROI pairs: 10.1%) also showed significant connectivity enhancement.

### Functional connectivity associated with stimulation-induced face sensorimotor symptoms

A significant association was noted between stimulation-induced facial sensorimotor symptoms and functional connectivity enhancement around response onset, and the strength of association was maximized at 430 ms after response onset (rho: +0.81; uncorrected p-value: 2.0 × 10^-13^; t-value: +9.9; **Figure 4E**). **Figure 2F** presents the spatial characteristics of enhanced functional connectivity 430 ms after response onset.

### Disentangling functional connectivity from high-gamma amplitude effects

As a post hoc analysis, we examined how local high-gamma amplitude and functional connectivity correlated with the probability (%) of five stimulation-induced manifestations. For auditory hallucination, Spearman’s rho was +0.82 with high-gamma and +0.82 with functional connectivity at 220 ms after stimulus onset. For receptive aphasia, it was +0.51 (high-gamma) and +0.54 (functional connectivity) at 60 ms after stimulus offset. For expressive aphasia, it was +0.74 and +0.78, respectively, at 380 ms before response onset. For speech arrest, both correlations were +0.78 at 50 ms before response onset. For face sensorimotor symptoms, they were +0.82 (high-gamma) and +0.81 (functional connectivity) at 430 ms after response onset.

## Discussion

### Innovation and significance

For the first time, this study provides a dynamic causal tractography atlas that visualizes millisecond-scale functional connectivity modulations through specific fasciculi in both hemispheres, clarifying the causal roles of 12 major fasciculi during auditory descriptive naming. In other words, it visualizes the mesoscale spatiotemporal dynamics of neural coordination across white matter pathways within distributed cortical networks essential for understanding and responding to spoken questions. We identified these causal roles by correlating the intensity of functional connectivity enhancements with the probability of speech-related clinical manifestations induced by electrical stimulation at the cortical sites where these fasciculi originate. The study revealed the sequential and alternating engagement of fasciculi with causal roles in auditory descriptive naming. No single fasciculus was consistently engaged throughout the task; rather, each fasciculus was active only during specific stages, including auditory perception, semantic-syntactic analysis, lexical retrieval, speech planning/initiation, and sensorimotor processes for articulation. Functional connectivity effects on electrical stimulation findings cannot be attributed solely to cortical high-gamma amplitudes. We found that incorporating functional connectivity through direct white matter pathways increased correlations with receptive and expressive aphasia by 5-6%, yet did not enhance correlations with sensory or motor manifestations. This observation may be consistent with the notion that semantic and syntactic processing, as well as lexical retrieval, rely on extensively distributed networks, whereas sensorimotor functions are mediated by more localized cortical structures.

Additionally, the study demonstrated that multiple distinct major fasciculi contribute simultaneously to each of these stages. Our data provide compelling evidence that enhanced functional connectivity at discrete time windows is associated with distinct language or speech-related functions, as confirmed by electrical stimulation mapping. Quantitative analysis further supported our hypothesis that the left arcuate fasciculus exhibited the earliest, most intense, and sustained connectivity enhancements. We also found that functional connectivity was highly left-hemispheric dominant several hundred milliseconds before the response, yet this left-hemispheric dominance appeared milder in left-handed individuals. Collectively, the evidence presented in this study advances current neurobiological models of speech network organization.

The robustness of our findings is supported by analyzing naming-related high-gamma amplitudes from 7,792 intracranial electrode sites across 106 patients, which, to our knowledge, represents the largest sample size reported to date. White matter streamlines were delineated using high-quality DWI tractography data from 1,065 healthy individuals in the HCP dataset (Yeh et al., 2018). The strict statistical criteria defining significant functional connectivity modulations ensured that our observed connectivity enhancements and diminutions were not attributable to chance. Additionally, we enhanced the generalizability of our atlas by clarifying the impacts of age, handedness, and epilepsy-related profiles on task-related high-gamma modulations. We released our white matter streamline template, intracranial EEG data, and analytic toolbox as open-source resources to facilitate replication of our analyses and construction of personalized dynamic tractography atlases. Below, we discuss the significance of functional connectivity modulations in major fasciculi as well as intra-hemispheric u-fibers not classified among the 12 major fasciculi.

### Arcuate fasciculus

As shown in **Figure 5**, the arcuate fasciculus contributes to auditory descriptive naming for the longest periods among the 12 major fasciculi studied in the current study. The arcuate fasciculus was defined as a dorsal fiber bundle connecting the temporal lobe to the frontal or parietal lobe (**Supplementary** Figure 3B) (Thiebaut de Schotten et al., 2011; Dick and Tremblay, 2012; Hansen et al., 2021; Yeh, 2022). Upon hearing auditory questions, the bilateral arcuate fasciculi between the STG and precentral gyri exhibited functional connectivity enhancement. Up to 10-15% of the arcuate pathways in each hemisphere demonstrated functional connectivity enhancements (**Figure 5**). These specific portions of the arcuate fasciculi comprises an articulatory/phonological loop, and the bilateral enhancement of its connectivity is attributed to auditory-motor transformation and short-term storage of speech stimuli via subvocal rehearsal (Baddeley, 2003; Warren et al., 2005; Bernal and Ardila, 2009; Cogan et al., 2014; Kambara et al., 2017; Nishida et al., 2017). Auditory-motor transformation refers to the conversion of sensory representations of sound into motor representations, a process suggested to be essential for speech repetition (Warren et al., 2005; Cogan et al., 2014). Behavioral observations from MRI-visible lesions, electrical stimulation, and hemodynamic responses on functional MRI (fMRI) suggest that the left arcuate fasciculus terminating at the precentral gyrus is crucial for the short-term storage and repetition of spoken words without requiring lexical retrieval (Bernal and Ardila, 2009; Janssen et al., 2023; Zhao et al., 2023). For example, lesions in the arcuate fasciculus connecting the left STG and precentral gyrus frequently result in speech repetition impairments (Catani et al., 2005; Bernal and Ardila, 2009). Our prior iEEG studies demonstrated that a task requiring sound repetition elicited high-gamma augmentation in the bilateral STG and precentral gyri simultaneously, and that single-pulse electrical stimulation of these high-gamma sites revealed direct connections through the arcuate fasciculi (Nishida et al., 2017; Sonoda et al., 2021). We also previously reported that high-gamma augmentation in the precentral gyri during the working memory maintenance period increased with memory load, indicating the involvement of the precentral gyri in the short-term storage of speech stimuli via subvocal rehearsal (Kambara et al., 2017). The bilateral functional connectivity enhancement of the arcuate fasciculi observed in this study is consistent with the notion that auditory-motor transformation is bilaterally distributed (Warren et al., 2005; Cogan et al., 2014). This bilateral distribution supports the ability to continue performing speech repetition even after lexical retrieval impairments following acute lesions in the left arcuate fasciculus (Shuren et al., 1995; Berthier et al., 2012).

Between the question offset and 300 ms before response onset, functional connectivity enhancement in the left arcuate fasciculus was extensive, whereas connectivity enhancement in the right arcuate fasciculus was less pronounced (**Figure 5**). During this period, 60–90% of the left arcuate pathways showed enhanced functional connectivity, while only 0–15% of the right arcuate pathways exhibited similar enhancements. Enhanced connectivity in the left arcuate fasciculus originated mainly from the STG, middle temporal gyrus (MTG), and inferior temporal gyrus (ITG), extending to the precentral gyrus, posterior inferior frontal gyrus (pIFG), and posterior middle frontal gyrus (pMFG). The probability of stimulation-induced receptive and expressive aphasia was best correlated with the intensity of connectivity enhancement at 60 ms after stimulus offset (**Figure 2C**) and 380 ms before response onset (**Figure 2D**), respectively. In other words, the functional connectivity enhancements shown in these figures likely reflect coordinated neural activity required for comprehending questions and generating relevant responses; among all pathways studied, the functional connectivity enhancement through the left arcuate fasciculus was the most prominent component.

These findings support the notion that the left arcuate fasciculus plays a dominant role in semantic-syntactic analysis and lexical retrieval. Our observations align with lesion studies, which show that damage to the left, but not the right, arcuate fasciculus leads to significant impairments in these cognitive processes (Bernal and Ardila, 2009; Marchina et al., 2011; Wilson et al., 2011; Fridriksson et al., 2013; Griffiths et al., 2013; Gajardo-Vidal et al., 2021; Giampiccolo and Duffau, 2022). A study of 134 stroke survivors demonstrated that persistent impairments in semantic analysis and lexical retrieval were causally associated with damage to the anterior portion of the left arcuate fasciculus rather than the overlying pIFG (Gajardo-Vidal et al., 2021). Additionally, investigators have reported that electrical stimulation of the left arcuate fasciculus transiently impairs semantic analysis and lexical retrieval (Duffau et al., 2002), and that stimulation of the IFG, connected to the temporal lobe neocortex via the arcuate fasciculus, transiently impairs syntactic analysis (Riva et al., 2022).

Neither the left nor right arcuate fasciculus exhibited significant functional connectivity enhancement 80- to 130-ms prior to response onset (**Figure 5**). The increased intensity of functional connectivity during this critical time was closely correlated with the probability of stimulation-induced speech arrest (**Figure 4D**), with the left extreme capsule, SLF and frontal aslant fasciculus among the white matter pathways showing enhanced connectivity. These findings suggest that the arcuate fasciculus plays a minimal or modest role in the planning and initiation of overt responses. An alternative interpretation is that the human brain engages in response planning and initiation only after the left arcuate fasciculus completes semantic-syntactic analysis and lexical retrieval. We could not find any prior studies that have distinctly separated the impact of arcuate fasciculus lesions on response initiation from their effects on semantic-syntactic processing or lexical retrieval.

After the onset of overt response, 10–20% of the bilateral arcuate pathways, including those connecting the STG and Rolandic areas, exhibited functional connectivity enhancements (**Figure 5**). A plausible explanation for this observation is that the arcuate fasciculus plays a role in monitoring one’s own voice and making responses with appropriate vocal tone (Zheng et al., 2010; Chang et al., 2013; Greenlee et al., 2013). A prior study reported that an acute lesion involving the right arcuate and SLF resulted in impaired prosody perception (Sammler et al., 2018). Another interpretation is that the functional connectivity enhancement during this period reflects the activation of the articulatory/phonological loop through the arcuate fasciculi for the short-term storage of one’s own speech (Baddeley, 2003; Kambara et al., 2017).

### Superior longitudinal fasciculus (SLF)

The SLF dorsally connects the parietal-occipital regions and the frontal lobe (**Supplementary Figure 3C**) (Catani et al., 2005; Dick and Tremblay, 2012). The functional connectivity enhancement in the left SLF was extensive and intensified between 300 ms before stimulus offset and response onset (**Figure 5**), while the right SLF showed connectivity diminution (**Supplementary Figure 5C**). Specifically, during this period, up to 15-45% of the left SLF pathways showed significant connectivity enhancements, whereas up to 10% of the right SLF pathways showed significant connectivity diminutions. Given the correlation between functional connectivity intensity and stimulation-induced receptive aphasia, expressive aphasia and speech arrest (**Figure 4**), the left SLF appears to play a role in semantic-syntactic analysis, lexical retrieval, and response initiation. Previous studies in patients undergoing tumor surgery have reported that surgical damage involving the left SLF and ILF was associated with postoperative impairments in lexical retrieval (Herbet et al., 2016; Tomasino et al., 2018). A study of 46 chronic post-stroke patients found that damaged fasciculi associated with impaired semantic processing included the left ILF as well as the left posterior cortex overlying the SLF (Alyahya et al., 2020). Additionally, a study involving 11 patients with brain tumors reported that electrical stimulation of the anterior-inferior portion of the left SLF resulted in speech arrest (Maldonado et al., 2011).

### Middle longitudinal fasciculus (MLF) and inferior longitudinal fasciculus (ILF)

The temporal dynamics of functional connectivity enhancement was similar between the left MLF and ILF. Between 300 ms before and after stimulus offset, 25-40% of these fasciculi demonstrated enhanced functional connectivity, with the intensity and extent gradually diminishing afterward (**Figure 5**). Given the association between increased functional connectivity intensity and stimulation-induced receptive aphasia (**Figure 4**), and considering that receptive aphasia can result from impairments in phonological or semantic analysis, the functional connectivity enhancements observed in the left MLF and ILF likely play roles in these processes.

The MLF is a ventral fiber bundle connecting the parietal or occipital lobe to the superior temporal gyrus (**Supplementary Figure 3D**). The functional role of the left MLF has been previously understudied. A DWI tractography study involving 29 healthy adult individuals reported that the integrity of the MLF was correlated with phoneme discrimination performance (Tremblay et al., 2019). Another DWI tractography study of 20 patients with primary progressive aphasia reported that the loss of integrity of the left MLF was correlated with impairment in word comprehension and retrieval but not with articulation or fluency (Luo et al., 2020). A study of eight patients with brain tumors reported that picture-naming errors induced by stimulation and resection were attributed to the effects of the left IFOF, but not the left MLF (De Witt Hamer et al., 2011). A future study differentiating the patterns of functional connectivity enhancement between auditory and picture naming tasks may provide insights into the functional role of the left MLF.

The ILF ventrally connects the occipital or parietal lobe to the temporal lobe (**Supplementary Figure 3E**). The functional connectivity enhancement dynamics in the left ILF are consistent with the notion that the left ILF contributes to semantic analysis, as indicated by previous studies. A DWI tractography study demonstrated that the anatomical integrity of the left ILF was impaired in 11 children with object recognition impairments compared to age- and sex-matched typically developing children (Ortibus et al., 2012). Additionally, a study of 58 patients with brain tumors reported an association between lesions involving the left ILF and poor performance in a famous face naming task (Burkhardt et al., 2023). Another study of 12 brain tumor patients found that electrical stimulation of the left ILF did not induce language-related manifestations, but a subset of patients developed transient naming errors following resection involving the left ILF (Mandonnet et al., 2007).

### Inferior fronto-occipital fasciculus (IFOF)

The IFOF ventrally connects the occipital-posterior temporal regions to the frontal lobe through the external capsule (**Supplementary Figure 3F**). The extent and intensity of functional connectivity enhancement in the left IFOF was initiated prior to stimulus offset, maximized around 300 ms after stimulus offset, and persisted until 240 ms before response onset (**Figure 5**). At 300 ms after stimulus offset, 35% of the left IFOF pathways showed significant functional connectivity enhancement. Given the correlation between functional connectivity intensity and stimulation-induced expressive and receptive aphasia (**Figure 4**), the left IFOF appears to play roles in both lexical retrieval and semantic analysis. A study of 15 stroke survivors reported that lesions in the left IFOF are associated with impaired semantic analysis (Harvey and Schnur, 2015). A study of 31 patients with brain tumor reported that infiltration to the left IFOF was associated with impaired semantic analysis (Zhao et al., 2023). Another study of 12 patients with brain tumor reported that electrical stimulation of the left IFOF elicited anomia (Mandonnet et al., 2007).

### Frontal aslant fasciculus

The frontal aslant fasciculus connects the IFG and the superior frontal gyrus (SFG) while running immediately medial to the anterior SLF (**Supplementary Figure 3G**). Functional connectivity enhancement in the left frontal aslant fasciculus was observed after stimulus offset and sustained until overt responses (**Figure 5**). All three frontal aslant pathways demonstrated significant functional connectivity enhancements at 380 ms and 50 ms before response onset, when connectivity intensity best correlated with stimulation-induced expressive aphasia and speech arrest, respectively (**Figure 4**). These observations suggest that the left frontal aslant fasciculus may play a causal role in transferring the retrieved word to speech preparation and initiation. Electrical stimulation and lesions involving the left frontal aslant fasciculus have been reported to result in speech arrest and impairments in movement initiation (Vassal and Boutet, 2014; Kinoshita et al., 2015; Chernoff et al., 2018).

### Parietal aslant fasciculus

The parietal aslant fasciculus is a fiber bundle coursing laterally to the lateral ventricle, connecting the parietal lobe to the temporal lobe (**Supplementary Figure 3H**). Functional connectivity enhancement was observed in up to 50% of the left parietal aslant pathways during the 600-ms period around stimulus offset, with intensity peaking at 150-190 ms after stimulus offset (**Figure 5**). Given the strong correlation between functional connectivity intensity and stimulation-induced receptive aphasia (**Figure 4**), the left parietal aslant fasciculus appears to play a role in semantic comprehension of questions. A previous case study reported that intraoperative electrical stimulation of the left parietal aslant fasciculus elicited semantic paraphasias (Chidambaram et al., 2023).

During the 400-ms period before response onset, up to 45% of the right parietal aslant pathways exhibited diminished functional connectivity (**Supplementary Figure 5D**). Additionally, left-handedness was associated with a milder reduction in functional connectivity within the right parietal aslant fasciculus. The causal role of this fasciculus in naming has not been previously clarified. One possible explanation for our observations is that the right parietal aslant fasciculus is not essential for lexical retrieval; thus, its neural coordination decreases to reallocate resources to left-hemispheric networks, particularly in right-handed individuals.

### Uncinate fasciculus

The uncinate fasciculus is a fiber bundle connecting the anterior portion of the temporal lobe to the ventral portion of the frontal lobe through the temporal stem (**Supplementary Figure 3I**). Between 200 ms before stimulus offset and 150 ms before response onset, 10-50% of the left uncinate fasciculus exhibited increased functional connectivity (**Figure 5**). Given the observed correlation between functional connectivity intensity and stimulation-induced responses (**Figure 4**), the left uncinate fasciculus likely contributes to semantic analysis and lexical retrieval. This fasciculus remains one of the least understood among the major fasciculi. Investigators infer that the left uncinate fasciculus plays a role in semantic and syntactic analyses (Friederici and Gierhan, 2013; Alyahya et al., 2020). A study of 10 patients with left-hemispheric stroke reported that impairment in semantic control was correlated with reduced resting-state functional connectivity on fMRI and decreased anatomical integrity of the uncinate fasciculus on DWI tractography (Harvey et al., 2013). Conversely, a study of 13 patients with brain tumors reported that neither electrical stimulation nor resection of the left uncinate fasciculus significantly impaired speech processes (Duffau et al., 2009).

### Cingulum fasciculus

The cingulum fasciculus is a fiber bundle coursing medially to the lateral ventricle, connecting the occipital, parietal, or temporal lobe to the frontal lobe (**Supplementary Figure 3J**). Between 400 ms after stimulus offset and 200 ms before response onset, 10-20% of the left cingulum pathways showed significant functional connectivity enhancements (**Figure 5**). Given the correlation between functional connectivity intensity and stimulation-induced manifestations (**Figure 4**), the left cingulum fasciculus appears to play a role in retrieval of words. A study of 29 patients with drug-resistant focal epilepsy demonstrated that naming errors were elicited by electrical stimulation of the left medial frontal and parietal regions via depth electrodes positioned around the cingulum fasciculus (Perrone-Bertolotti et al., 2020). Additionally, a study of five patients with intractable pain found that stimulation of the left, but not the right, cingulum impaired short-term recall of visual objects (Fedio and Ommaya, 1970).

### Extreme capsule

The extreme capsule was defined as a fiber bundle connecting the precentral, postcentral, or paracentral gyrus to the IFG or insular gyrus through the external capsule (**Supplementary Figure 3K**) (Yeh et al., 2018). Between stimulus offset and response onset, as well as during overt articulation, 25-50% of the left extreme capsule exhibited enhanced functional connectivity, with connectivity intensity highest prior to response onset. The observed correlation between functional connectivity intensity and stimulation-induced symptoms (**Figure 4**) suggests that the left extreme capsule may play a causal role in word retrieval and speech initiation. A study of 123 patients with left hemispheric stroke showed that lesions in the left extreme capsule was an independent predictor of aphasia, defined by the Toke Test (Martinez Oeckel et al., 2021). A study of 38 patients with brain tumor reported that intraoperative stimulation of the left extreme capsule elicited picture naming error (Papagno et al., 2011).

### Intra-hemispheric u-fibers

Intra-hemispheric u-fibers, defined as those outside the 12 major fasciculi, were extensively distributed (**Supplementary Figure 3L**). Between stimulus offset and response onset, more than 10% of these u-fibers in the left hemisphere exhibited enhanced functional connectivity (**Figure 5**). This observation suggests that, rather than consistent enhancement of the same u-fiber, specific pairs of neighboring cortices were transiently engaged at different locations and time windows, facilitating local coordination at distinct stages of the naming process.

Key observations reveal that functional connectivity via local u-fibers involving the left IFG or precentral gyrus was enhanced 400 ms after question offset and subsided after response onset (**Figure 2D**). Given the correlation between functional connectivity intensity and stimulation-induced expressive aphasia and speech arrest (**Figure 4**), this enhancement likely reflects lexical retrieval and speech planning following the semantic-syntactic analysis of questions. An fMRI study in 26 healthy individuals reported that increased hemodynamic activation in the pars triangularis of the left IFG was linked to semantic processing, while activation in the left precentral gyrus and pars opercularis of the left IFG was related to phonological processing. (Katzev et al., 2013) A study of 134 stroke survivors demonstrated that long-term impairments in lexical retrieval were more strongly associated with damage to the anterior portion of the left arcuate fasciculus than with damage to the left IFG (Gajardo-Vidal et al., 2021).

Additionally, we observed that functional connectivity via local u-fibers in the left IFG was transiently diminished during the listening phase of questions (**Figure 2B**). This reduction suggests a suppression of neural resources required for lexical retrieval in response to hearing *wh-*questions. Such suppression may optimize neural resource allocation, (Norman, 1975) enhancing auditory processing by reducing cognitive load in the left IFG and its associated white matter tracts. This prioritization may facilitate the perception of question stimuli within the STG and connected fasciculi.

### Callosal fibers

Callosal fibers connect the left and right hemispheres via the corpus callosum (**Supplementary Figure 3M**). Enhancement of inter-hemispheric functional connectivity through these fibers was observed throughout the task (**Figure 5**), with different portions of the callosal fibers engaged at various stages of the naming process. Upon hearing the questions, enhanced connectivity was prominent in the posterior portion of the corpus callosum (**Figure 2A**). In contrast, prior to and during overt responses, functional connectivity enhancement was more pronounced in the anterior corpus callosum (**Figure 2F**). A possible explanation for the enhancement of inter-hemispheric functional connectivity during stimulus listening is that it integrates neural representations of temporal speech features, such as the timing and rhythm of speech sounds, processed by the left STG, (Zatorre and Belin, 2001; Chang et al., 2010; Floegel et al., 2020) and spectral features, including tone and pitch, processed by the right STG (Zatorre and Belin, 2001; Floegel et al., 2020). One possible explanation for the enhancement of inter-hemispheric functional connectivity during overt responses is that it supports the coordination of symmetric and synchronous movements of the mouth structures, which are essential for speech initiation and overt articulation (Hoptman and Davidson, 1994; Hervé et al., 2013). The causal role of the corpus callosum in initiating speech is highlighted by reports of mutism immediately after callosotomy (Sussman et al., 1983; Crutchfield et al., 1994).

### Anterior commissural fibers

The anterior commissural fibers are those connecting the left and right temporal lobes through the anterior commissure (**Supplementary Figure 3N**). In the present study, only one connecting between the left and right ITG was found in our tractography analysis. We failed to observe functional connectivity enhancement through the anterior commissural fibers (**Figure 5**).

### Impacts of patient demographics on neural measures

In our study cohort, older patients exhibited shorter response times compared to younger patients. Specifically, each additional √year (e.g., from 9 to 16 years) was associated with a 0.244-second reduction in response time. This shorter response time associated with greater high-gamma augmentation in the left precentral gyrus between 200-500 ms after question offset. For every one-second reduction in response time, there was an 8-9% increase in high-gamma amplitude in the left precentral gyrus during this window. These findings suggest that individuals with shorter response times activate the left precentral gyrus earlier, and that older individuals tend to respond more quickly. Therefore, caution is needed when interpreting our dynamic tractography atlas at 200-500 ms after question onset, as age-related differences in response time may influence connectivity patterns. According to our model, young adults may exhibit about 2% greater functional connectivity enhancement in the left precentral gyrus during this time window compared to school-aged children. We did not observe a significant effect of response time on neural measures time-locked to question onset or response onset.

We also found that left-handedness was linked to higher high-gamma amplitude (i.e., less degree of high-gamma attenuation) in certain right-hemispheric ROIs 200-500 ms before response onset. Specifically, compared to right-handed individuals, left-handed individuals exhibited high-gamma amplitudes higher in the right entorhinal cortex and right supramarginal gyrus by 9-14%. This finding suggests that left-hemispheric dominant neural coordination during the lexical retrieval period is more enhanced in right-handed individuals, whereas left-handed individuals show a less pronounced left-hemispheric dominance. Our observation is consistent with fMRI studies reporting that left-handed individuals, compared to right-handed ones, less frequently exhibit left-hemispheric dominant hemodynamic activation during language tasks. (Szaflarski et al., 2002; Mazoyer et al., 2014) Aside from handedness, we did not find any other patient demographics or epilepsy-related profiles associated with high-gamma measures in the present study.

### Methodological considerations

The spatial extent of intracranial electrode placement was determined solely by clinical needs, resulting in iEEG signal sampling from limited areas in each patient. To overcome these spatial limitations, we aggregated iEEG data across multiple individuals and used the group mean to assess network dynamics, aiming to understand shared neural processes. Several iEEG investigators have inferred neural interactions within this framework (Kunii et al., 2013; Burke et al., 2014; Avanzini et al., 2016; Solomon et al., 2017). To account for inter-individual differences in behavior and neural dynamics, we thoroughly evaluated potential confounding factors, specifically considering patient age, handedness, epileptogenic hemisphere, MRI-visible lesions, antiseizure polytherapy, and response time. A higher number of antiseizure medications indicates a greater seizure-related cognitive burden (Kwan and Brodie, 2001). Response times also serve as a cognitive performance measure for the naming task. Thus, our analytic approach accounts for the effect of a patient’s cognitive skill on neural measures. Including too many covariates in the mixed model analysis could introduce collinearity issues. In the **Supplementary Document**, we provide the results of ancillary analyses for interested readers, examining (1) the relationship between a history of bilateral tonic–clonic seizures and task behaviors, and (2) the relationship between patient age and the stimulus intensity required to induce clinical manifestations.

To define the spatial extent of ROIs, we utilized the Desikan-Killiany atlas, one of the most widely used atlases in neuroscience research (Desikan et al., 2006), and delineated white matter streamlines connecting pairs of these ROIs. Due to the varying sizes of the ROIs, we opted not to perform statistical comparisons of the extent or intensity of functional connectivity modulations between different fasciculi within a given hemisphere, as such comparisons could be misleading. However, we considered it feasible to compare the extent or intensity of functional connectivity modulations within the same fasciculus across different time windows. We acknowledge that our analytical approach resulted in false negatives when delineating genuine streamlines between ROIs. To ensure clarity in our visualizations, we chose to delineate a single white matter streamline for each ROI pair. This approach was necessary to effectively visualize all major fasciculi in bird’s-eye, posterior, and lateral views. Including too many lateral fasciculi would obscure the visualization of medial fasciculi, while an excess of superior fasciculi would hinder the visibility of inferior fasciculi.

The principle of defining functional connectivity enhancement in our study parallels that used in fMRI-based studies, where distinct cortical ROIs with statistically similar task-related hemodynamic activation patterns are considered functionally connected. (Sun et al., 2004; Crossley et al., 2013; Glasser et al., 2016) By incorporating the presence of direct streamlines connecting these cortical ROIs as a criterion, the current study enhanced the biological plausibility of the observed connectivity enhancements. It is suggested that cortical regions with simultaneous high-frequency neural responses are involved in coordinated interactions (Singer, 1993; Buonomano and Merzenich, 1998). Our previous iEEG study demonstrated that single-pulse electrical stimulation at a cortical site elicited neural responses within 50 ms specifically at distinct cortical sites showing significant, simultaneous, and sustained naming-related high-gamma augmentation (Sonoda et al., 2021). Collective evidence supports the notion that neural coordination can plausibly occur between these cortical ROIs directly connected by white matter streamlines.

We inferred the causal roles of specific fasciculi using clinical manifestations induced by cortical stimulation. Subcortical stimulation, often performed for adult patients with brain tumor during awake craniotomy, was not conducted in our study cohort, as it was infeasible given that most participants were children and the eloquent cortex was identified through standard-care management during extraoperative iEEG monitoring (Nakai et al., 2017; Sonoda et al., 2022).

## Abbreviations

CI: confidence interval
DTI: diffusion tensor imaging
DWI: diffusion-weighted imaging
fMRI: functional magnetic resonance imaging
HCP: Human Connectome Project
IFOF: inferior fronto-occipital fasciculus
ILF: inferior longitudinal fasciculus
ITG: inferior temporal gyrus
iEEG: intracranial electroencephalography
MLF: middle longitudinal fasciculus
MTG: middle-temporal gyrus
MNI: Montreal Neurological Institute
pIFG: posterior inferior frontal gyrus
pMFG: posterior middle frontal gyrus
ROI: region of interest
SOZ: seizure onset zone
SFG: superior frontal gyrus
SLF: superior longitudinal fasciculus
STG: superior temporal gyrus

## Acknowledgments

We are grateful to Sandeep Sood, MD, Alanna Carlson, MS, LLP, and Jamie MacDougall, RN, BSN, CPN at Children’s Hospital of Michigan for the collaboration and assistance in performing the studies described above.

## Supplementary material

Supplementary material is available online.

## References

Almairac, F., Herbet, G., Moritz-Gasser, S., Menjot de Champfleur, N., Duffau, H., 2015. The left inferior fronto-occipital fasciculus subserves language semantics: A multilevel lesion study. Brain. Struct. Funct. 220, 1983–1995. 10.1007/s00429-014-0773-1.

Alyahya, R.S.W., Halai, A.D., Conroy, P., Lambon Ralph, M.A., 2020. A unified model of post-stroke language deficits including discourse production and their neural correlates. Brain. 143, 1541–1554. 10.1093/brain/awaa074.

Anumanchipalli, G.K., Chartier, J., Chang, E.F., 2019. Speech synthesis from neural decoding of spoken sentences. Nature. 568, 493–498. 10.1038/s41586-019-1119-1.

Arya, R., Horn, P.S., Crone, N.E., 2018. ECoG high-gamma modulation versus electrical stimulation for presurgical language mapping. Epilepsy. Behav. 79, 26–33. 10.1016/j.yebeh.2017.10.044.

Asano, E., Juhász, C., Shah, A., Sood, S., Chugani, H.T., 2009. Role of subdural electrocorticography in prediction of long-term seizure outcome in epilepsy surgery. Brain. 132, 1038–1047. 10.1093/brain/awp025.

Avanzini, P., Abdollahi, R.O., Sartori, I., Caruana, F., Pelliccia, V., Casaceli, G., Mai, R., Lo Russo, G., Rizzolatti, G., Orban, G.A., 2016. Four-dimensional maps of the human somatosensory system. Proc. Natl. Acad. Sci. U. S. A. 113, E1936–E1943. 10.1073/pnas.1601889113.

Baddeley, A., 2003. Working memory and language: An overview. J. Commun. Disord. 36, 189–208. 10.1016/S0021-9924(03)00019-4.

Ball, T., Kern, M., Mutschler, I., Aertsen, A., Schulze-Bonhage, A., 2009. Signal quality of simultaneously recorded invasive and non-invasive EEG. Neuroimage. 46, 708–716. 10.1016/j.neuroimage.2009.02.028.

Bernal, B., Ardila, A., 2009. The role of the arcuate fasciculus in conduction aphasia. Brain. 132, 2309–2316. 10.1093/brain/awp206.

Berthier, M.L., Lambon Ralph, M.A., Pujol, J., Green, C., 2012. Arcuate fasciculus variability and repetition: The left sometimes can be right. Cortex. 48, 133–143. 10.1016/j.cortex.2011.06.014.

Boatman, D., Freeman, J., Vining, E., Pulsifer, M., Miglioretti, D., Minahan, R., Carson, B., Brandt, J., McKhann, G., 1999. Language recovery after left hemispherectomy in children with late-onset seizures. Ann. Neurol. 46, 579–586. 10.1002/1531-8249(199910)46:4<579::AID-ANA5>3.0.CO;2-K

Buonomano, D.V., Merzenich, M.M., 1998. Cortical plasticity: From synapses to maps. Annu. Rev. Neurosci. 21, 149–186. 10.1146/annurev.neuro.21.1.149.

Burke, J.F., Long, N.M., Zaghloul, K.A., Sharan, A.D., Sperling, M.R., Kahana, M.J., 2014. Human intracranial high-frequency activity maps episodic memory formation in space and time. NeuroImage 85, 834–843. 10.1016/j.neuroimage.2013.06.067.

Burkhardt, E., Zemmoura, I., Hirsch, F., Lemaitre, A.L., Deverdun, J., Moritz-Gasser, S., Duffau, H., Herbet, G., 2023. The central role of the left inferior longitudinal fasciculus in the face-name retrieval network. Hum. Brain. Mapp. 44, 3254–3270. 10.1002/hbm.26279.

Carl, C., Açık, A., König, P., Engel, A.K., Hipp, J.F., 2012. The saccadic spike artifact in MEG. Neuroimage. 59, 1657–1667. 10.1016/j.neuroimage.2011.09.020.

Catani, M., Jones, D.K., Ffytche, D.H., 2005. Perisylvian language networks of the human brain. Ann. Neurol. 57, 8–16. 10.1002/ana.20319.

Chang, E.F., Rieger, J.W., Johnson, K., Berger, M.S., Barbaro, N.M., Knight, R.T., 2010. Categorical speech representation in human superior temporal gyrus. Nat. Neurosci. 13, 1428–1432. 10.1038/nn.2641.

Chang, E.F., Niziolek, C.A., Knight, R.T., Nagarajan, S.S., Houde, J.F., 2013. Human cortical sensorimotor network underlying feedback control of vocal pitch. Proc. Natl. Acad. Sci. U. S. A. 110, 2653–2658. 10.1073/pnas.1216827110.

Chang, E.F., Raygor, K.P., Berger, M.S., 2015. Contemporary model of language organization: An overview for neurosurgeons. J. Neurosurg. 122, 250–261. 10.3171/2014.10.JNS132647.

Chernoff, B.L., Teghipco, A., Garcea, F.E., Sims, M.H., Paul, D.A., Tivarus, M.E., Smith, S. O., Pilcher, W.H., Mahon, B.Z., 2018. A role for the frontal aslant tract in speech planning: A neurosurgical case study. J. Cogn. Neurosci. 30, 752–769. 10.1162/jocn_a_01244.

Chidambaram, S., Anthony, D., Jansen, T., Vigo, V., Fernandez Miranda, J.C., 2023. Intraoperative augmented reality fiber tractography complements cortical and subcortical mapping. World. Neurosurg. X. 20, 100226. 10.1016/j.wnsx.2023.100226.

Cogan, G.B., Thesen, T., Carlson, C., Doyle, W., Devinsky, O., Pesaran, B., 2014. Sensory-motor transformations for speech occur bilaterally. Nature. 507, 94–98. 10.1038/nature12935.

Crossley, N.A., Mechelli, A., Vértes, P.E., Winton-Brown, T.T., Patel, A.X., Ginestet, C.E., McGuire, P., Bullmore, E.T., 2013. Cognitive relevance of the community structure of the human brain functional coactivation network. Proc. Natl. Acad. Sci. U. S. A. 110, 11583–11588. 10.1073/pnas.1220826110.

Crutchfield, J.S., Sawaya, R., Meyers, C.A., Moore, B.D., 1994. Postoperative mutism in neurosurgery. Report of two cases. J. Neurosurg. 81, 115–121. 10.3171/jns.1994.81.1.0115.

De Witt Hamer, P.C., Moritz-Gasser, S., Gatignol, P., Duffau, H., 2011. Is the human left middle longitudinal fascicle essential for language? A brain electrostimulation study. Hum. Brain. Mapp. 32, 962–973. 10.1002/hbm.21082.

Desikan, R.S., Ségonne, F., Fischl, B., Quinn, B.T., Dickerson, B.C., Blacker, D., Buckner, R. L., Dale, A.M., Maguire, R.P., Hyman, B.T., Albert, M.S., Killiany, R.J., 2006. An automated labeling system for subdividing the human cerebral cortex on MRI scans into gyral based regions of interest. Neuroimage. 31, 968–980. 10.1016/j.neuroimage.2006.01.021.

Dick, A.S., Tremblay, P., 2012. Beyond the arcuate fasciculus: Consensus and controversy in the connectional anatomy of language. Brain. 135, 3529–3550. 10.1093/brain/aws222.

Duffau, H., Capelle, L., Sichez, N., Denvil, D., Lopes, M., Sichez, J.P., Bitar, A., Fohanno, D., 2002. Intraoperative mapping of the subcortical language pathways using direct stimulations. An anatomo-functional study. Brain. 125, 199–214. 10.1093/brain/awf016.

Duffau, H., Gatignol, P., Moritz-Gasser, S., Mandonnet, E., 2009. Is the left uncinate fasciculus essential for language? A cerebral stimulation study. J. Neurol. 256, 382–389. 10.1007/s00415-009-0053-9.

Fedio, P., Ommaya, A.K., 1970. Bilateral cingulum lesions and stimulation in man with lateralized impairment in short-term verbal memory. Exp. Neurol. 29, 84–91. 10.1016/0014-4886(70)90039-7.

Flinker, A., Korzeniewska, A., Shestyuk, A.Y., Franaszczuk, P.J., Dronkers, N.F., Knight, R. T., Crone, N.E., 2015. Redefining the role of Broca’s area in speech. Proc. Natl. Acad. Sci. U. S. A. 112, 2871–2875. 10.1073/pnas.1414491112.

Floegel, M., Fuchs, S., Kell, C.A., 2020. Differential contributions of the two cerebral hemispheres to temporal and spectral speech feedback control. Nat. Commun. 11, 2839. 10.1038/s41467-020-16743-2.

Frauscher, B., von Ellenrieder, N., Zelmann, R., Doležalová, I., Minotti, L., Olivier, A., Hall, J., Hoffmann, D., Nguyen, D. K., Kahane, P., Dubeau, F., Gotman, J., 2018. Atlas of the normal intracranial electroencephalogram: Neurophysiological awake activity in different cortical areas. Brain. 141, 1130–1144. 10.1093/brain/awy035.

Fridriksson, J., Guo, D., Fillmore, P., Holland, A., Rorden, C., 2013. Damage to the anterior arcuate fasciculus predicts non-fluent speech production in aphasia. Brain. 136, 3451–3460. 10.1093/brain/awt267.

Friederici, A.D., Gierhan, S.M.E., 2013. The language network. Curr. Opin. Neurobiol. 23, 250–254. 10.1016/j.conb.2012.10.002.

Gajardo-Vidal, A., Lorca-Puls, D.L., Ploras Team, Warner, H., Pshdary, B., Crinion, J.T., Leff, A.P., Hope, T.M.H., Geva, S., Seghier, M.L., Green, D.W., Bowman, H., Price, C.J., 2021. Damage to Broca’s area does not contribute to long-term speech production outcome after stroke. Brain. 144, 817–832. 10.1093/brain/awaa460.

Gathercole, S.E., Willis, C.S., Baddeley, A.D., Emslie, H., 1994. The children’s test of nonword repetition: A test of phonological working memory. Memory. 2, 103–127. 10.1080/09658219408258940.

Ghosh, S.S., Kakunoori, S., Augustinack, J., Nieto-Castanon, A., Kovelman, I., Gaab, N., Christodoulou, J.A., Triantafyllou, C., Gabrieli, J.D.E., Fischl, B., 2010. Evaluating the validity of volume-based and surface-based brain image registration for developmental cognitive neuroscience studies in children 4 to 11 years of age. Neuroimage. 53, 85–93. 10.1016/j.neuroimage.2010.05.075.

Giampiccolo, D., Duffau, H., 2022. Controversy over the temporal cortical terminations of the left arcuate fasciculus: A reappraisal. Brain. 145, 1242–1256. 10.1093/brain/awac057.

Glasser, M.F., Smith, S.M., Marcus, D.S., Andersson, J.L.R., Auerbach, E.J., Behrens, T.E. J., Coalson, T.S., Harms, M.P., Jenkinson, M., Moeller, S., Robinson, E.C., Sotiropoulos, S.N., Xu, J., Yacoub, E., Ugurbil, K., Van Essen, D.C., 2016. The Human Connectome Project’s neuroimaging approach. Nat. Neurosci. 19, 1175–1187. 10.1038/nn.4361.

Greenlee, J.D.W., Behroozmand, R., Larson, C.R., Jackson, A.W., Chen, F., Hansen, D.R., Oya, H., Kawasaki, H., Howard, M. A., 2013. Sensory-motor interactions for vocal pitch monitoring in non-primary human auditory cortex. PLoS One. 8, e60783. 10.1371/journal.pone.0060783.

Griffiths, J.D., Marslen-Wilson, W.D., Stamatakis, E.A., Tyler, L.K., 2013. Functional organization of the neural language system: Dorsal and ventral pathways are critical for syntax. Cereb. Cortex. 23, 139–147. 10.1093/cercor/bhr386.

Hamberger, M.J., 2007. Cortical language mapping in epilepsy: A critical review. Neuropsychol. Rev. 17, 477–489. 10.1007/s11065-007-9046-6.

Hansen, C.B., Yang, Q., Lyu, I., Rheault, F., Kerley, C., Chandio, B.Q., Fadnavis, S., Williams, O., Shafer, A.T., Resnick, S.M., Zald, D.H., Cutting, L.E., Taylor, W.D., Boyd, B., Garyfallidis, E., Anderson, A.W., Descoteaux, M., Landman, B.A., Schilling, K.G., 2021. Pandora: 4-D white matter bundle population-based atlases derived from diffusion MRI fiber tractography. Neuroinformatics. 19, 447–460. 10.1007/s12021-020-09497-1.

Harvey, D.Y., Wei, T., Ellmore, T.M., Hamilton, A.C., Schnur, T.T., 2013. Neuropsychological evidence for the functional role of the uncinate fasciculus in semantic control. Neuropsychologia. 51, 789–801. 10.1016/j.neuropsychologia.2013.01.028.

Harvey, D.Y., Schnur, T.T., 2015. Distinct loci of lexical and semantic access deficits in aphasia: Evidence from voxel-based lesion-symptom mapping and diffusion tensor imaging. Cortex. 67, 37–58. 10.1016/j.cortex.2015.03.004.

Herbet, G., Moritz-Gasser, S., Boiseau, M., Duvaux, S., Cochereau, J., Duffau, H., 2016. Converging evidence for a cortico-subcortical network mediating lexical retrieval. Brain. 139, 3007–3021. 10.1093/brain/aww220.

Hervé, P.Y., Zago, L., Petit, L., Mazoyer, B., Tzourio-Mazoyer, N., 2013. Revisiting human hemispheric specialization with neuroimaging. Trends. Cogn. Sci. 17, 69–80. 10.1016/j.tics.2012.12.004.

Hickok, G., Poeppel, D., 2000. Towards a functional neuroanatomy of speech perception. Trends. Cogn. Sci. 4, 131–138. 10.1016/S1364-6613(00)01463-7.

Hoechstetter, K., Bornfleth, H., Weckesser, D., Ille, N., Berg, P., Scherg, M., 2004. BESA source coherence: A new method to study cortical oscillatory coupling. Brain. Topogr. 16, 233–238. 10.1023/B:BRAT.0000032857.55223.5d.

Hoptman, M.J., Davidson, R.J., 1994. How and why do the two cerebral hemispheres interact? Psychol. Bull. 116, 195–219. 10.1037/0033-2909.116.2.195.

Janssen, N., Kessels, R.P.C., Mars, R.B., Llera, A., Beckmann, C.F., Roelofs, A., 2023. Dissociating the functional roles of arcuate fasciculus subtracks in speech production. Cereb. Cortex. 33, 2539–2547. 10.1093/cercor/bhac224.

Jenkinson, M., Beckmann, C.F., Behrens, T.E.J., Woolrich, M.W., Smith, S.M., 2012. FSL. Neuroimage. 62, 782–790. 10.1016/j.neuroimage.2011.09.015.

Kambara, T., Brown, E.C., Jeong, J.W., Ofen, N., Nakai, Y., Asano, E., 2017. Spatio-temporal dynamics of working memory maintenance and scanning of verbal information. Clin. Neurophysiol. 128, 882–891. 10.1016/j.clinph.2017.03.005.

Kambara, T., Sood, S., Alqatan, Z., Klingert, C., Ratnam, D., Hayakawa, A., Nakai, Y., Luat, A.F., Agarwal, R., Rothermel, R., Asano, E., 2018. Presurgical language mapping using event-related high-gamma activity: The Detroit procedure. Clin. Neurophysiol. 129, 145–154. 10.1016/j.clinph.2017.10.018.

Katzev, M., Tüscher, O., Hennig, J., Weiller, C., Kaller, C.P., 2013. Revisiting the functional specialization of left inferior frontal gyrus in phonological and semantic fluency: The crucial role of task demands and individual ability. J. Neurosci. 33, 7837–7845. 10.1523/JNEUROSCI.3147-12.2013.

Kinoshita, M., Menjot de Champfleur, N., Deverdun, J., Moritz-Gasser, S., Herbet, G., Duffau, H., 2015. Role of fronto-striatal tract and frontal aslant tract in movement and speech: An axonal mapping study. Brain. Struct. Funct. 220, 3399–3412. 10.1007/s00429-014-0863-0.

Kitazawa, Y., Sonoda, M., Sakakura, K., Mitsuhashi, T., Firestone, E., Ueda, R., Kambara, T., Iwaki, H., Luat, A.F., Marupudi, N.I., Sood, S., Asano, E., 2023. Intra- and inter-hemispheric network dynamics supporting object recognition and speech production. Neuroimage. 270, 119954. 10.1016/j.neuroimage.2023.119954.

Kunii, N., Kamada, K., Ota, T., Greenblatt, R.E., Kawai, K., Saito, N., 2013. The dynamics of language-related high-gamma activity assessed on a spatially normalized brain. Clin. Neurophysiol. 124, 91–100. 10.1016/j.clinph.2012.06.006.

Kural, M.A., Duez, L., Hansen, V.S., Larsson, P.G., Rampp, S., Schulz, R., Tankisi, H., Wennberg, R., Bibby, B.M., Scherg, M., Beniczky, S., 2020. Criteria for defining interictal epileptiform discharges in EEG: A clinical validation study. Neurology. 94, e2139–e2147. 10.1212/WNL.0000000000009439.

Kuroda, N., Sonoda, M., Miyakoshi, M., Nariai, H., Jeong, J.W., Motoi, H., Luat, A.F., Sood, S., Asano, E., 2021. Objective interictal electrophysiology biomarkers optimize prediction of epilepsy surgery outcome. Brain. Commun. 3, fcab042. 10.1093/braincomms/fcab042.

Kwan, P., Brodie, M.J., 2001. Neuropsychological effects of epilepsy and antiepileptic drugs. Lancet. 357, 216–222. 10.1016/S0140-6736(00)03600-X.

Leszczyński, M., Barczak, A., Kajikawa, Y., Ulbert, I., Falchier, A. Y., Tal, I., Haegens, S., Melloni, L., Knight, R.T., Schroeder, C.E., 2020. Dissociation of broadband high-frequency activity and neuronal firing in the neocortex. Sci. Adv. 6, eabb0977. 10.1126/sciadv.abb0977.

Levelt, W.J.M., 1999. Models of word production. Trends Cogn. Sci. 3, 223–232. 10.1016/S1364-6613(99)01319-4.

Liberman, A. M., Whalen, D. H., 2000. On the relation of speech to language. Trends. Cogn. Sci. 4, 187–196. 10.1016/S1364-6613(00)01471-6.

Liebenthal, E., Binder, J.R., Spitzer, S.M., Possing, E.T., Medler, D.A., 2005. Neural substrates of phonemic perception. Cereb. Cortex, 15, 1621–1631. 10.1093/cercor/bhi040.

Lu, J., Zhao, Z., Zhang, J., Wu, B., Zhu, Y., Chang, E.F., Wu, J., Duffau, H., Berger, M.S., 2021. Functional maps of direct electrical stimulation-induced speech arrest and anomia: A multicentre retrospective study. Brain. 144, 2541–2553. 10.1093/brain/awab125.

Luo, C., Makaretz, S., Stepanovic, M., Papadimitriou, G., Quimby, M., Palanivelu, S., Dickerson, B.C., Makris, N., 2020. Middle longitudinal fascicle is associated with semantic processing deficits in primary progressive aphasia. Neuroimage. Clin. 25, 102115. 10.1016/j.nicl.2019.102115.

Maldonado, I.L., Moritz-Gasser, S., Duffau, H., 2011. Does the left superior longitudinal fascicle subserve language semantics? A brain electrostimulation study. Brain. Struct. Funct. 216, 263–274. 10.1007/s00429-011-0309-x.

Mandonnet, E., Nouet, A., Gatignol, P., Capelle, L., Duffau, H., 2007. Does the left inferior longitudinal fasciculus play a role in language? A brain stimulation study. Brain. 130, 623–629. 10.1093/brain/awl361.

Marchina, S., Zhu, L.L., Norton, A., Zipse, L., Wan, C.Y., Schlaug, G., 2011. Impairment of speech production predicted by lesion load of the left arcuate fasciculus. Stroke. 42, 2251–2256. 10.1161/STROKEAHA.110.606103.

Martinez Oeckel, A., Rijntjes, M., Glauche, V., Kümmerer, D., Kaller, C. P., Egger, K., Weiller, C., 2021. The extreme capsule and aphasia: Proof-of-concept of a new way relating structure to neurological symptoms. Brain. Commun. 3, fcab040. 10.1093/braincomms/fcab040.

Mazoyer, B., Zago, L., Jobard, G., Crivello, F., Joliot, M., Perchey, G., Mellet, E., Petit, L., Tzourio-Mazoyer, N., 2014. Gaussian mixture modeling of hemispheric lateralization for language in a large sample of healthy individuals balanced for handedness. PLoS One. 9, e101165. 10.1371/journal.pone.0101165.

Mercier, M.R., Dubarry, A.S., Tadel, F., Avanzini, P., Axmacher, N., Cellier, D., Del Vecchio, M., Hamilton, L.S., Hermes, D., Kahana, M.J., Knight, R.T., Llorens, A., Megevand, P., Melloni, L., Miller, K.J., Piai, V., Puce, A., Ramsey, N.F., Schwiedrzik, C. M., Smith, S.E., Stolk, A., Swann, N.C., Vansteensel, M.J., Voytek, B., Wang, L., Lachaux, J.P., Oostenveld, R., 2022. Advances in human intracranial electroencephalography research, guidelines and good practices. Neuroimage. 260, 119438. 10.1016/j.neuroimage.2022.119438.

Möddel, G., Lineweaver, T., Schuele, S.U., Reinholz, J., Loddenkemper, T., 2009. Atypical language lateralization in epilepsy patients. Epilepsia. 50, 1505–1516. 10.1111/j.1528-1167.2008.02000.x.

Nagasawa, T., Matsuzaki, N., Juhász, C., Hanazawa, A., Shah, A., Mittal, S., Sood, S., Asano, E., 2011. Occipital gamma-oscillations modulated during eye movement tasks: simultaneous eye tracking and electrocorticography recording in epileptic patients. Neuroimage 58, 1101–1109. 10.1016/j.neuroimage.2011.06.085.

Nakai, Y., Jeong, J.W., Brown, E. C., Rothermel, R., Kojima, K., Kambara, T., Shah, A., Mittal, S., Sood, S., Asano, E., 2017. Three- and four-dimensional mapping of speech and language in patients with epilepsy. Brain. 140, 1351–1370. 10.1093/brain/awx051.

Nishida, M., Juhász, C., Sood, S., Chugani, H.T., Asano, E., 2008. Cortical glucose metabolism positively correlates with gamma-oscillations in nonlesional focal epilepsy. Neuroimage. 42, 1275–1284. 10.1016/j.neuroimage.2008.06.027.

Nishida, M., Korzeniewska, A., Crone, N.E., Toyoda, G., Nakai, Y., Ofen, N., Brown, E.C., Asano, E., 2017. Brain network dynamics in the human articulatory loop. Clin. Neurophysiol. 128, 1473–1487. 10.1016/j.clinph.2017.05.002.

Norman, D.A., Bobrow, D.G., 1975. On data-limited and resource-limited processes. Cogn. Psychol. 7, 44–64. 10.1016/0010-0285(75)90004-3.

Ono, H., Sonoda, M., Sakakura, K., Kitazawa, Y., Mitsuhashi, T., Firestone, E., Jeong, J.W., Luat, A.F., Marupudi, N.I., Sood, S., Asano, E., 2023. Dynamic cortical and tractography atlases of proactive and reactive alpha and high-gamma activities. Brain. Commun. 5, fcad111. 10.1093/braincomms/fcad111.

Ortibus, E., Verhoeven, J., Sunaert, S., Casteels, I., de Cock, P., Lagae, L., 2012. Integrity of the inferior longitudinal fasciculus and impaired object recognition in children: A diffusion tensor imaging study. Dev. Med. Child. Neurol. 54, 38–43. 10.1111/j.1469-8749.2011.04147.x.

Papagno, C., Gallucci, M., Casarotti, A., Castellano, A., Falini, A., Fava, E., Giussani, C., Carrabba, G., Bello, L., Caramazza, A., 2011. Connectivity constraints on cortical reorganization of neural circuits involved in object naming. Neuroimage. 55, 1306–1313. 10.1016/j.neuroimage.2011.01.005.

Papp, N., Ktonas, P., 1977. Critical evaluation of complex demodulation techniques for the quantification of bioelectrical activity. Biomed. Sci. Instrum. 13, 135–145.

Perrone-Bertolotti, M., Alexandre, S., Jobb, A.S., De Palma, L., Baciu, M., Mairesse, M.P., Hoffmann, D., Minotti, L., Kahane, P., David, O., 2020. Probabilistic mapping of language networks from high frequency activity induced by direct electrical stimulation. Hum. Brain. Mapp. 41, 4113–4126. 10.1002/hbm.25112.

Lambon Ralph, M.A., Jefferies, E., Patterson, K., Rogers, T.T., 2017. The neural and computational bases of semantic cognition. Nat. Rev. Neurosci. 18, 42–55. 10.1038/nrn.2016.150.

Ray, S., Crone, N.E., Niebur, E., Franaszczuk, P.J., Hsiao, S.S., 2008. Neural correlates of high-gamma oscillations (60-200 Hz) in macaque local field potentials and their potential implications in electrocorticography. J. Neurosci. 28, 11526–11536. 10.1523/JNEUROSCI.2848-08.2008.

Riva, M., Wilson, S.M., Cai, R., Castellano, A., Jordan, K.M., Henry, R.G., Gorno Tempini, M.L., Berger, M.S., Chang, E.F., 2022. Evaluating syntactic comprehension during awake intraoperative cortical stimulation mapping. J. Neurosurg. 138, 1403–1410. 10.3171/2022.8.JNS221335.

Sakakura, K., Kuroda, N., Sonoda, M., Mitsuhashi, T., Firestone, E., Luat, A.F., Marupudi, N.I., Sood, S., Asano, E., 2023. Developmental atlas of phase-amplitude coupling between physiologic high-frequency oscillations and slow waves. Nat. Commun. 14, 6435. 10.1038/s41467-023-42091-y.

Sammler, D., Cunitz, K., Gierhan, S.M.E., Anwander, A., Adermann, J., Meixensberger, J., Friederici, A.D., 2018. White matter pathways for prosodic structure building: A case study. Brain. Lang. 183, 1–10. 10.1016/j.bandl.2018.05.001.

Scheeringa, R., Fries, P., Petersson, K.M., Oostenveld, R., Grothe, I., Norris, D.G., Hagoort, P., Bastiaansen, M.C.M., 2011. Neuronal dynamics underlying high- and low-frequency EEG oscillations contribute independently to the human BOLD signal. Neuron, 69, 572–583. 10.1016/j.neuron.2010.11.044.

Shuren, J.E., Schefft, B.K., Yeh, H.S., Privitera, M.D., Cahill, W.T., Houston, W., 1995. Repetition and the arcuate fasciculus. J. Neurol. 242, 596–598. 10.1007/BF00868813.

Singer, W., 1993. Synchronization of cortical activity and its putative role in information processing and learning. Annu. Rev. Physiol. 55, 349–374. 10.1146/annurev.ph.55.030193.002025.

Skeide, M.A., Friederici, A.D., 2016. The ontogeny of the cortical language network. Nat. Rev. Neurosci. 17, 323–332. 10.1038/nrn.2016.23.

Solomon, E.A., Kragel, J.E., Sperling, M.R., Sharan, A., Worrell, G., Kucewicz, M., Inman, C.S., Lega, B., Davis, K.A., Stein, J.M., Jobst, B.C., Zaghloul, K.A., Sheth, S.A., Rizzuto, D.S., Kahana, M.J., 2017. Widespread theta synchrony and high-frequency desynchronization underlie enhanced cognition. Nat. Commun. 8, 1704. 10.1038/s41467-018-06876-w.

Sonoda, M., Silverstein, B.H., Jeong, J.W., Sugiura, A., Nakai, Y., Mitsuhashi, T., Rothermel, R., Luat, A.F., Sood, S., Asano, E., 2021. Six-dimensional dynamic tractography atlas of language connectivity in the developing brain. Brain. 144, 3340 – 3354. 10.1093/brain/awab225.

Sonoda, M., Rothermel, R., Carlson, A., Jeong, J.W., Lee, M.H., Hayashi, T., Luat, A.F., Sood, S., Asano, E., 2022. Naming-related spectral responses predict neuropsychological outcome after epilepsy surgery. Brain. 145, 517–530. 10.1093/brain/awab318.

Stolk, A., Griffin, S., van der Meij, R., Dewar, C., Saez, I., Lin, J.J., Piantoni, G., Schoffelen, J.M., Knight, R.T., Oostenveld, R., 2018. Integrated analysis of anatomical and electrophysiological human intracranial data. Nat. Protoc. 13, 1699–1723. 10.1038/s41596-018-0009-6.

Sun, F.T., Miller, L.M., D’Esposito, M., 2004. Measuring interregional functional connectivity using coherence and partial coherence analyses of fMRI data. Neuroimage. 21, 647–658. 10.1016/j.neuroimage.2003.09.056.

Sussman, N.M., Gur, R.C., Gur, R.E., O’Connor, M.J., 1983. Mutism as a consequence of callosotomy. J. Neurosurg. 59, 514–519. 10.3171/jns.1983.59.3.0514.

Szaflarski, J.P., Binder, J.R., Possing, E.T., McKiernan, K.A., Ward, B.D., Hammeke, T.A., 2002. Language lateralization in left-handed and ambidextrous people: fMRI data. Neurology. 59, 238–244. 10.1212/wnl.59.2.238.

Szaflarski, J.P., Holland, S.K., Schmithorst, V.J., Byars, A.W., 2006. fMRI study of language lateralization in children and adults. Hum. Brain. Mapp. 27, 202–212. 10.1002/hbm.20177.

Tate, M.C., Herbet, G., Moritz-Gasser, S., Tate, J.E., Duffau, H., 2014. Probabilistic map of critical functional regions of the human cerebral cortex: Broca’s area revisited. Brain. 137, 2773–2782. 10.1093/brain/awu168.

Taylor, P.N., Papasavvas, C.A., Owen, T.W., Schroeder, G.M., Hutchings, F.E., Chowdhury, F.A., Diehl, B., Duncan, J.S., McEvoy, A.W., Miserocchi, A., de Tisi, J., Vos, S.B., Walker, M.C., Wang, Y., 2022. Normative brain mapping of interictal intracranial EEG to localize epileptogenic tissue. Brain. 145, 939–949. 10.1093/brain/awab380.

Thiebaut de Schotten, M., Ffytche, D.H., Bizzi, A., Dell’Acqua, F., Allin, M., Walshe, M., Murray, R., Williams, S.C., Murphy, D.G.M., Catani, M., 2011. Atlasing location, asymmetry and inter-subject variability of white matter tracts in the human brain with MR diffusion tractography. Neuroimage. 54, 49–59. 10.1016/j.neuroimage.2010.07.055.

Tomasino, B., Tronchin, G., Marin, D., Maieron, M., Fabbro, F., Cubelli, R., Luzzatti, C., 2018. Noun–verb naming dissociation in neurosurgical patients. Aphasiology. 33, 1418–1440. 10.1080/02687038.2018.1542658.

Tremblay, P., Perron, M., Deschamps, I., Kennedy-Higgins, D., Houde, J.C., Dick, A.S., Descoteaux, M., 2019. The role of the arcuate and middle longitudinal fasciculi in speech perception in noise in adulthood. Hum. Brain. Mapp. 40, 226–241. 10.1002/hbm.24367.

Ueda, R., Sakakura, K., Mitsuhashi, T., Sonoda, M., Firestone, E., Kuroda, N., Kitazawa, Y., Uda, H., Luat, A.F., Johnson, E.L., Ofen, N., Asano, E., 2024. Cortical and white matter substrates supporting visuospatial working memory. Clin. Neurophysiol. 162, 9–27. 10.1016/j.clinph.2024.03.008.

Uematsu, M., Matsuzaki, N., Brown, E.C., Kojima, K., Asano, E., 2013. Human occipital cortices differentially exert saccadic suppression: Intracranial recording in children. Neuroimage 83, 224–236. 10.1016/j.neuroimage.2013.06.058.

Vassal, F., Boutet, C., Lemaire, J.J., Nuti, C., 2014. New insights into the functional significance of the frontal aslant tract: An anatomo-functional study using intraoperative electrical stimulations combined with diffusion tensor imaging-based fiber tracking. Br. J. Neurosurg. 28, 685–687. 10.3109/02688697.2014.889810.

Warren, J.E., Wise, R.J.S., Warren, J.D., 2005. Sounds do-able: Auditory-motor transformations and the posterior temporal plane. Trends. Neurosci. 28, 636–643. 10.1016/j.tins.2005.09.010.

Wilson, S.M., Galantucci, S., Tartaglia, M.C., Rising, K., Patterson, D.K., Henry, M.L., Ogar, J.M., DeLeon, J., Miller, B.L., Gorno-Tempini, M.L., 2011. Syntactic processing depends on dorsal language tracts. Neuron. 72, 397–403. 10.1016/j.neuron.2011.09.014.

Wise, R.J., Greene, J., Büchel, C., Scott, S. K., 1999. Brain regions involved in articulation. The Lancet, 353(9158), 1057–1061. 10.1016/S0140-6736(98)07491-1

Woolnough, O., Donos, C., Rollo, P.S., Forseth, K.J., Lakretz, Y., Crone, N.E., Fischer-Baum, S., Dehaene, S., Tandon, N., 2021. Spatiotemporal dynamics of orthographic and lexical processing in the ventral visual pathway. Nat. Hum. Behav. 5, 389–398. 10.1038/s41562-020-00982-w.

Yeh, F.C., 2022. Population-based tract-to-region connectome of the human brain and its hierarchical topology. Nat. Commun. 13, 4933. 10.1038/s41467-022-32595-4.

Yeh, F.C., Panesar, S., Fernandes, D., Meola, A., Yoshino, M., Fernandez-Miranda, J.C., Vettel, J.M., Verstynen, T., 2018. Population-averaged atlas of the macroscale human structural connectome and its network topology. Neuroimage, 178, 57–68. 10.1016/j.neuroimage.2018.05.027.

Yuval-Greenberg, S., Tomer, O., Keren, A.S., Nelken, I., Deouell, L.Y., 2008. Transient induced gamma-band response in EEG as a manifestation of miniature saccades. Neuron. 58, 429–441. 10.1016/j.neuron.2008.03.027.

Zatorre, R.J., Belin, P., 2001. Spectral and temporal processing in human auditory cortex. Cereb. Cortex. 11, 946–953. 10.1093/cercor/11.10.946.

Zhao, Z., Huang, C.C., Yuan, S., Zhang, J., Lin, C.P., Lu, J., Duffau, H., Wu, J., 2023. Convergence of the arcuate fasciculus and third branch of the superior longitudinal fasciculus with direct cortical stimulation-induced speech arrest area in the anterior ventral precentral gyrus. J. Neurosurg. 139, 1140–1151. 10.3171/2023.1.JNS222575.

Zheng, Z.Z., Munhall, K.G., Johnsrude, I.S., 2010. Functional overlap between regions involved in speech perception and in monitoring one’s own voice during speech production. J. Cogn. Neurosci. 22, 1770–1781. 10.1162/jocn.2009.21324.

Zhong, A.J., Baldo, J.V., Dronkers, N.F., Ivanova, M.V., 2022. The unique role of the frontal aslant tract in speech and language processing. Neuroimage. Clin. 34, 103020. 10.1016/j.nicl.2022.103020.

