## Supplemental tables and figures for "Dynamic Causal Tractography Analysis of Auditory Descriptive Naming: An Intracranial Study of 106 Patients"

**in**

**An Intracranial Study of 106 Patients**

This document includes:

**Supplementary Tables 1-2.**

**Supplementary Figures 1-5.**

**Legend for Supplementary Video 1.**

**Results from Ancillary Analyses.**

| **Patient** | **Age at** | **Sex** |  |  |  |  |  |  | **Median** |
| --- | --- | --- | --- | --- | --- | --- | --- | --- | --- |
|  | **surgery** |  | **Handed**  **ness** | **MRI** | **Sampled**  **hemisphere** | **Resected**  **site** | **SOZ** | **Number of**  **antiseizure medications** | **response time** |
|  | **(years)** |  |  |  |  |  |  |  | **(seconds)** |
| 1 | 16 | M | Rt | Tumor | Lt | Lt | Lt F | 3 (LEV, OXC, CBZ) | 0.751 |
| 2 | 17 | F | Rt | Nonlesion | Rt | Rt | Rt O | 2 (OXC, LEV) | 1.001 |
| 3 | 15 | F | Rt | Tumor | Rt | Rt | NA | 1 (OXC) | 0.827 |
| 4 | 14 | M | Rt | Tumor | Lt | Lt | NA | 1 (OXC) | 0.668 |
| 5 | 16 | F | Rt | Nonlesion | Lt and Rt | Rt | Rt TPO | 3 (CZP, TPM, PHT) | 1.450 |
| 6 | 17 | F | Rt | Nonlesion | Lt and Rt | Lt | Lt F | 1 (LTG) | 0.883 |
| 7 | 10 | M | Rt | Dysplasia | Rt | Rt | Rt O | 3 (TPM, LEV, OXC) | 0.949 |
| 8 | 8 | F | Rt | Nonlesion | Rt | Rt | NA | 2 (LTG, OXC) | 1.306 |
| 9 | 8 | M | Rt | Tumor | Lt and Rt | Lt | NA | 1 (OXC) | 2.158 |
| 10 | 11 | F | Rt | Nonlesion | Rt | Rt | Rt T | 2 (OXC, LEV) | 1.339 |
| 11 | 16 | M | Rt | Nonlesion | Rt | Rt | Rt T | 1 (OXC) | 1.799 |
| 12 | 18 | F | Rt | Dysplasia | Lt and Rt | Rt | Rt PO | 3 (ZNS, OXC, LTG) | 0.664 |
| 13 | 17 | M | Rt | Tumor | Lt | Lt | Lt T | 1 (OXC) | 0.620 |
| 14 | 8 | M | Lt | Tumor | Lt | Lt | NA | 0 | 1.979 |
| 15 | 17 | M | Rt | Nonlesion | Lt | Lt | NA | 2 (OXC, LEV) | 0.684 |
| 16 | 14 | M | Rt | Nonlesion | Lt | Lt | Lt T | 1 (VPA) | 2.285 |
| 17 | 10 | F | Rt | Tumor | Lt | Lt | Lt TP | 2 (OXC, TPM) | 0.719 |
| 18 | 10 | M | Lt | Nonlesion | Lt and Rt | NA | Lt FT, Rt F | 2 (VPA, CBZ) | 1.265 |
| 19 | 15 | M | Rt | Tumor | Lt | Lt | Lt T | 2 (LEV, LTG) | 1.155 |
| 20 | 19 | F | Rt | Nonlesion | Lt and Rt | Rt | Rt P | 3 (OXC, LTG, LCM) | 0.762 |
| 21 | 8 | M | Rt | Dysplasia | Lt and Rt | Rt | Rt FTPO | 2 (OXC, LTG) | 1.055 |
| 22 | 14 | F | Rt | Nonlesion | Lt | Lt | Lt T | 3 (OXC, LTG, TPM) | 1.285 |
| 23 | 11 | M | Rt | Dysplasia | Rt | Rt* | Rt FP | 2 (OXC, LEV) | 0.808 |
| 24 | 13 | M | Rt | Tumor | Rt | Rt | Rt FP | 1 (OXC) | 0.627 |
| 25 | 10 | M | Rt | Tumor | Rt | Rt | Rt T | 1 (OXC) | 1.220 |
| 26 | 5 | M | Rt | Nonlesion | Lt | Lt | Lt F | 3 (OXC, LEV, VPA) | 1.831 |
| 27 | 16 | F | Rt | Nonlesion | Lt and Rt | Lt | Lt PO | 2 (LEV, OXC) | 0.904 |
| 28 | 16 | M | Rt | Nonlesion | Rt | Rt | Rt TPO | 2 (LEV, OXC) | 0.983 |
| 29 | 37 | F | Rt | Tumor | Lt | Lt | NA | 1 (LEV) | 0.832 |
| 30 | 11 | F | Rt | Nonlesion | Rt | Rt | Rt FP | 2 (LEV, LTG) | 2.290 |
| 31 | 21 | F | Lt | Tumor | Lt | Lt | Lt O | 1 (LEV) | 0.446 |
| 32 | 15 | F | Rt | Tumor | Lt | Lt | Lt T | 1 (LEV) | 1.301 |
| 33 | 5 | F | Rt | Others | Lt | Lt | Lt FTPO | 5 (LCM, LEV, OXC, ZNS, CZP) | 2.178 |
| 34 | 14 | F | Rt | Dysplasia | Lt | Lt | NA | 3 (LEV, OXC, LCM) | 1.099 |
| 35 | 28 | F | Rt | Nonlesion | Lt | Lt | Lt T | 2 (CBZ, LCM) | 2.563 |
| 36 | 14 | F | Rt | Others | Lt and Rt | Rt | Rt T | 2 (LEV, TPM) | 1.425 |
| 37 | 13 | F | Rt | Nonlesion | Rt | Rt | NA | 3 (LEV, LTG, OXC) | 2.958 |
| 38 | 41 | F | Rt | Nonlesion | Lt | Lt | NA | 2 (LEV, PHT) | 0.736 |
| 39 | 12 | M | Rt | Nonlesion | Lt | Lt | Lt T | 3 (LCM, OXC, VPA) | 2.289 |
| 40 | 8 | M | Rt | Nonlesion | Lt and Rt | Rt | Rt F | 1 (LCM) | 1.329 |
| 41 | 10 | M | Rt | Others | Rt | Rt | NA | 1 (OXC) | 1.058 |
| 42 | 12 | M | Rt | Nonlesion | Rt | Rt | Rt T | 2 (VPA, LCM) | 1.870 |
| 43 | 28 | M | Rt | Tumor | Rt | Rt | Rt T | 1 (CBZ) | 0.523 |
| 44 | 27 | F | Rt | Tumor | Lt and Rt | Lt | Lt F | 2 (LEV, LCM) | 1.023 |
| 45 | 17 | M | Rt | Others | Rt | Rt | Rt T | 2 (LEV, LCM) | 0.900 |
| 46 | 14 | M | Rt | Nonlesion | Lt and Rt | Rt | NA | 3 (OXC, LTG, CLB) | 1.127 |
| 47 | 15 | F | Rt | Nonlesion | Lt and Rt | Rt | Rt T | 3 (LTG, LEV, ZNS) | 1.629 |
| 48 | 6 | F | Lt | Others | Lt and Rt | Lt | Lt T | 2 (VPA, LTG) | 1.321 |
| 49 | 17 | M | Rt | Dysplasia | Rt | Rt | Rt PI | 2 (OXC, TPM) | 1.023 |
| 50 | 4 | M | Rt | Dysplasia | Lt and Rt | Rt | Rt T | 2 (LCM, OXC) | 1.825 |
| 51 | 12 | F | Rt | Others | Lt | Lt | Lt T | 2 (LTG, LCM) | 0.914 |
| 52 | 5 | F | Rt | Others | Lt | Lt | Lt T | 2 (LEV, LCM) | 1.996 |
| 53 | 9 | F | Rt | Tumor | Lt | Lt | Lt T | 1 (CBZ) | 1.508 |
| 54 | 30 | M | Rt | Nonlesion | Lt and Rt | Rt | Rt T | 3 (PHT, CBZ, LCM) | 1.360 |
| 55 | 13 | M | Rt | Nonlesion | Rt | Rt | NA | 1 (LTG) | 0.718 |
| 56 | 17 | M | Lt | Nonlesion | Rt | Rt | Rt O | 2 (LTG, LCM) | 0.544 |
| 57 | 16 | M | Rt | Nonlesion | Lt | Lt | Lt F | 2 (LTG, OXC) | 1.016 |
| 58 | 8 | M | Rt | Tumor | Rt | Rt | Rt TO | 1 (OXC) | 2.273 |
| 59 | 13 | F | Rt | Nonlesion | Lt | Lt | Lt T | 1 (LEV) | 1.167 |
| 60 | 14 | F | Rt | Nonlesion | Rt | Rt | Rt P | 2 (LCM, LEV) | 1.164 |
| 61 | 12 | M | Rt | Nonlesion | Lt | Lt | Lt T | 1 (VPA) | 2.217 |
| 62 | 13 | F | Rt | Dysplasia | Rt | Rt | Rt T | 4 (LEV, CLB, PHT, LTG) | 2.732 |
| 63 | 11 | F | Rt | Tumor | Lt | Lt | NA | 2 (OXC, LEV) | 1.384 |
| 64 | 15 | M | Rt | Nonlesion | Rt | Rt | Rt F | 2 (LCM, VPA) | 1.688 |
| 65 | 14 | F | Rt | Dysplasia | Lt | Lt | Lt T | 2 (OXC, VPA) | 3.067 |
| 66 | 11 | F | Rt | Nonlesion | Rt | Rt | Rt P | 2 (LEV, OXC) | 1.127 |
| 67 | 12 | F | Rt | Tumor | Rt | Rt | Rt P | 1 (LEV) | 1.152 |
| 68 | 17 | M | Lt | Nonlesion | Lt | Lt | Lt F | 2 (OXC, LTG) | 1.048 |
| 69 | 11 | F | Rt | Nonlesion | Rt | Rt | Rt F | 2 (OXC, LEV) | 1.860 |
| 70 | 17 | M | Rt | Nonlesion | Lt | Lt | Lt T | 3 (OXC, LEV, LCM) | 0.780 |
| 71 | 10 | F | Rt | Dysplasia | Rt | Rt | NA | 3 (LEV, LCM, ZNS) | 1.762 |
| 72 | 13 | M | Rt | Dysplasia | Rt | Rt | Rt TPO | 3 (LEV, LCM, PHT) | 2.359 |
| 73 | 11 | M | Rt | Dysplasia | Rt | Rt | Rt P | 1 (LEV) | 1.964 |
| 74 | 16 | M | Rt | Dysplasia | Lt | Lt | Lt TO | 3 (OXC, LEV, LCM) | 1.833 |
| 75 | 15 | M | Rt | Nonlesion | Lt and Rt | Lt | Lt T | 1 (VPA) | 1.583 |
| 76 | 15 | M | Rt | Nonlesion | Lt | Lt | Lt T | 1 (VPA, LCM) | 1.129 |
| 77 | 10 | M | Rt | Nonlesion | Rt | Rt | Rt P | 1 (LCM) | 0.987 |
| 78 | 10 | M | Rt | Others | Rt | Rt | Rt T | 2 (OXC, LTG) | 1.068 |
| 79 | 16 | M | Rt | Nonlesion | Lt and Rt | NA | NA | 1 (OXC) | 0.724 |
| 80 | 12 | F | Rt | Dysplasia | Rt | Rt | Rt T | 2 (OXC, ZNS) | 1.487 |
| 81 | 8 | M | Rt | Tumor | Lt | Lt | Lt T | 2 (VPA, LEV) | 1.282 |
| 82 | 14 | F | Rt | Dysplasia | Rt | Rt | Rt T | 2 (OXC, LCM) | 2.191 |
| 83 | 19 | M | Rt | Dysplasia | Rt | Rt | Rt O | 2 (LEV, OXC) | 1.635 |
| 84 | 6 | F | Rt | Nonlesion | Lt | Lt | Lt PO | 2 (OXC, LCM) | 1.332 |
| 85 | 8 | M | Rt | Nonlesion | Rt | Rt | NA | 2 (LTG, LCM) | 1.807 |
| 86 | 16 | M | Rt | Nonlesion | Rt | Rt | Rt F | 1 (OXC) | 2.267 |
| 87 | 19 | F | Rt | Nonlesion | Lt and Rt | NA | Lt F | 1 (OXC) | 1.919 |
| 88 | 15 | F | Rt | Nonlesion | Rt | Rt | Rt P | 2 (ZNS, LTG) | 1.497 |
| 89 | 14 | F | Rt | Tumor | Rt | Rt | Rt I | 2 (LCM, LTG) | 1.729 |
| 90 | 5 | F | Lt | Dysplasia | Lt and Rt | Rt | Rt F | 1 (LEV) | 2.338 |
| 91 | 16 | F | Rt | Nonlesion | Rt | NA | NA | 2 (TPM, CLB) | 1.363 |
| 92 | 13 | M | Rt | Dysplasia | Lt | Lt | Lt O | 3 (LEV, LCM, OXC) | 1.563 |
| 93 | 16 | F | Rt | Nonlesion | Lt and Rt | Rt | Rt F | 2 (OXC, LTG) | 0.915 |
| 94 | 9 | M | Rt | Tumor | Lt | Lt | Lt T | 1 (OXC) | 2.304 |
| 95 | 13 | M | Rt | Nonlesion | Lt and Rt | Rt | Rt F | 2 (OXC, LEV) | 1.078 |
| 96 | 11 | M | Rt | Dysplasia | Lt | Lt | Lt TO | 5 (OXC, CLB, LCM, VPA, PER) | 2.339 |
| 97 | 13 | F | Rt | Nonlesion | Rt | Rt | Rt TO | 3 (CBZ, OXC, ESL) | 2.279 |
| 98 | 20 | M | Rt | Nonlesion | Rt | Rt | Rt F | 2 (LCM, ESL) | 1.332 |
| 99 | 15 | F | Rt | Nonlesion | Lt and Rt | Rt | Rt TPO; Lt TO | 2 (OXC, TPM) | 1.577 |
| 100 | 8 | F | Rt | Dysplasia | Lt and Rt | Rt | Rt P | 3 (OXC, CLB, LCM) | 2.196 |
| 101 | 6 | M | Rt | Others | Lt and Rt | Rt | Rt TP | 3 (CLB, OXC, LCM) | 0.865 |
| 102 | 16 | M | Rt | Tumor | Lt | Lt | NA | 1 (LEV) | 2.867 |
| 103 | 17 | M | Lt | Tumor | Lt | Lt | NA | 3 (LEV, OXC, CLB) | 1.384 |
| 104 | 13 | M | Rt | Nonlesion | Lt and Rt | Rt | Rt F | 3 (DZP, VPA, OXC) | 1.994 |
| 105 | 17 | M | Rt | Nonlesion | Lt | Lt** | NA | 2 (LEV, OXC) | 1.162 |
| 106 | 15 | F | Rt | Others | Lt and Rt | Rt | Rt TPO | 2 (LEV, CLB) | 1.767 |

**Supplementary Table 1 Patient profiles.** Forty-nine females (F), and 57 males (M) were included in the study. Lt: left. Rt: right. SOZ: seizure onset zone. MRI lesions other than dysplasia and tumor included focal cortical atrophy (patient 33, 101, 106), hippocampal sclerosis (patients 36, 48, 51, 52, 78), arachnoid cyst (patient 41), and arteriovenous malformations (patient 45). *: multiple subpial transections on the right Rolandic areas. **: responsive neurostimulation employed to the left temporal region. F: Frontal. T: Temporal. O: Occipital. P: Parietal. I: insula. CBZ: Carbamazepine. CLB: Clobazam. CZP: Clonazepam. DZP: Diazepam. ESL: Eslicarbazepine. LCM: Lacosamide. LEV: Levetiracetam. LTG: Lamotrigine. OXC: Oxcarbazepine. PER: Perampanel. PHT: Phenytoin. TPM: Topiramate. VPA: Valproic acid. ZNS: Zonisamide.

| **Regions of interest** | **Left** | **Right** |
| --- | --- | --- |
| SFG: Superior frontal gyrus | 122 (26) | 202 (43) |
| RMF: Rostral middle frontal gyrus | 182 (35) | 330 (49) |
| CMF: Caudal middle frontal gyrus | 181 (40) | 235 (51) |
| MOF: Medial orbitofrontal gyrus | 24 (18) | 34 (22) |
| LOF: Lateral orbitofrontal gyrus | 98 (33) | 100 (33) |
| POP: Pars opercularis | 126 (37) | 97 (42) |
| POR: Pars orbitalis | 72 (27) | 77 (26) |
| PTR: Pars triangularis | 89 (28) | 167 (40) |
| CAC: Caudal anterior cingulate gyrus | 5 ( 5) | 12 ( 7) |
| STG: Superior temporal gyrus | 308 (44) | 332 (51) |
| MTG: Middle temporal gyrus | 244 (42) | 194 (43) |
| ITG: Inferior temporal gyrus | 159 (42) | 189 (43) |
| FG: Fusiform gyrus | 197 (41) | 211 (49) |
| EG: Entorhinal gyrus | 31 (15) | 70 (28) |
| PHG: Parahippocampal gyrus | 23 (13) | 23 (16) |
| PreCG: Precentral gyrus | 383 (46) | 522 (56) |
| PoCG: Postcentral gyrus | 360 (49) | 403 (55) |
| PCL: Paracentral lobule | 24 (11) | 54 (22) |
| SPG: Superior parietal gyrus | 57 (15) | 46 (16) |
| IPG: Inferior parietal gyrus | 81 (21) | 150 (34) |
| SMG: Supramarginal gyrus | 298 (45) | 317 (49) |
| LOG: Lateral occipital gyrus | 195 (34) | 210 (40) |
| Pcun: Precuneus gyrus | 41 (19) | 61 (24) |
| CG: Cuneus gyrus | 18 (12) | 33 (19) |
| LG: Lingual gyrus | 111 (30) | 166 (44) |
| ICG: Isthmus cingulate gyrus | 16 ( 9) | 37 (20) |
| PCG: Posterior cingulate gyrus | 19 ( 9) | 41 (20) |
| RAC: Rostral anterior cingulate gyrus | 3 ( 1) | 4 ( 2) |
| IG: Insular gyrus | 2 ( 1) | 0 ( 0) |
| TTG: Transverse temporal gyrus | 0 ( 0) | 0 ( 0) |
| Per: Pericalcarine gyrus | 5 ( 3) | 1 ( 1) |
| Total | 3474 (63) | 4318 (67) |

**Supplementary Table 2 Number of artifact-free, nonepileptic electrode sites in regions of interest (ROIs).** Numbers in parentheses indicate the number of contributing patients. The ROI-based analysis was performed using those with at least five cortical electrode sites from at least three patients in each of the left and right hemispheres. Consequently, the ROI-based analysis was not performed for RAC, IG, TTG, or Per.

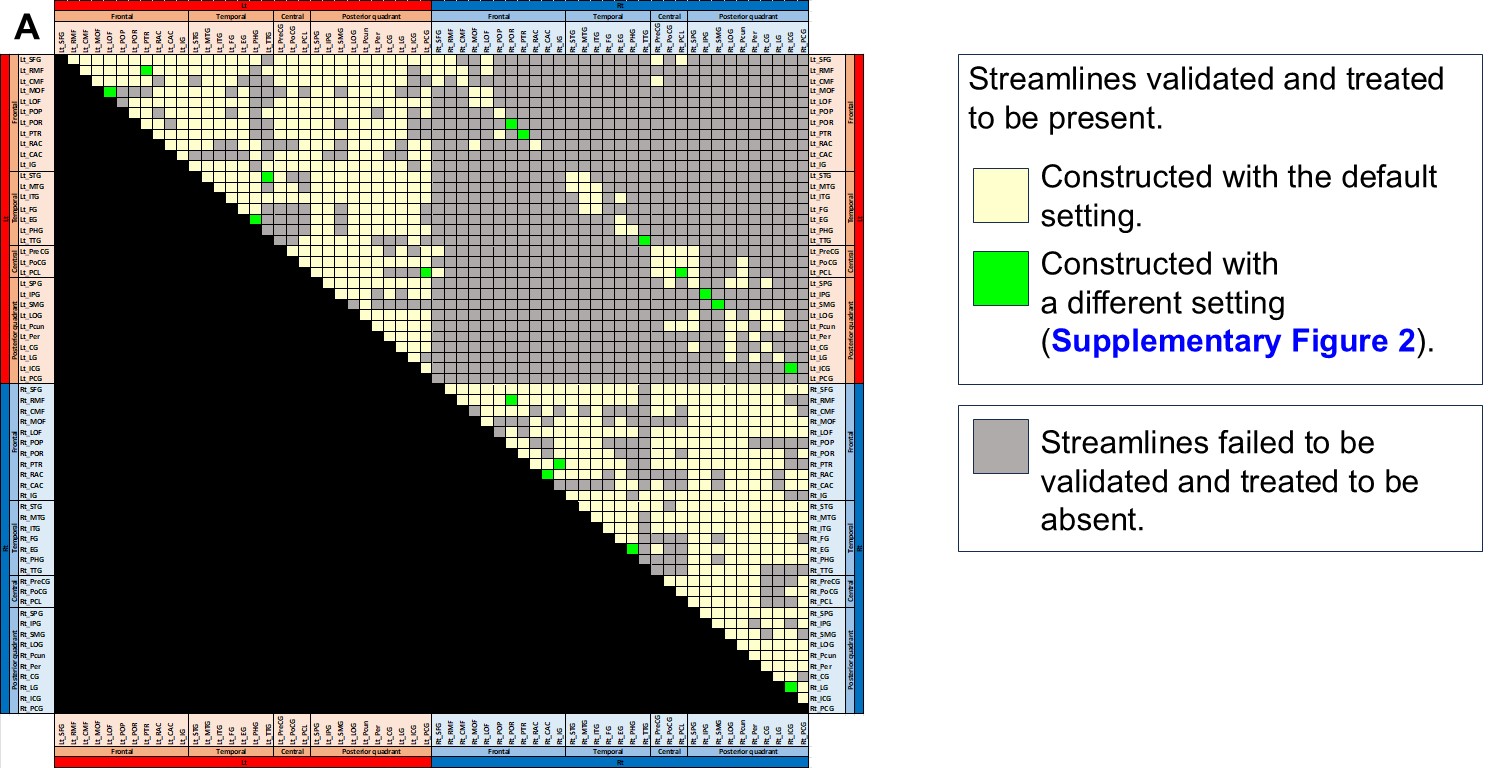

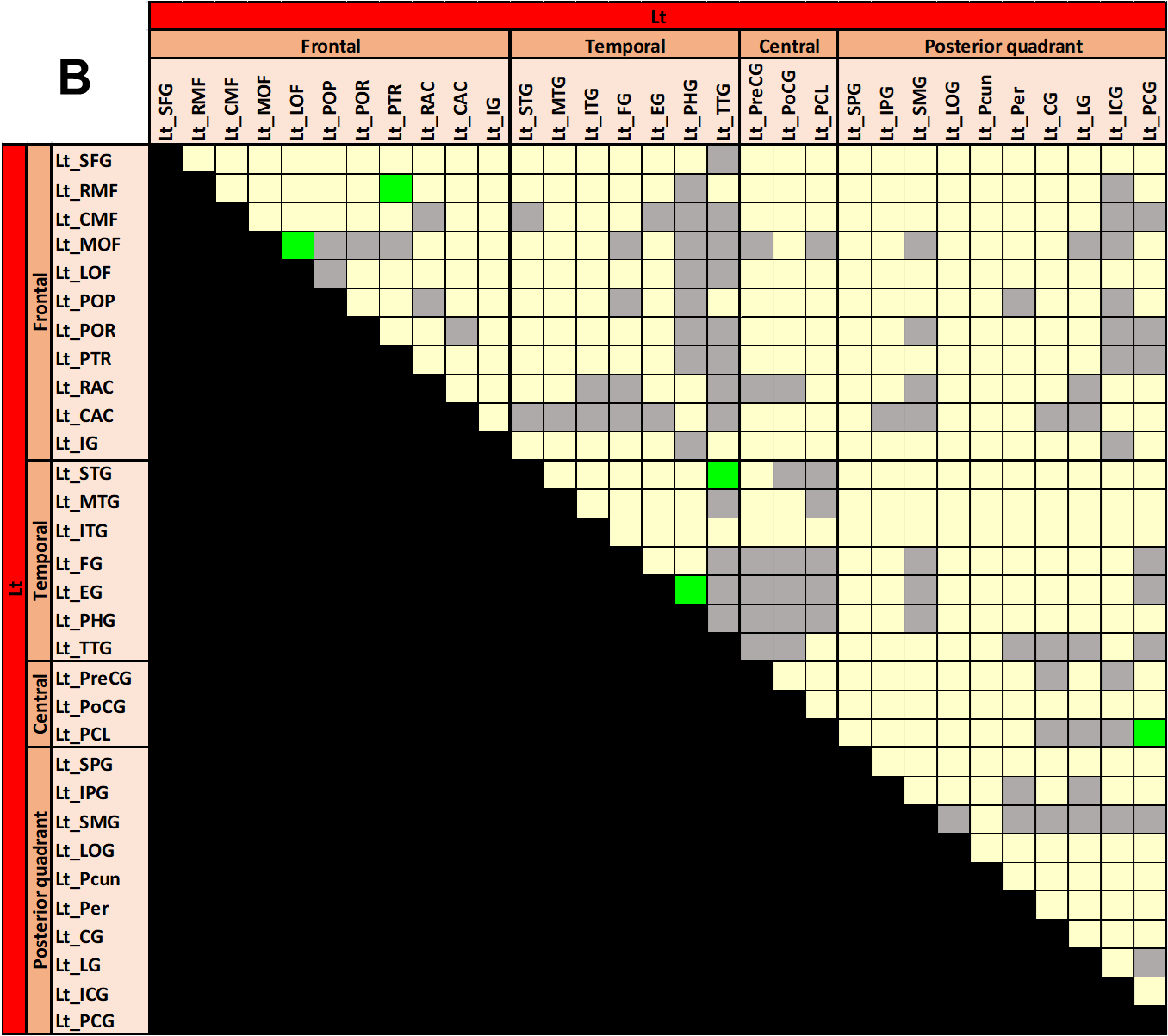

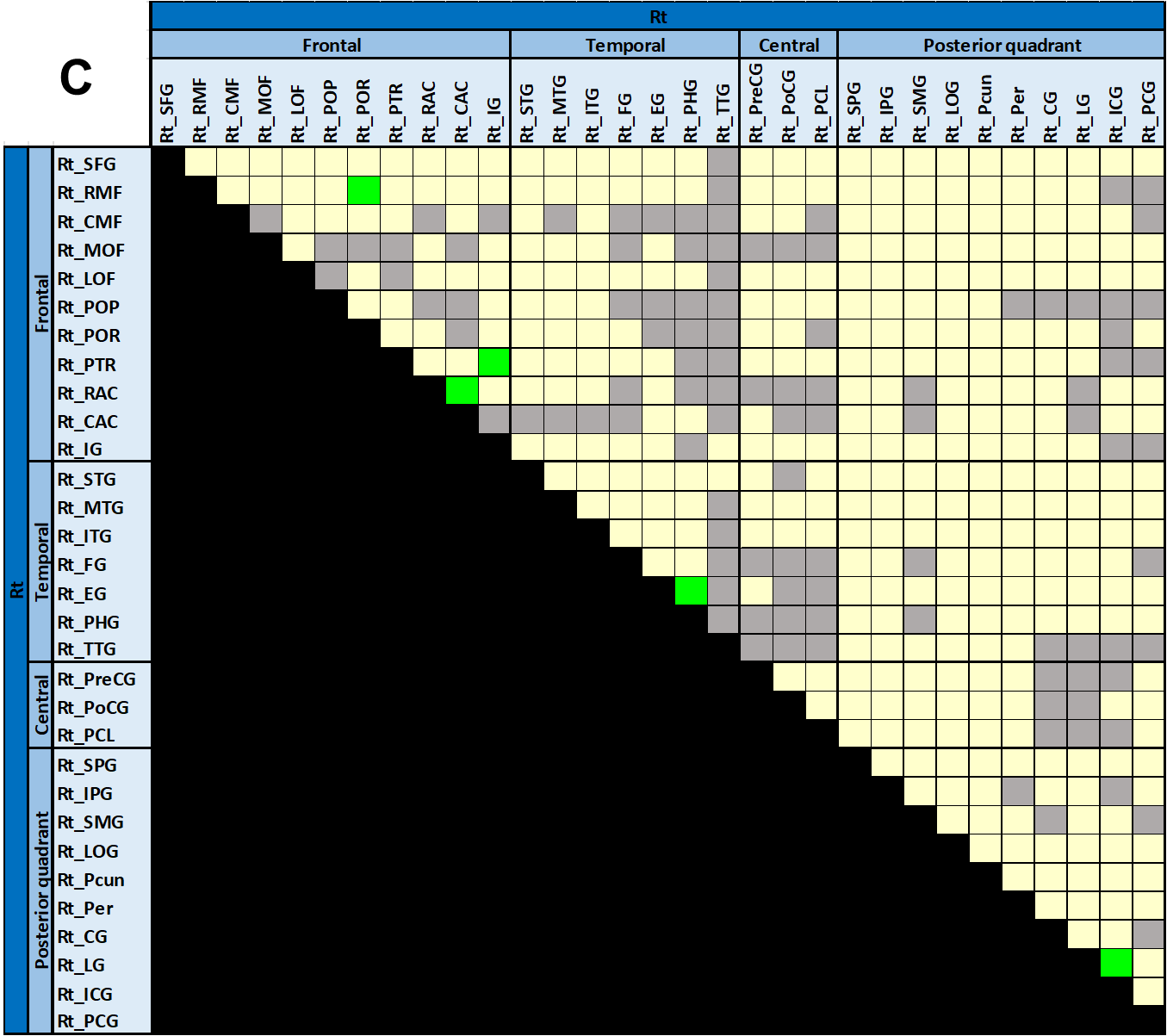

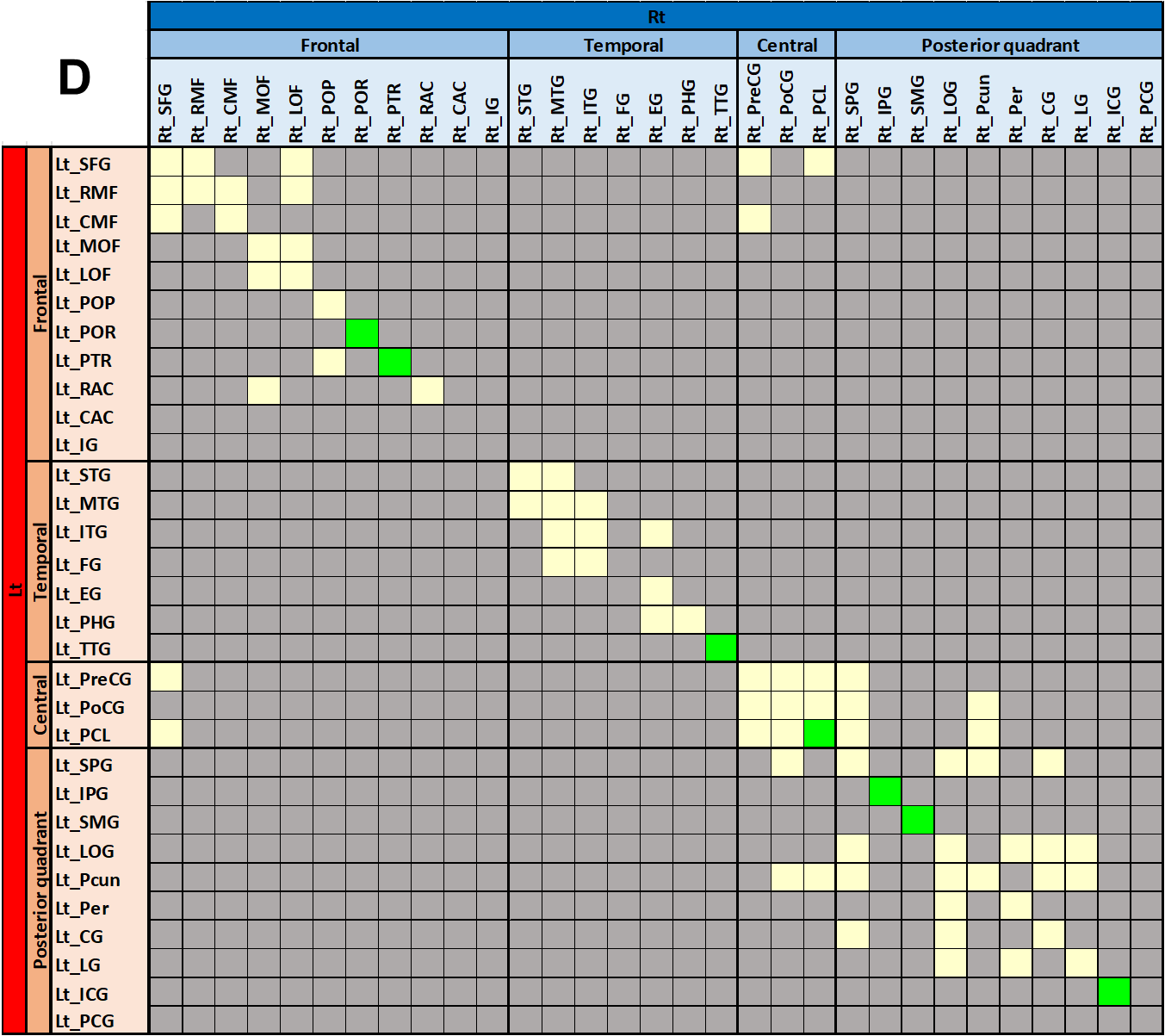

**Supplementary Figure 1 Construction and visualization of anatomical white matter streamlines on tractography.** (A) Here, we present the anatomical white matter streamline connectivity matrix used in the analysis of functional connectivity modulations. (B) Zoomed image focusing on the intra-hemispheric white matter streamlines within the left hemisphere. (C) Zoomed image focusing on the intra-hemispheric white matter streamlines within the right hemisphere. (D) Zoomed image focusing on the inter-hemispheric white matter streamlines. Pairs of regions of interest (ROIs) indicated by cream and light green squares reflect the presence of white matter streamlines validated and visualized in the functional connectivity atlas in the present study (cream: constructed with the default parameter; light green: constructed with a different parameter as detailed in **Supplementary Figure 2**). Pairs of ROIs indicated by gray squares reflect the failure to construct or validate white matter streamlines; thus, anatomical connectivity between these ROIs was not visualized in the functional connectivity atlas. Abbreviated anatomical cortical ROIs: Refer to the list in **Supplementary Table 2**. The anatomical white matter streamline template is available as open-source material to allow investigators to replicate our tractography analysis (<https://github.com/a8k8nn0/TractographyAtlas>).

Using the DSI Studio script (<http://dsi-studio.labsolver.org/>) within the MNI standard space, we constructed and visualized the streamline connecting each pair of ROIs with the shortest length among those detected using the specified parameters as shown below. The default fiber tracking parameters were a quantitative anisotropy threshold of 0.05, a maximum turning angle of 70°, and a streamline length of 20 to 250 mm. Board-certified neurosurgeons (A.K. and R.K.) employed a visual validation procedure, excluding artifactual streamlines and those involving the brainstem, basal ganglia, or thalamus from the tractography analysis. Artifactual streamlines included those with highly irregular paths, looping back on themselves, or terminating in unlikely locations such as the cerebrospinal fluid space (Melhem et al., 2002; Mukherjee et al., 2008; Tournier et al., 2011; Guevara et al., 2012; Donahue et al., 2016). We then visualized a validated streamline connecting each ROI pair with the shortest distance. Only if local u-fibers between immediately adjacent ROIs or callosal fibers between precisely homotopic ROIs failed to be constructed by the aforementioned procedure, we employed different tracking parameters for these pairs. To maximize the transparency of our methodological approach, in **Supplementary Figure 2**, we annotate the fiber tracking parameters used for each streamline visualization. In this study, we aimed to inclusively visualize biologically plausible white matter pathways. This decision was partly based on our previous study (Kitazawa et al., 2023), which found that the default fiber tracking parameters failed to visualize some local u-fibers between immediately adjacent ROIs and inter-hemispheric fibers between precisely homotopic cortical ROIs. Collective evidence suggests that immediately neighboring ROIs (Schmahmann and Pandya, 2008; Schilling et al., 2018). and precisely homotopic cortical ROIs (Boussaoud et al., 2005; Huang et al., 2005; Hofer et al., 2008; Umeoka et al., 2009; Terada et al., 2012; Caminiti et al., 2013; O'Reilly et al., 2013; Fenlon and Richards, 2015; Horowitz et al., 2015; Lacuey et al., 2015; Lacuey et al., 2016; Ruddy et al., 2017; Lehner et al., 2018; Mancuso et al., 2019; Mitsuhashi et al., 2021; Innocenti et al., 2022) are directly connected by white matter fibers and capable of bidirectional communication.

**Supplementary reference**

Boussaoud, D., Tanné-Gariépy, J., Wannier, T., Rouiller, E.M., 2005. Callosal connections of dorsal versus ventral premotor areas in the macaque monkey: A multiple retrograde tracing study. BMC Neurosci. 6, 67. https://doi.org/10.1186/1471-2202-6-67.

Caminiti, R., Carducci, F., Piervincenzi, C., Battaglia-Mayer, A., Confalone, G., Visco-Comandini, F., Pantano, P., Innocenti, G.M., 2013. Diameter, length, speed, and conduction delay of callosal axons in macaque monkeys and humans: Comparing data from histology and magnetic resonance imaging diffusion tractography. J. Neurosci. 33, 14501–14511. https://doi.org/10.1523/JNEUROSCI.0761-13.2013.

Donahue, C.J., Sotiropoulos, S.N., Jbabdi, S., Hernandez-Fernandez, M., Behrens, T.E., Dyrby, T.B., Coalson, T., Kennedy, H., Knoblauch, K., Van Essen, D.C., Glasser, M.F., 2016. Using diffusion tractography to predict cortical connection strength and distance: A quantitative comparison with tracers in the monkey. J. Neurosci. 36, 6758–6770. https://doi.org/10.1523/JNEUROSCI.0493-16.2016.

Fenlon, L.R., Richards, L.J., 2015. Contralateral targeting of the corpus callosum in normal and pathological brain function. Trends. Neurosci. 38, 264–272. https://doi.org/10.1016/j.tins.2015.02.007.

Guevara, P., Duclap, D., Poupon, C., Marrakchi-Kacem, L., Fillard, P., Le Bihan, D., Leboyer, M., Houenou, J., Mangin, J.F., 2012. Automatic fiber bundle segmentation in massive tractography datasets using a multi-subject bundle atlas. Neuroimage. 61, 1083–1099. https://doi.org/10.1016/j.neuroimage.2012.02.071.

Hofer, S., Merboldt, K.D., Tammer, R., Frahm, J., 2008. Rhesus monkey and human share a similar topography of the corpus callosum as revealed by diffusion tensor MRI in vivo. Cereb. Cortex. 18, 1079–1084. https://doi.org/10.1093/cercor/bhm141.

Horowitz, A., Barazany, D., Tavor, I., Bernstein, M., Yovel, G., Assaf, Y., 2015. In vivo correlation between axon diameter and conduction velocity in the human brain. Brain. Struct. Funct. 220, 1777–1788. https://doi.org/10.1007/s00429-014-0871-0.

Huang, H., Zhang, J., Jiang, H., Wakana, S., Poetscher, L., Miller, M.I., van Zijl, P.C.M., Hillis, A.E., Wytik, R., Mori, S., 2005. DTI tractography based parcellation of white matter: Application to the mid-sagittal morphology of corpus callosum. Neuroimage. 26, 195–205. https://doi.org/10.1016/j.neuroimage.2005.01.019.

Innocenti, G. M., Schmidt, K., Milleret, C., Fabri, M., Knyazeva, M.G., Battaglia-Mayer, A., Aboitiz, F., Ptito, M., Caleo, M., Marzi, C.A., Barakovic, M., Lepore, F., Caminiti, R., 2022. The functional characterization of callosal connections. Prog. Neurobiol. 208, 102186. https://doi.org/10.1016/j.pneurobio.2021.102186.

Kitazawa, Y., Sonoda, M., Sakakura, K., Mitsuhashi, T., Firestone, E., Ueda, R., Kambara, T., Iwaki, H., Luat, A.F., Marupudi, N.I., Sood, S., Asano, E., 2023. Intra- and inter-hemispheric network dynamics supporting object recognition and speech production. Neuroimage. 270, 119954. https://doi.org/10.1016/j.neuroimage.2023.119954.

Lacuey, N., Zonjy, B., Kahriman, E. S., Kaffashi, F., Miller, J., Lüders, H.O., 2015. Functional connectivity between right and left mesial temporal structures. Brain. Struct. Funct. 220, 2617–2623. https://doi.org/10.1007/s00429-014-0810-0.

Lacuey, N., Zonjy, B., Kahriman, E.S., Marashly, A., Miller, J., Lhatoo, S.D., Lüders, H.O., 2016. Homotopic reciprocal functional connectivity between anterior human insulae. Brain. Struct. Funct. 221, 2695–2701. https://doi.org/10.1007/s00429-015-1065-0.

Lehner, K.R., Yeagle, E.M., Argyelan, M., Klimaj, Z., Du, V., Megevand, P., Hwang, S.T., Mehta, A.D., 2018. Validation of corpus callosotomy after laser interstitial thermal therapy: A multimodal approach. J. Neurosurg. 131, 1095–1105. https://doi.org/10.3171/2018.4.JNS172588.

Mancuso, L., Uddin, L.Q., Nani, A., Costa, T., Cauda, F., 2019. Brain functional connectivity in individuals with callosotomy and agenesis of the corpus callosum: A systematic review. Neurosci. Biobehav. Rev. 105, 231–248. https://doi.org/10.1016/j.neubiorev.2019.07.004.

Melhem, E.R., Mori, S., Mukundan, G., Kraut, M.A., Pomper, M.G., van Zijl, P.C.M., 2002. Diffusion tensor MR imaging of the brain and white matter tractography. AJR. Am. J. Roentgenol. 178, 3–16. https://doi.org/10.2214/ajr.178.1.1780003.

Mitsuhashi, T., Sonoda, M., Jeong, J.W., Silverstein, B.H., Iwaki, H., Luat, A.F., Sood, S., Asano, E., 2021. Four-dimensional tractography animates propagations of neural activation via distinct interhemispheric pathways. Clin. Neurophysiol. 132, 520–529. https://doi.org/10.1016/j.clinph.2020.11.030.

Mukherjee, P., Chung, S.W., Berman, J.I., Hess, C.P., Henry, R.G., 2008. Diffusion tensor MR imaging and fiber tractography: Technical considerations. AJNR. Am. J. Neuroradiol. 29, 843–852. https://doi.org/10.3174/ajnr.A1052.

O'Reilly, J.X., Croxson, P.L., Jbabdi, S., Sallet, J., Noonan, M.P., Mars, R.B., Browning, P.G.F., Wilson, C.R.E., Mitchell, A.S., Miller, K.L., Rushworth, M.F.S., Baxter, M.G., 2013. Causal effect of disconnection lesions on interhemispheric functional connectivity in rhesus monkeys. Proc. Natl. Acad. Sci. U. S. A. 110, 13982–13987. https://doi.org/10.1073/pnas.1305062110.

Ruddy, K.L., Leemans, A., Carson, R.G., 2017. Transcallosal connectivity of the human cortical motor network. Brain. Struct. Funct. 222, 1243–1252. https://doi.org/10.1007/s00429-016-1274-1.

Schilling, K., Gao, Y., Janve, V., Stepniewska, I., Landman, B.A., Anderson, A.W., 2018. Confirmation of a gyral bias in diffusion MRI fiber tractography. Hum. Brain. Mapp. 39, 1449–1466. https://doi.org/10.1002/hbm.23936.

Schmahmann, J.D., Pandya, D.N., 2008. Disconnection syndromes of basal ganglia, thalamus, and cerebrocerebellar systems. Cortex. 44, 1037–1066. https://doi.org/10.1016/j.cortex.2008.04.004.

Terada, K., Umeoka, S., Usui, N., Baba, K., Usui, K., Fujitani, S., Matsuda, K., Tottori, T., Nakamura, F., Inoue, Y., 2012. Uneven interhemispheric connections between left and right primary sensori-motor areas. Hum. Brain. Mapp. 33, 14–26. https://doi.org/10.1002/hbm.21189.

Tournier, J.D., Mori, S., Leemans, A., 2011. Diffusion tensor imaging and beyond. Magn. Reason. Med. 65, 1532–1556. https://doi.org/10.1002/mrm.22924.

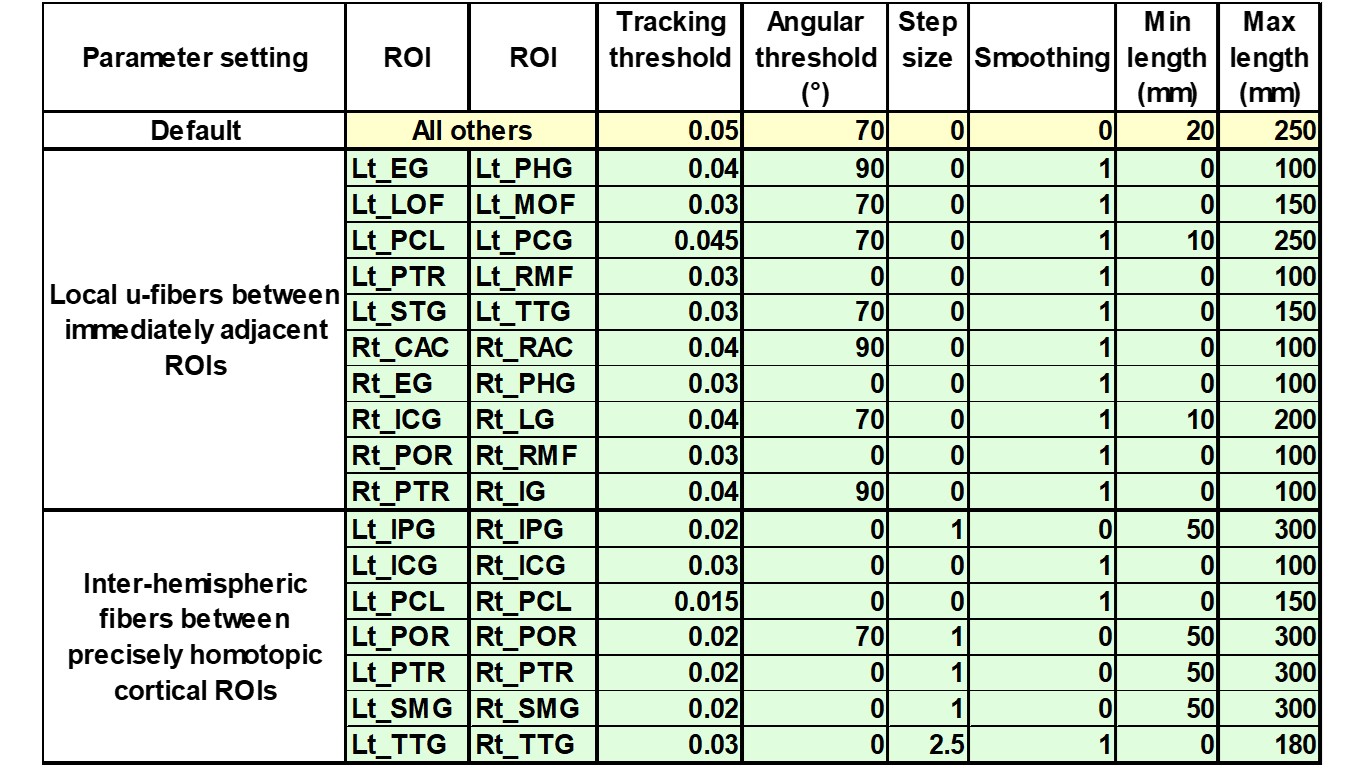
Umeoka, S., Terada, K., Baba, K., Usui, K., Matsuda, K., Tottori, T., Usui, N., Nakamura, F., Inoue, Y., Fujiwara, T., Mihara, T., 2009. Neural connection between bilateral basal temporal regions: Cortico-cortical evoked potential analysis in patients with temporal lobe epilepsy. Neurosurgery. 64, 847–855. https://doi.org/10.1227/01.NEU.0000344001.26669.92.

**Supplementary Figure 2 Fiber tracking parameters used to construct anatomical white matter streamlines on tractography.** A total of 788 white matter streamlines were constructed with the default parameter setting and visually validated. Ten streamlines reflecting local u-fibers connecting immediately neighboring cortical regions of interest (ROIs) and seven streamlines reflecting inter-hemispheric fibers connecting precisely homotopic ROIs were constructed using a different parameter setting. These streamlines were likewise visually validated.

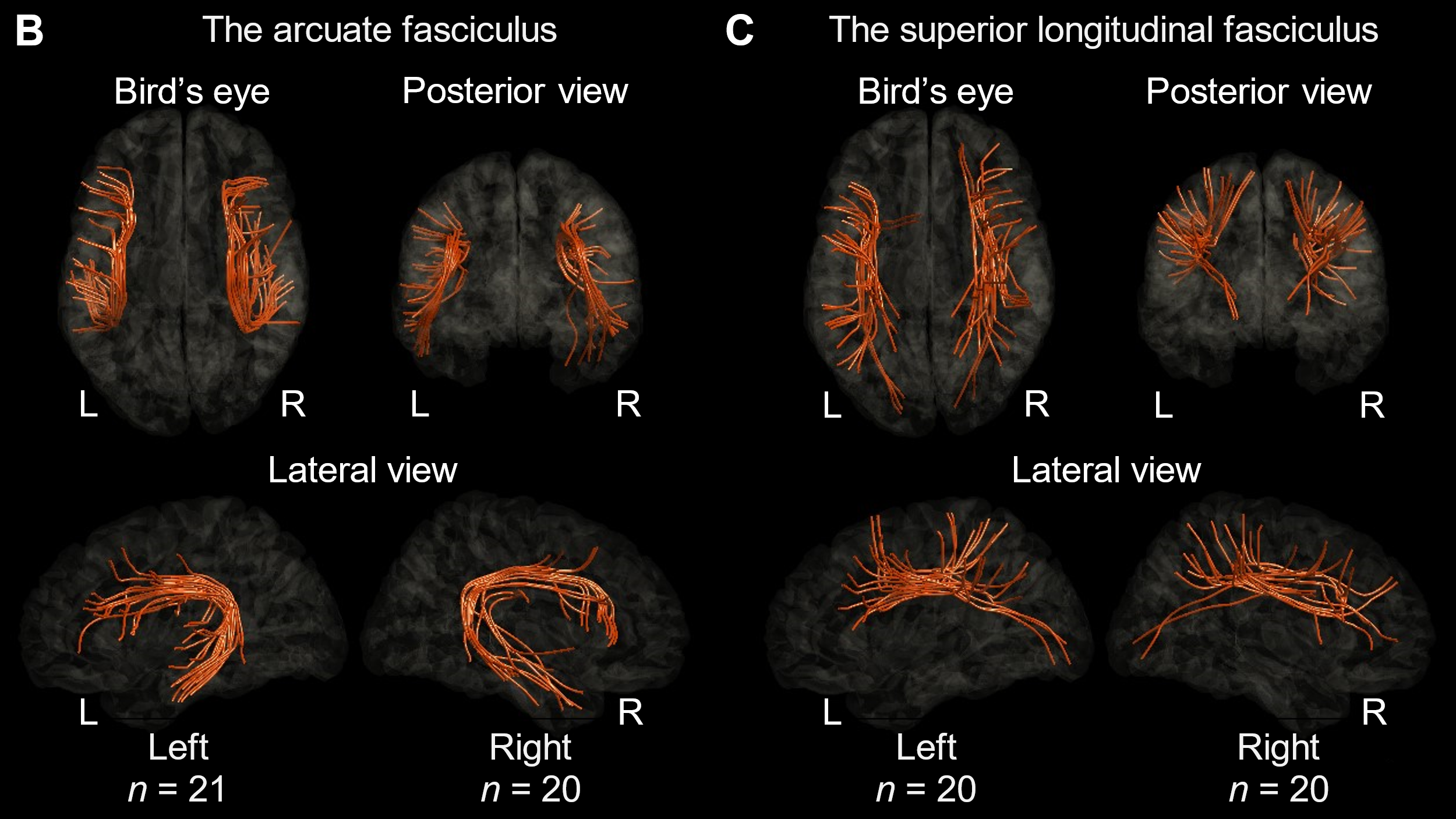

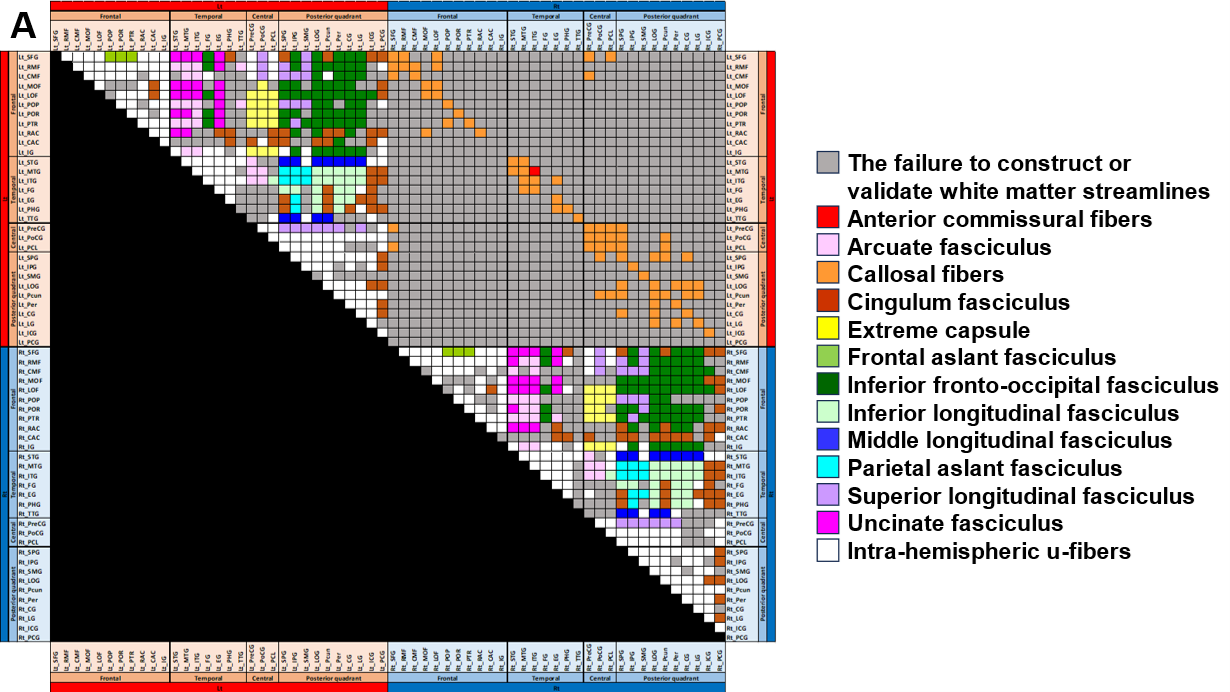

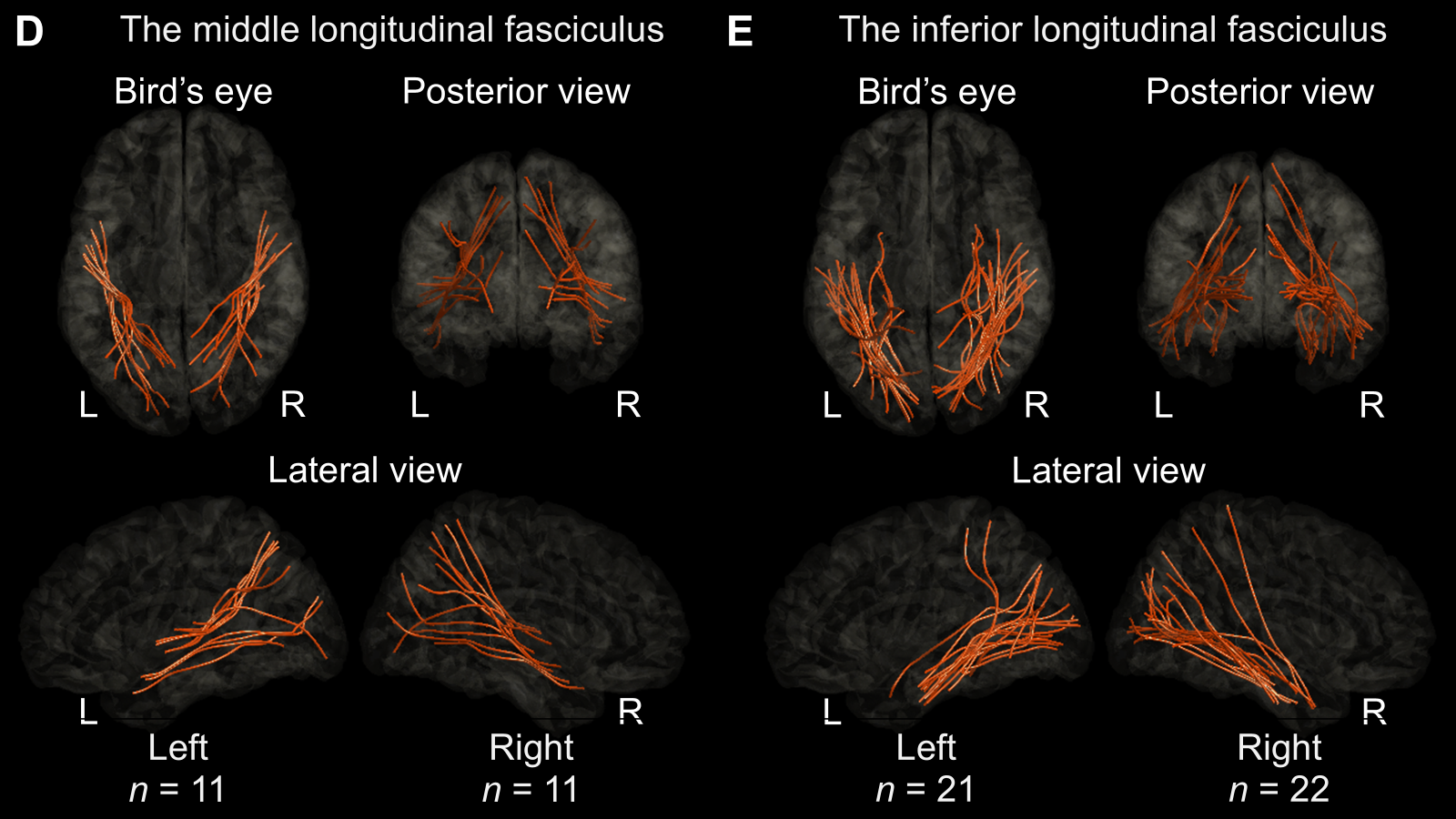

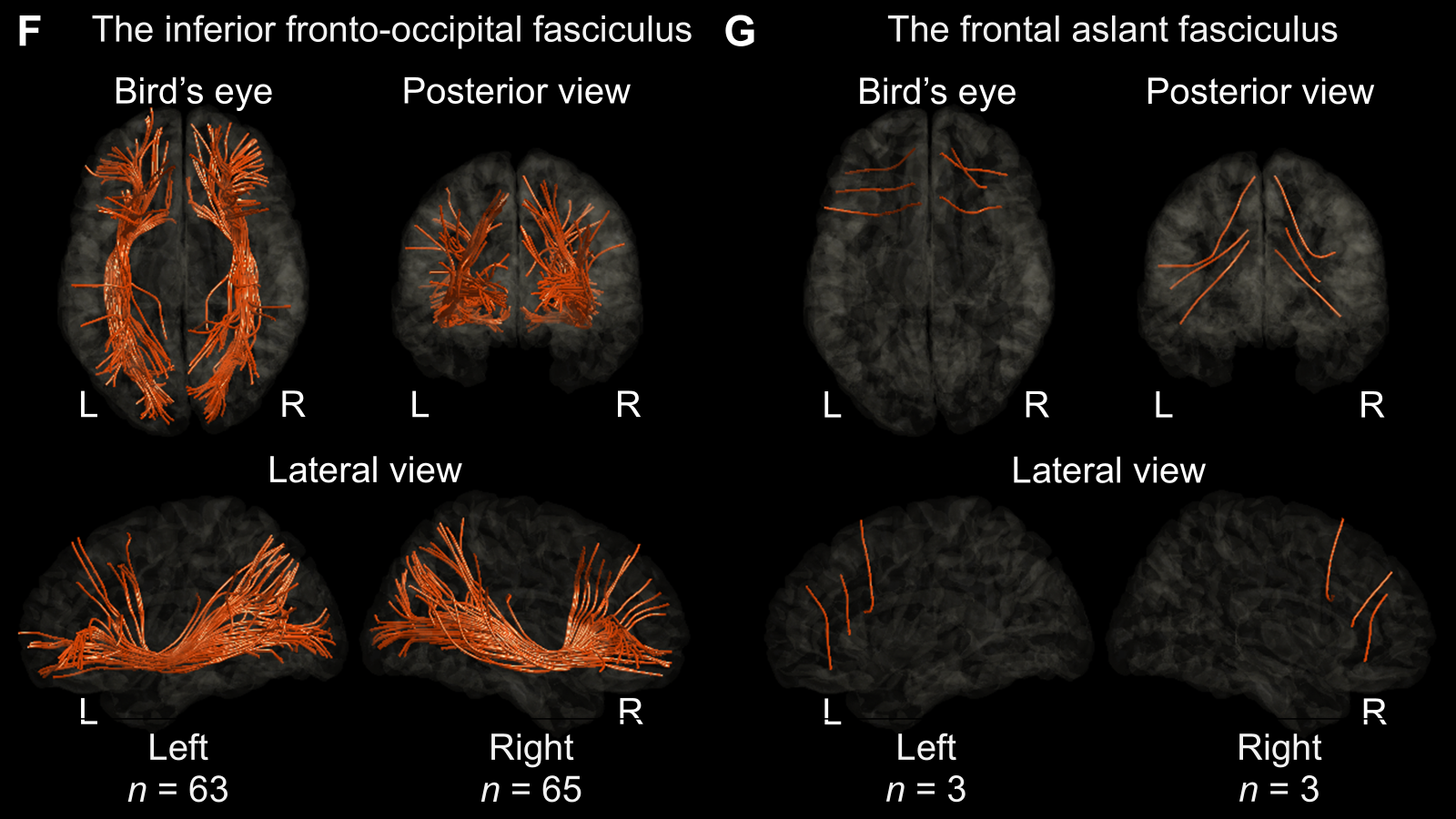

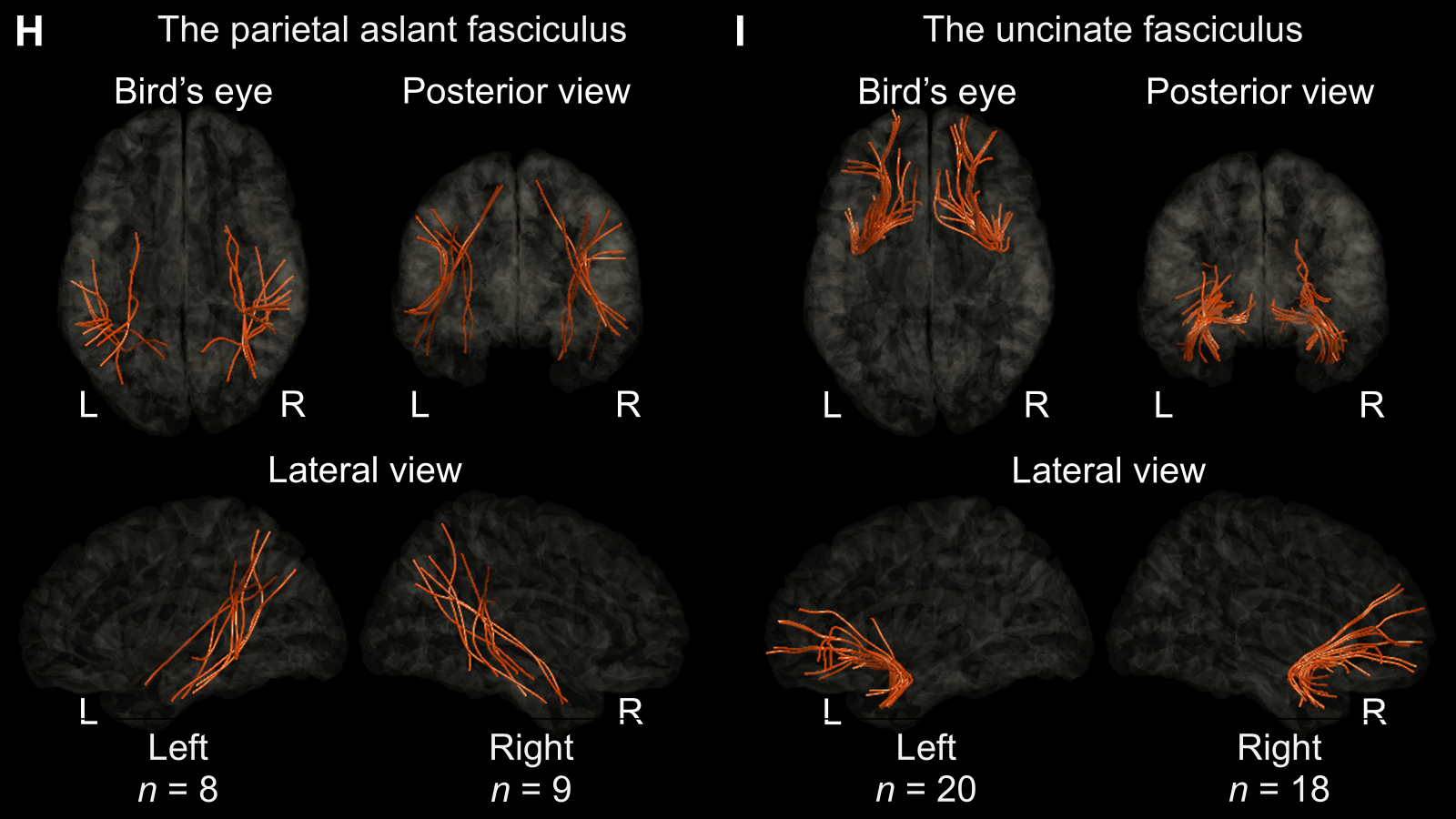

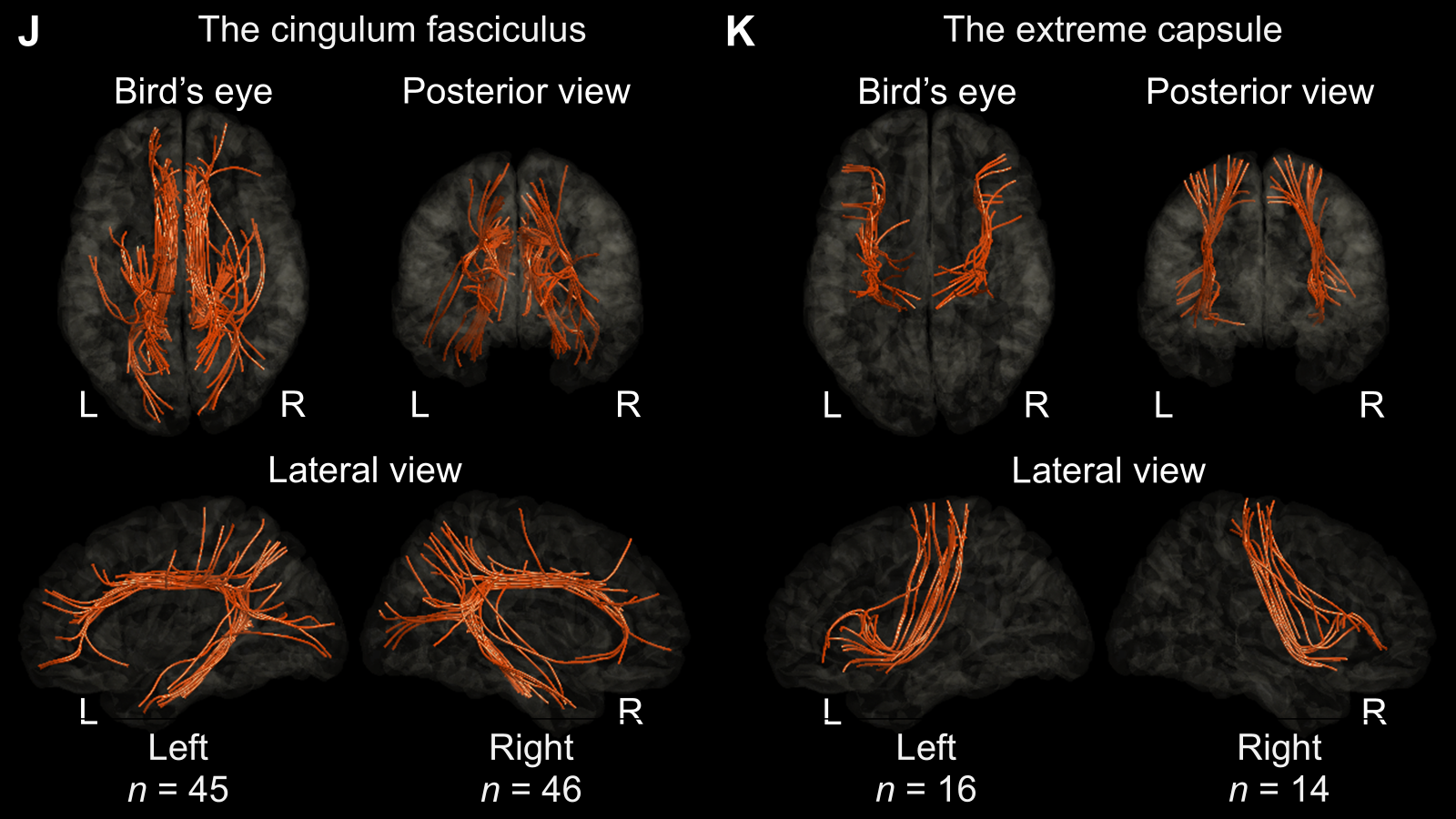

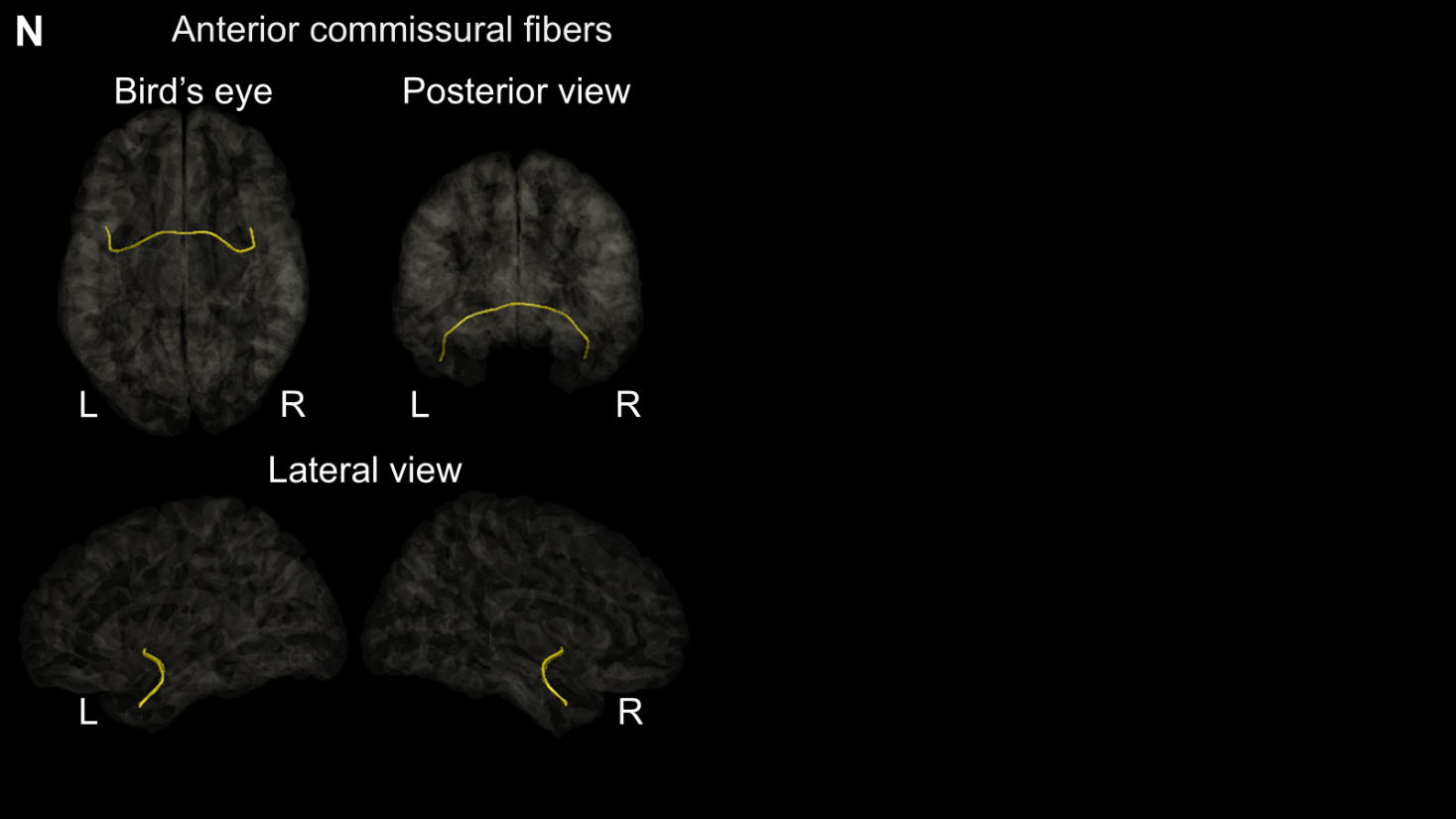

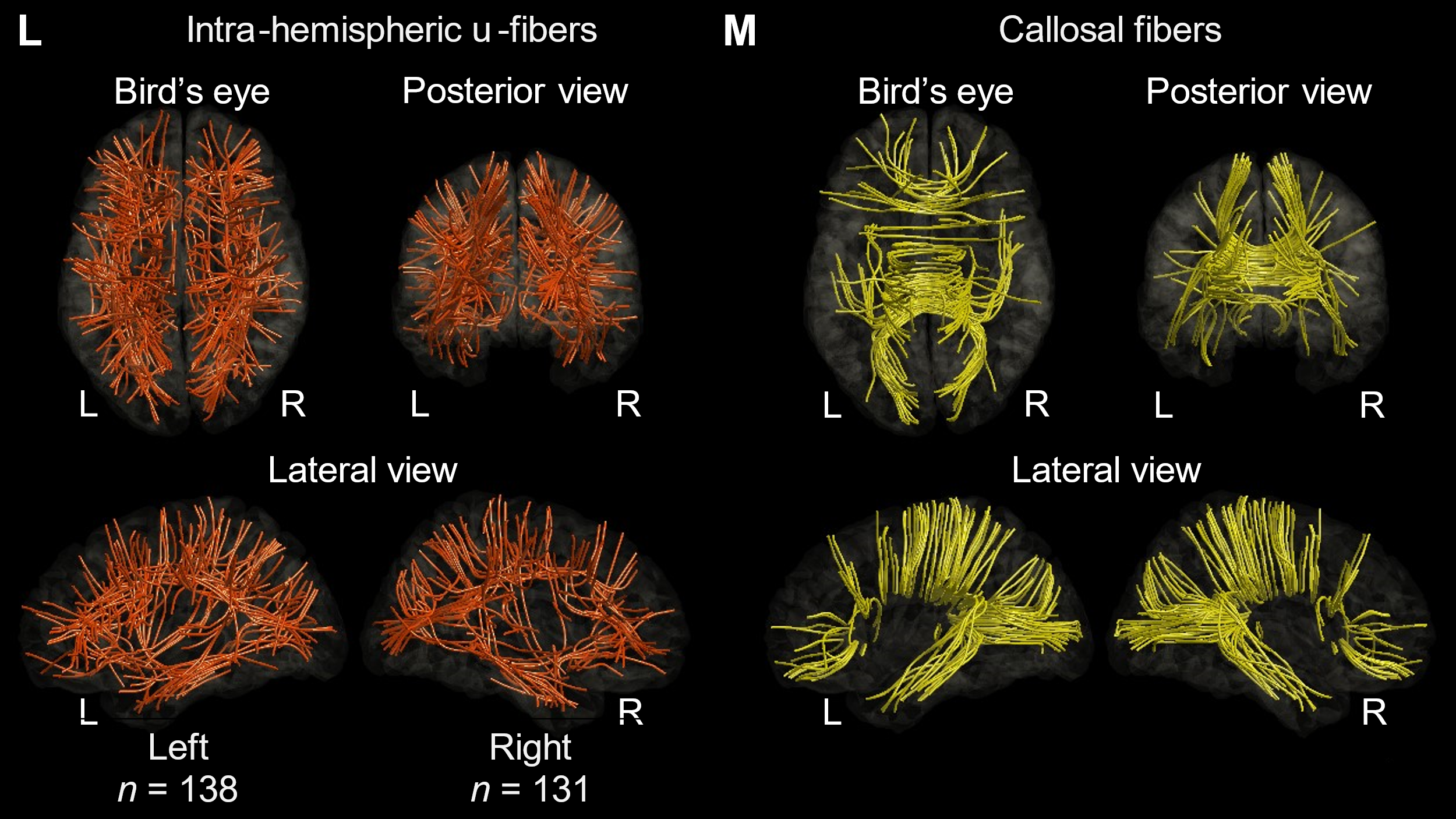

**Supplementary Figure 3 Anatomical labels of white matter streamlines**. (A) In this white matter streamline connectivity matrix, each pair of cortical regions of interest (ROIs) is assigned a corresponding fasciculus label. Board-certified neurosurgeons (A.K. and R.K.) classified each white matter streamline based on anatomical connectivity atlases previously reported (Thiebaut de Schotten et al., 2011; Dick and Tremblay, 2012; Yeh et al., 2018; Hansen et al., 2021; Yeh, 2022). (B) The arcuate fasciculus was defined as a dorsal fiber bundle connecting the temporal lobe to the frontal or parietal lobe. The anatomical courses of white matter streamlines classified as the arcuate fasciculus are visualized. (C) The superior longitudinal fasciculus (SLF) was defined as a dorsal fiber bundle coursing laterally to the lateral ventricle, connecting the occipital or parietal lobe to the frontal lobe. (D) The middle longitudinal fasciculus (MLF) was defined as a ventral fiber bundle connecting the parietal or occipital lobe to the superior temporal gyrus. (E) The inferior longitudinal fasciculus (ILF) was defined as a ventral fiber bundle connecting the occipital or parietal lobe to the temporal lobe. (F) The inferior fronto-occipital fasciculus (IFOF) was defined as a ventral fiber bundle coursing laterally to the lateral ventricle and medially to the insula, connecting the occipital or parietal lobe to the frontal lobe. (G) The frontal aslant fasciculus was defined as a fiber bundle connecting the inferior frontal gyrus to the superior frontal gyrus. (H) The parietal aslant fasciculus was defined as a fiber bundle coursing laterally to the lateral ventricle, connecting the parietal lobe to the temporal lobe. (I) The uncinate fasciculus was defined as a fiber bundle connecting the anterior portion of the temporal lobe to the ventral portion of the frontal lobe through the temporal stem. (J) The cingulum fasciculus was defined as a fiber bundle coursing medially to the lateral ventricle, connecting the occipital, parietal, or temporal lobe to the frontal lobe. (K) The extreme capsule was defined as a fiber bundle connecting the precentral, postcentral, or paracentral gyrus to the inferior frontal or insular gyrus through the external capsule. (L) Intra-hemispheric u-fibers were defined as those not classified as the major fasciculi mentioned above. (M) Callosal fibers were defined as those connecting the left and right hemispheres through the corpus callosum. (N) Anterior commissural fibers were defined as those connecting the left and right temporal lobes through the anterior commissure.

**Supplementary reference**

Dick, A.S., Tremblay, P., 2012. Beyond the arcuate fasciculus: Consensus and controversy in the connectional anatomy of language. Brain. 135, 3529–3550. https://doi.org/10.1093/brain/aws222.

Hansen, C.B., Yang, Q., Lyu, I., Rheault, F., Kerley, C., Chandio, B.Q., Fadnavis, S., Williams, O., Shafer, A.T., Resnick, S. M., Zald, D.H., Cutting, L.E., Taylor, W.D., Boyd, B., Garyfallidis, E., Anderson, A.W., Descoteaux, M., Landman, B.A., Schilling, K.G., 2021. Pandora: 4-D white matter bundle population-based atlases derived from diffusion MRI fiber tractography. Neuroinformatics. 19, 447–460. https://doi.org/10.1007/s12021-020-09497-1.

Thiebaut de Schotten, M., Ffytche, D.H., Bizzi, A., Dell'Acqua, F., Allin, M., Walshe, M., Murray, R., Williams, S.C., Murphy, D.G.M., Catani, M., 2011. Atlasing location, asymmetry and inter-subject variability of white matter tracts in the human brain with MR diffusion tractography. Neuroimage, 54, 49–59. https://doi.org/10.1016/j.neuroimage.2010.07.055.

Yeh, F.C., Panesar, S., Fernandes, D., Meola, A., Yoshino, M., Fernandez-Miranda, J.C., Vettel, J.M., Verstynen, T., 2018. Population-averaged atlas of the macroscale human structural connectome and its network topology. Neuroimage. 178, 57–68. https://doi.org/10.1016/j.neuroimage.2018.05.027.

Yeh, F.C., 2022. Population-based tract-to-region connectome of the human brain and its hierarchical topology. Nat. Commun. 13(1), 4933. https://doi.org/10.1038/s41467-022-32595-4.

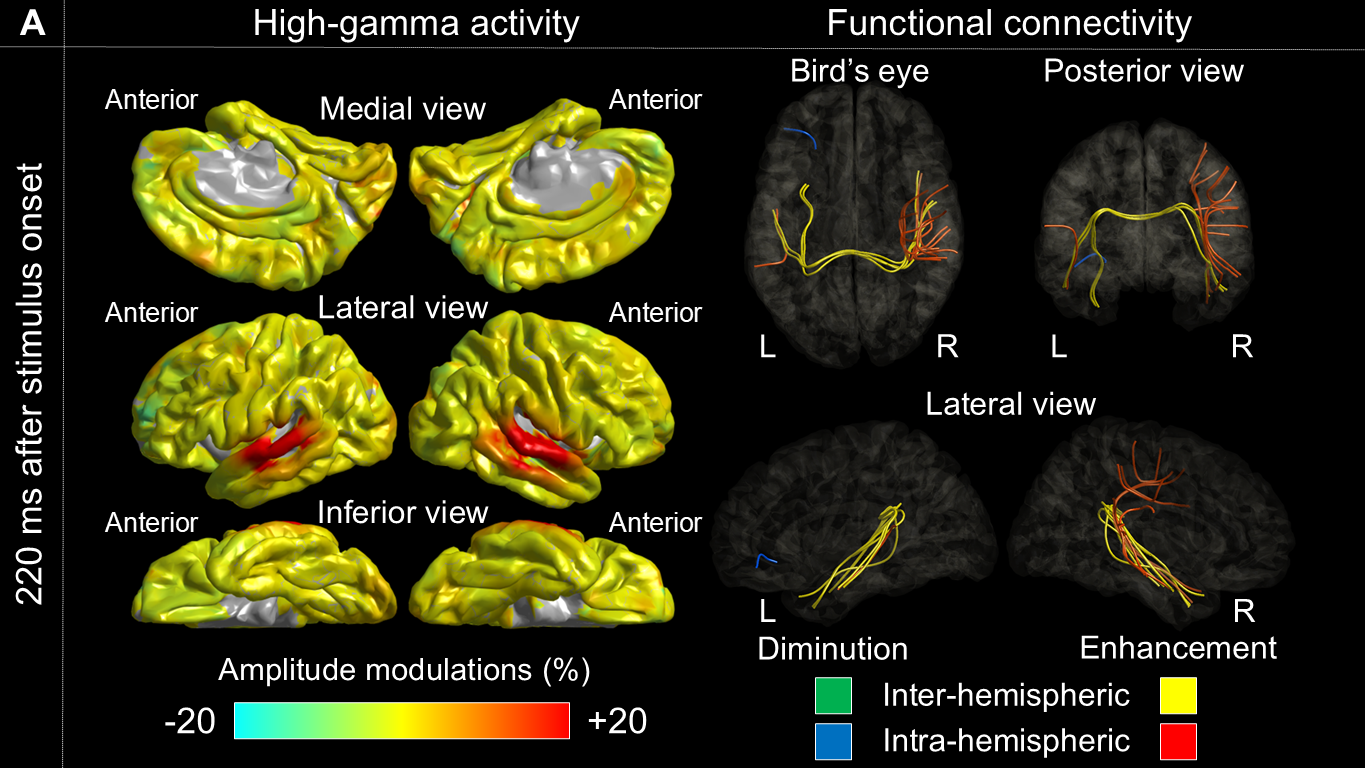

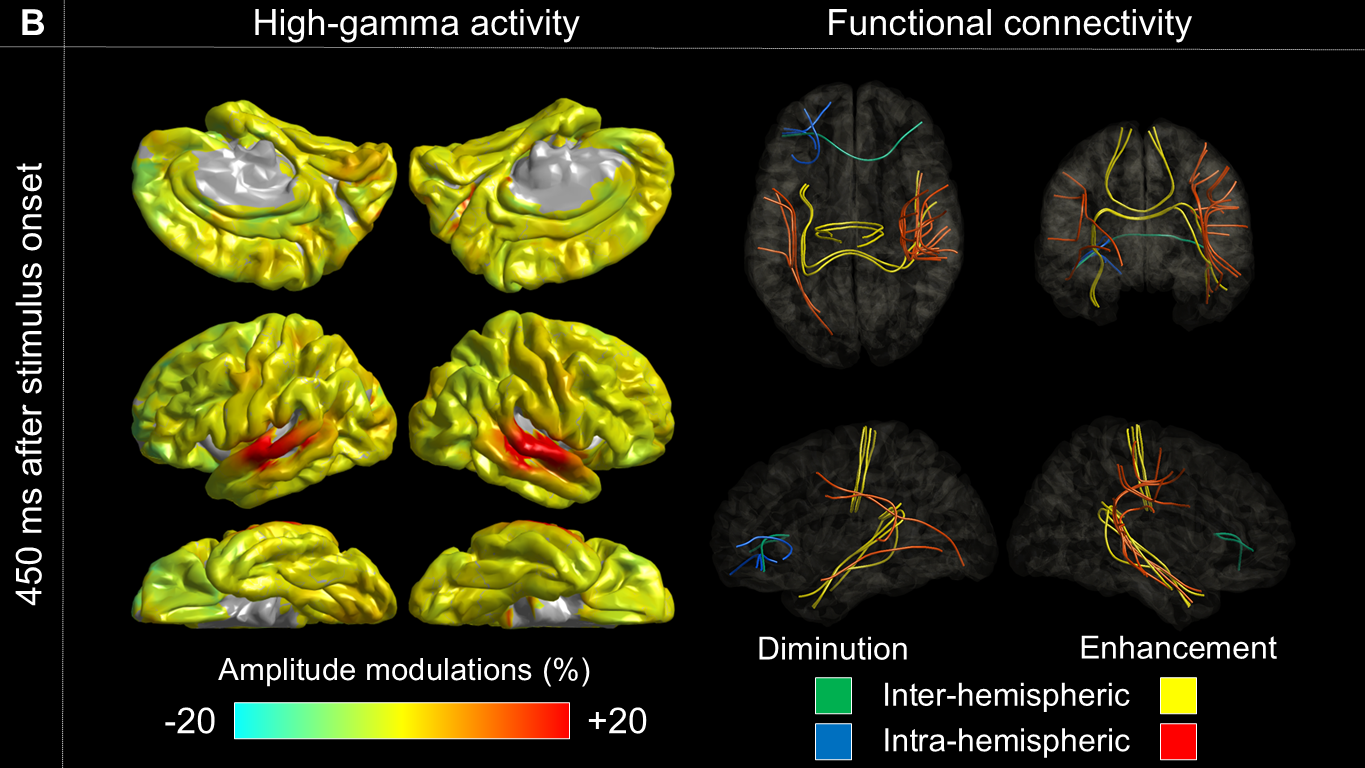

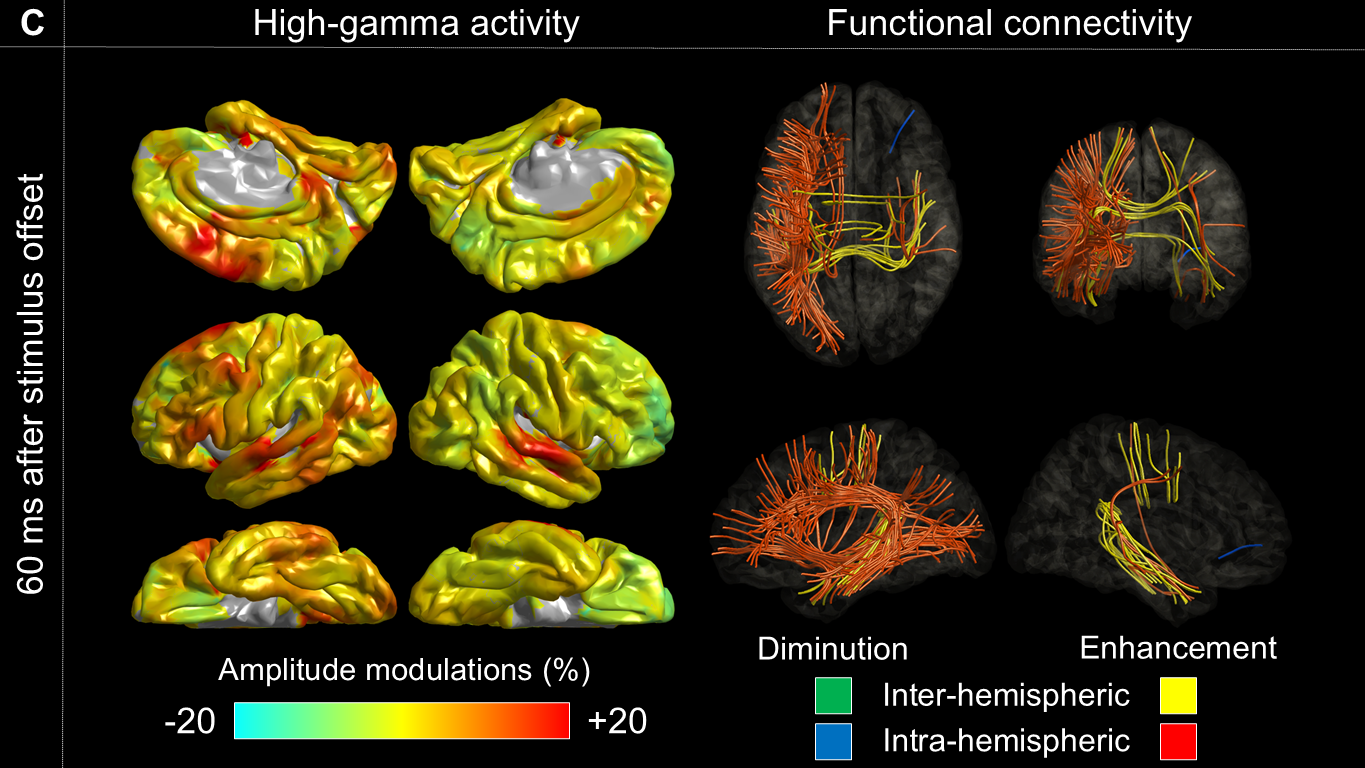

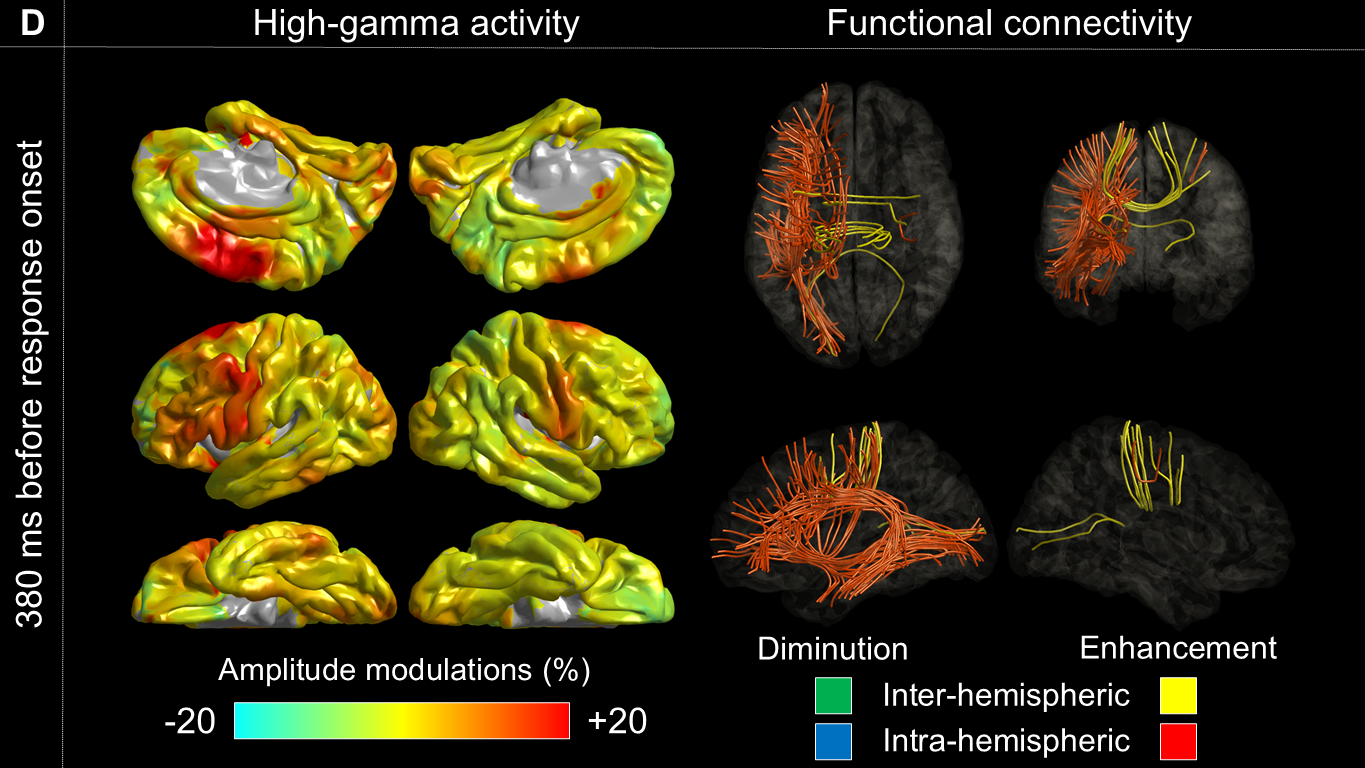

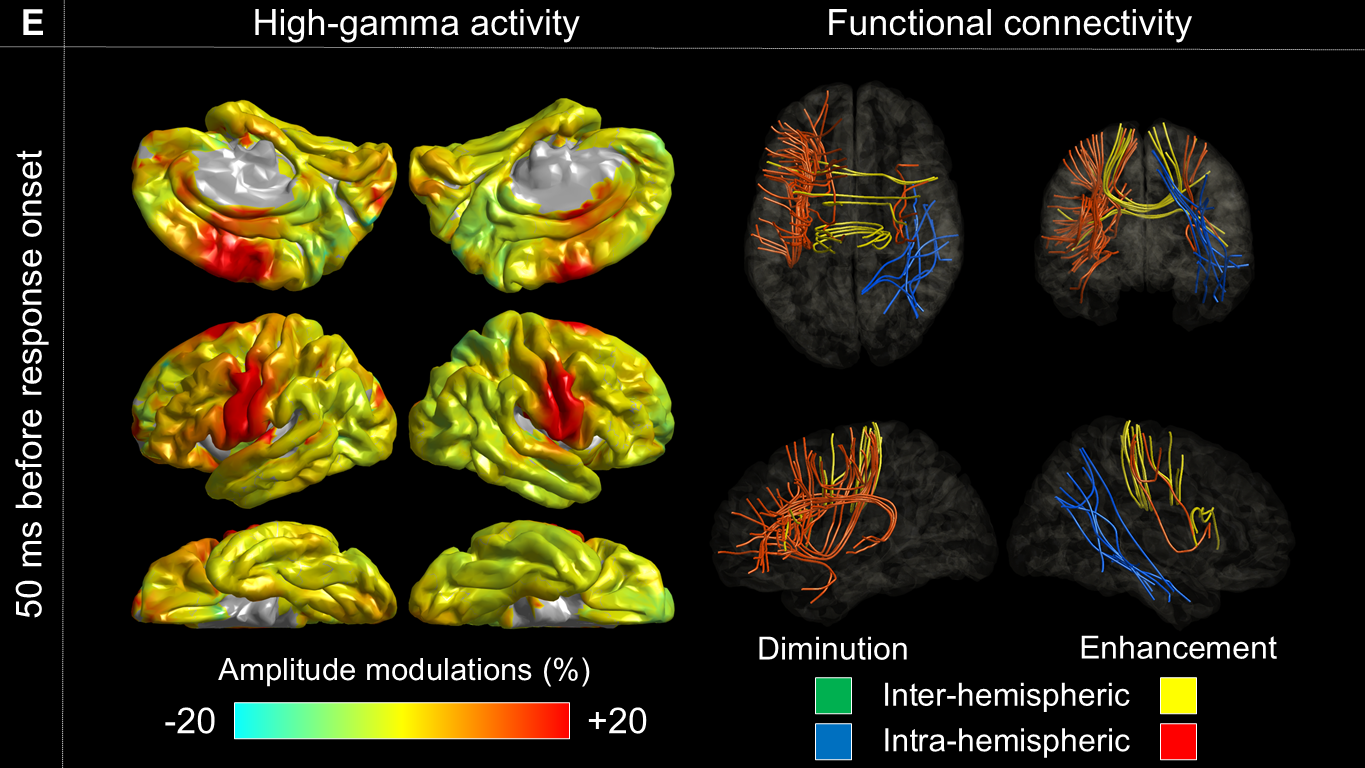

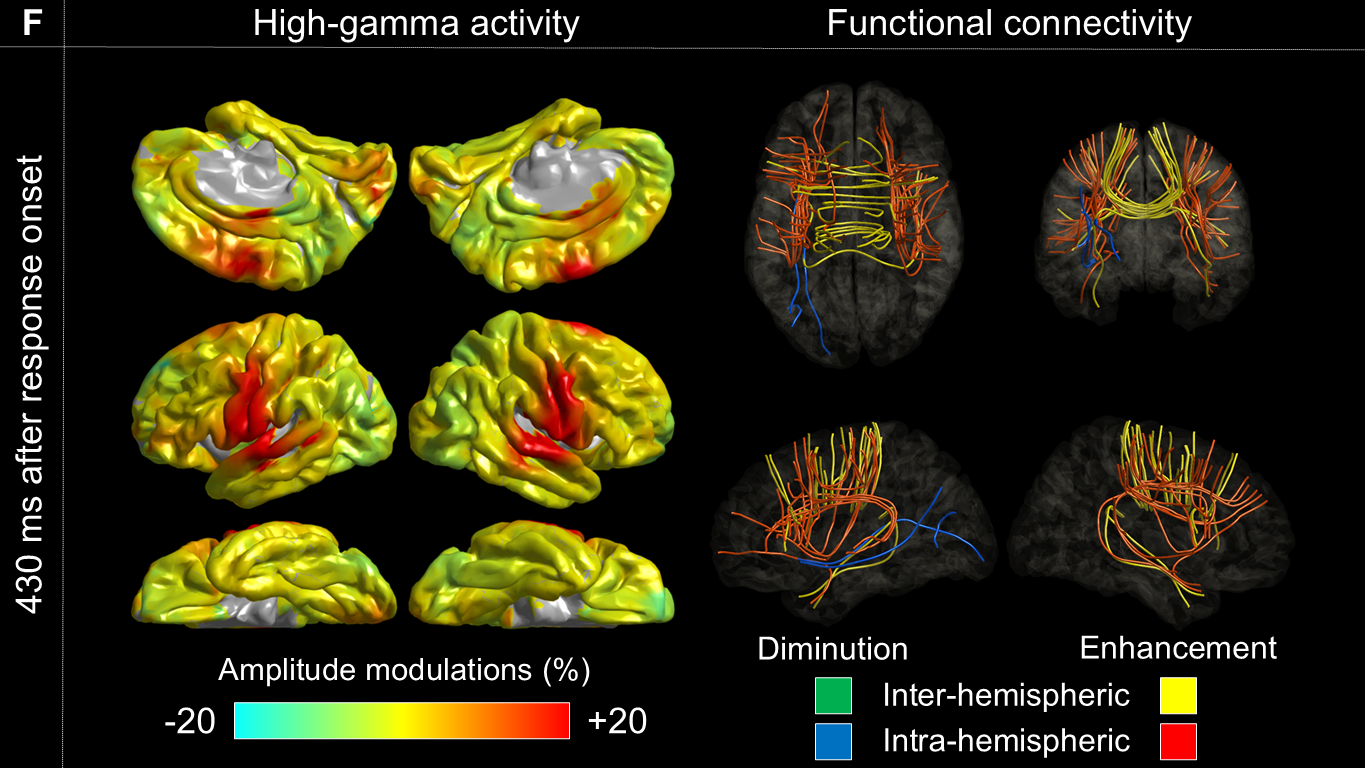

**Supplementary Figure 4 Naming-related modulations of cortical high-gamma amplitude and functional connectivity.** The left-side snapshots present task-related high-gamma amplitude modulations. The right-side snapshots present task-related modulations of functional connectivity. Orange and yellow streamlines: intra-hemispheric and inter-hemispheric functional connectivity enhancement. Blue and green streamlines: intra-hemispheric and inter-hemispheric functional connectivity diminution. (A) 220 ms after stimulus onset (association with auditory hallucination). (B) 450 ms after stimulus onset. (C) 60 ms after stimulus offset (association with receptive aphasia). (D) 380 ms before response onset (association with expressive aphasia). (E) 50 ms before response onset (association with speech arrest). (F) 430 ms after response onset (association with face sensorimotor symptoms). For a comprehensive overview of the network dynamics, please refer to **Supplementary Video 1**. L: left. R: right.

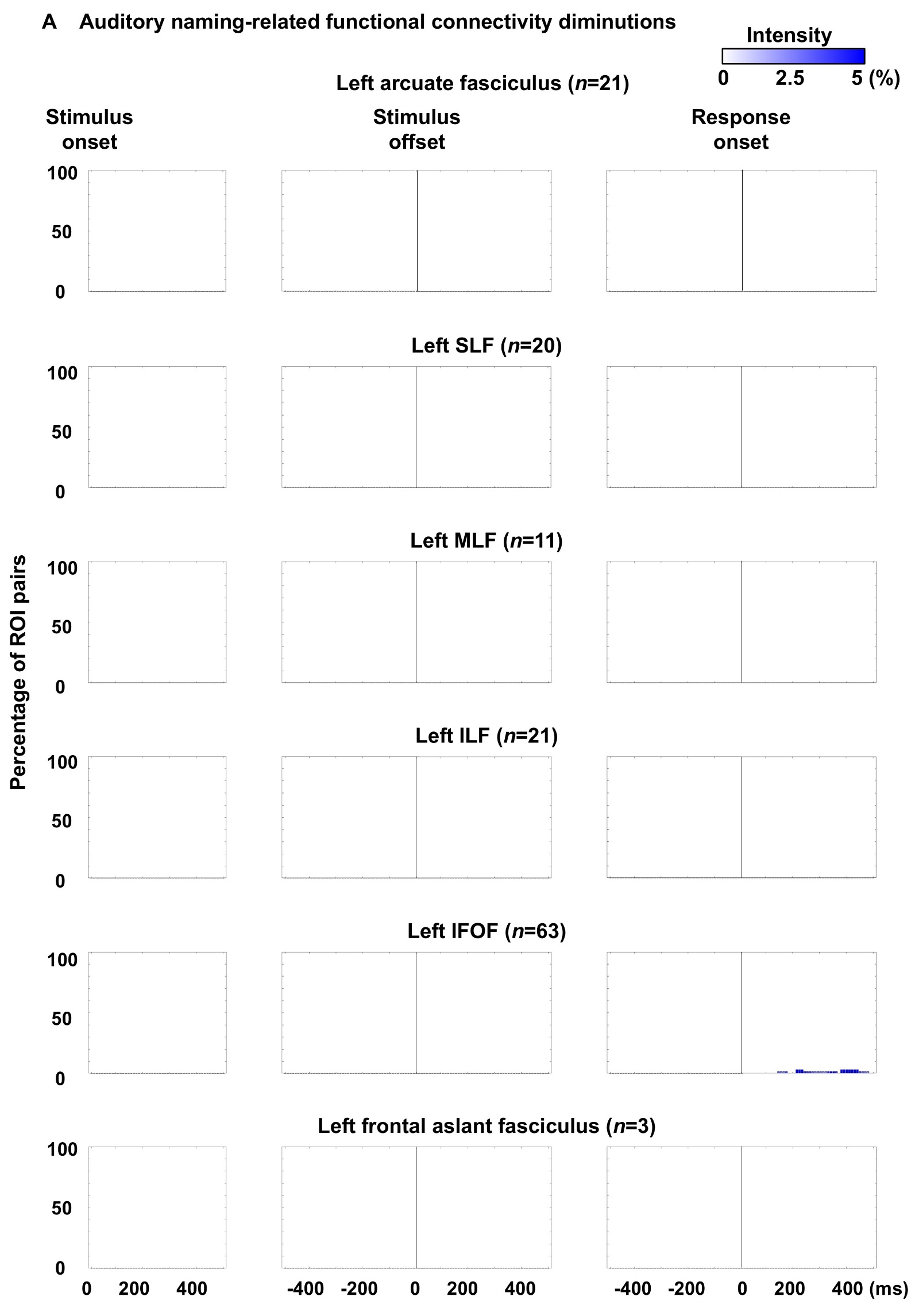

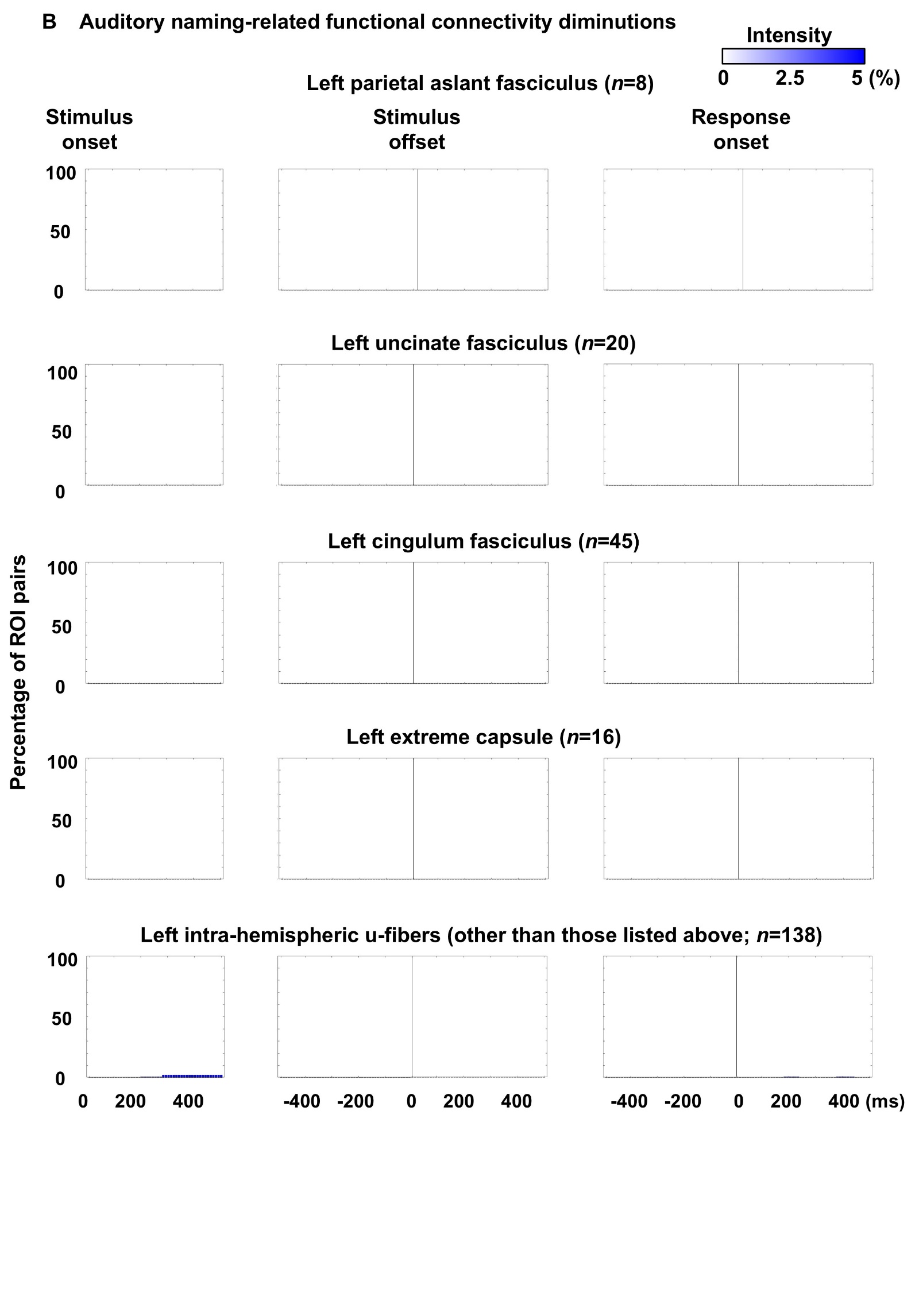

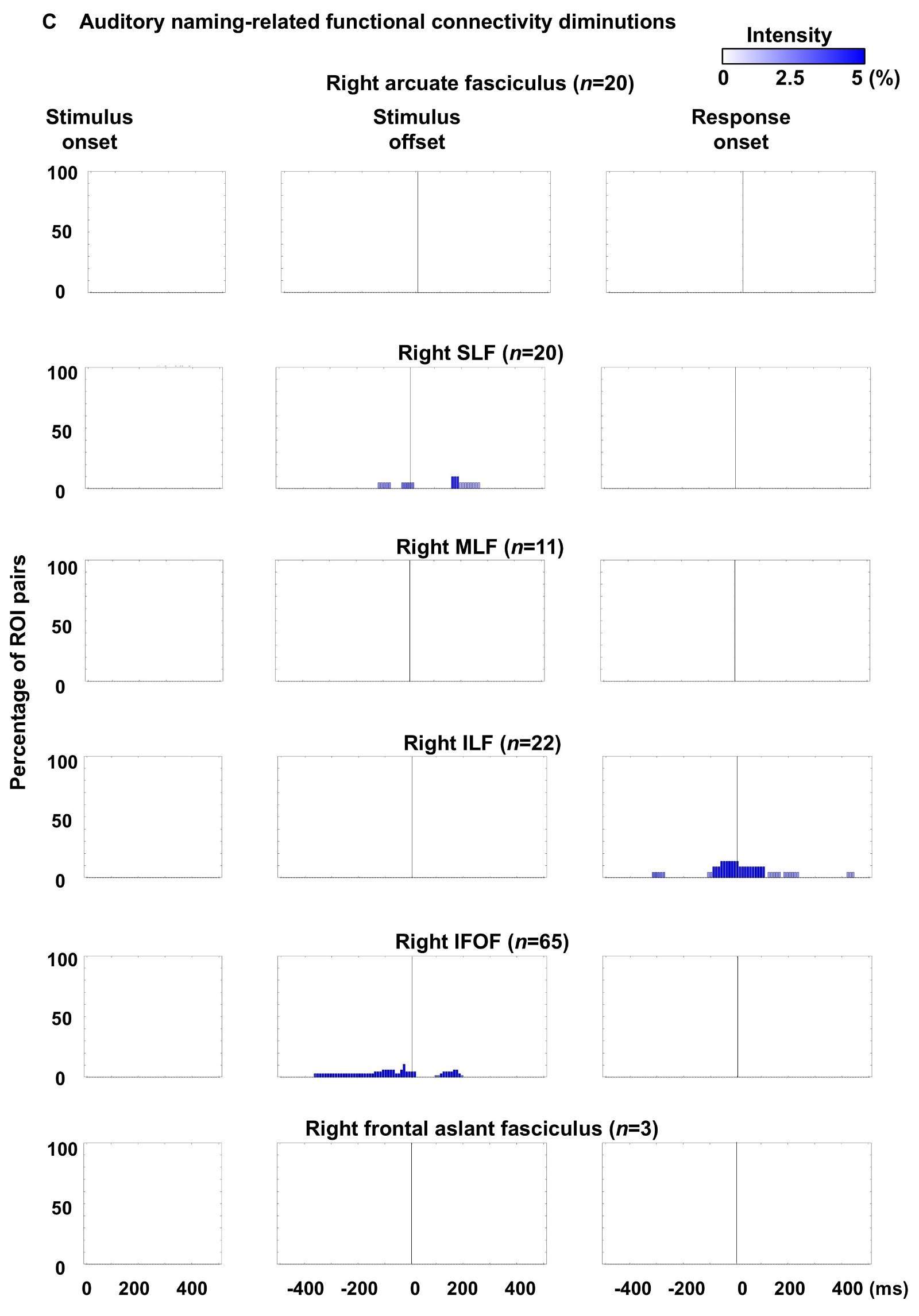

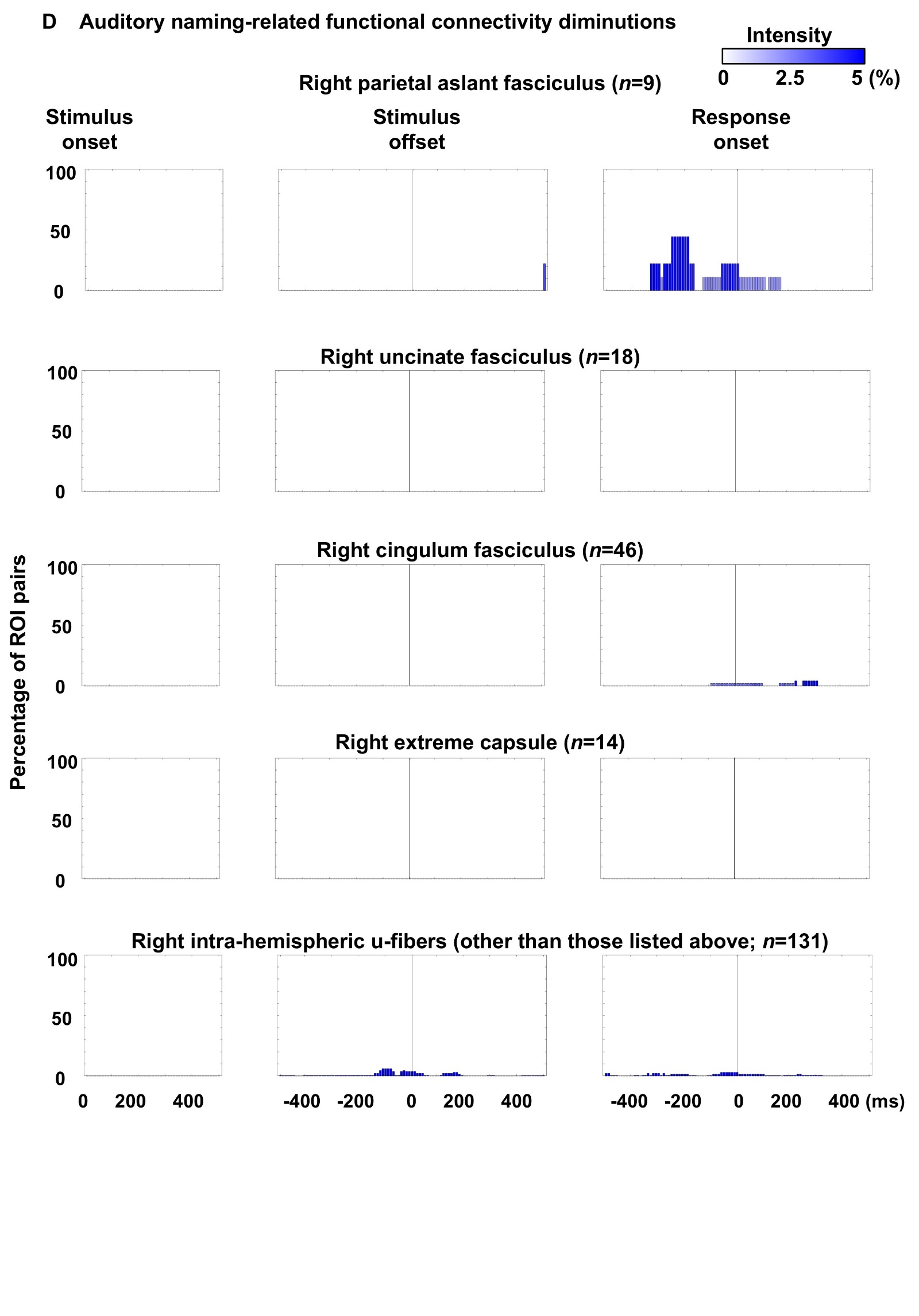

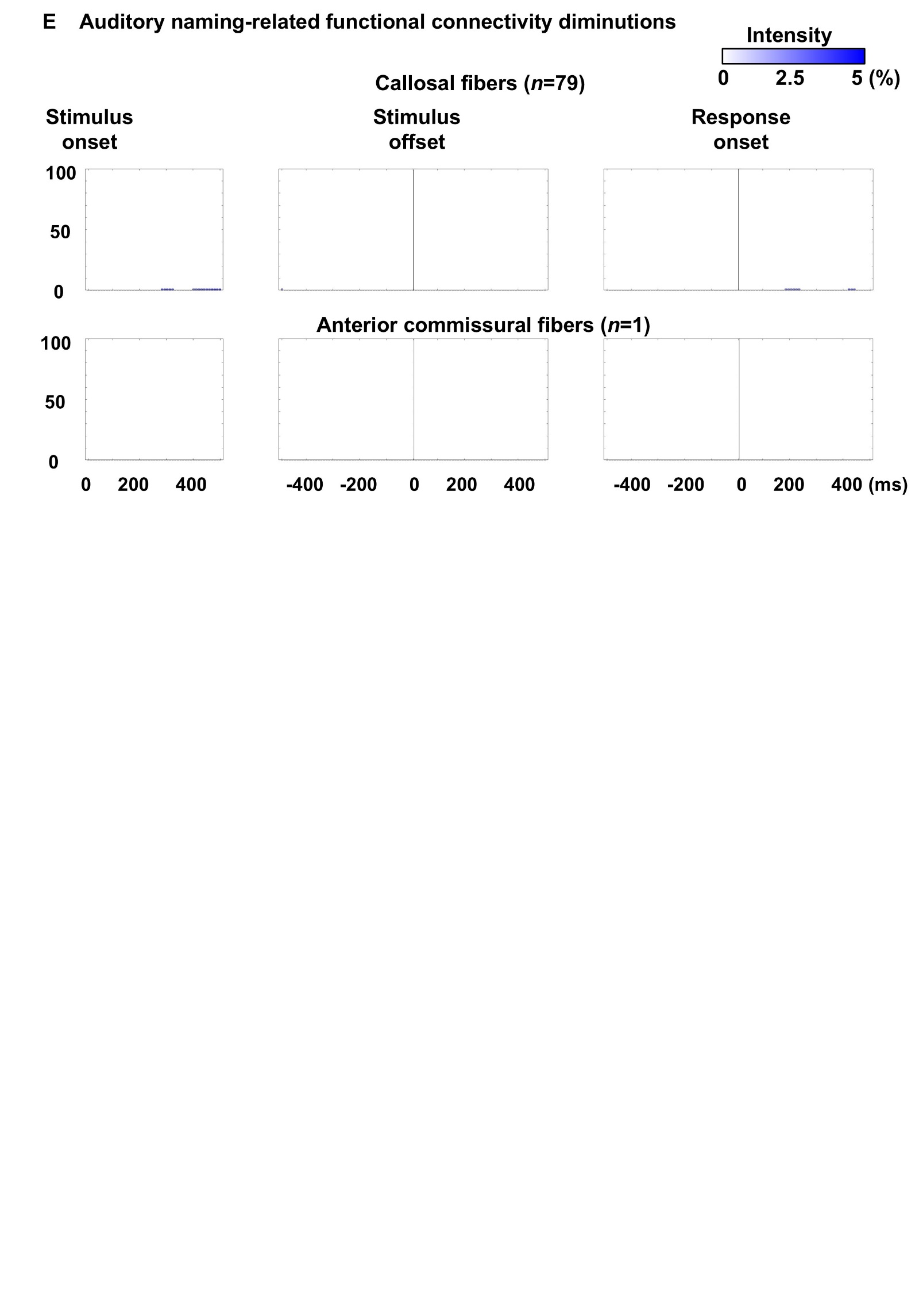

**Supplementary Figure 5 Dynamics of functional connectivity diminution through each fasciculus.** The time windows when functional connectivity was significantly diminished are indicated. (A, B) Left intra-hemispheric pathways. (C, D) Right intra-hemispheric pathways. (E) Inter-hemispheric pathways. The bar height indicates the proportion of cortical ROI pairs that showed significant functional connectivity diminution within a given time window, while the bar color represents the intensity of this diminution. **Figure 5** in the main text presents the dynamics of functional connectivity enhancement.

**Video legend**

**Supplementary Video 1 Naming-related modulations of cortical high-gamma amplitude and functional connectivity.** The left-side video images present task-related high-gamma amplitude modulations. The right-side video images present task-related modulations of functional connectivity. Orange and yellow streamlines indicate intra-hemispheric and inter-hemispheric functional connectivity enhancement, respectively. Blue and green streamlines indicate intra-hemispheric and inter-hemispheric functional connectivity diminution, respectively. L: left. R: right.

**Results from Ancillary Analyses**

Using a Mann–Whitney U test, we explored whether patients with a history of bilateral tonic–clonic seizures took more antiseizure medications immediately before iEEG recording and had a longer response time during the auditory naming task. This ancillary analysis indicated that there was no significant difference in the number of antiseizure medications between those with and without such a history (uncorrected p-value: 0.14). Likewise, there was no significant difference in response time between the two groups (uncorrected p-value: 0.47).

Using Spearman’s rank correlation test, we examined the hypothesis that older patient age is associated with a lower stimulus intensity required to induce clinical symptoms (Aungaroon et al., 2017). The observed correlations between patient age and the minimum stimulus intensity necessary to elicit a given clinical symptom did not reach statistical significance after Bonferroni correction. Given five comparisons and a familywise alpha of 0.05, the Bonferroni correction sets the per-comparison alpha at 0.01. Therefore, any uncorrected p-value must be below 0.01 to remain significant.

Spearman’s rho	Uncorrected p-value

Auditory hallucination	-0.197	0.465

Receptive aphasia	-0.521	0.022

Expressive aphasia	-0.202	0.275

Speech arrest	-0.196	0.213

Face sensorimotor symptoms	-0.213	0.051

**Supplementary reference**

Aungaroon, G., Zea Vera, A., Horn, P.S., Byars, A.W., Greiner, H.M., Tenney, J.R., Arthur, T.M., Crone, N.E., Holland, K.D., Mangano, F.T., Arya, R., 2017. After-discharges and seizures during pediatric extra-operative electrical cortical stimulation functional brain mapping: Incidence, thresholds, and determinants. Clin. Neurophysiol. 128, 2078–2086.
